# Dynamic Kinetic Models Capture Cell-Free Metabolism for Improved Butanol Production

**DOI:** 10.1101/2022.09.20.508127

**Authors:** Jacob P. Martin, Blake J. Rasor, Jonathon DeBonis, Ashty S. Karim, Michael C. Jewett, Keith E.J. Tyo, Linda J. Broadbelt

**Affiliations:** Department of Chemical and Biological Engineering, Northwestern University, Evanston, IL, USA 60208; Center for Synthetic Biology, Northwestern University, Evanston, IL, USA 60208; Chemistry of Life Processes Institute, Northwestern University, Evanston, IL, USA 60208

**Keywords:** Metabolic Modeling, Kinetic Modeling, Ensemble Modeling, ODE Models, Dynamic Models, Cell-Free Systems, Butanol Production, Parameter Optimization, Parameter Estimation, Metabolic Control Analysis, Systems Biology, Computational Biology

## Abstract

Cell-free systems are useful tools for prototyping metabolic pathways and optimizing the production of various bioproducts. Mechanistically-based kinetic models are uniquely suited to analyze dynamic experimental data collected from cell-free systems and provide vital qualitative insight. However, to date, dynamic kinetic models have not been applied with rigorous biological constraints or trained on adequate experimental data to the degree that they would give high confidence in predictions and broadly demonstrate the potential for widespread use of such kinetic models. In this work, we construct a large-scale dynamic model of cell-free metabolism with the goal of understanding and optimizing butanol production in a cell-free system. Using a novel combination of parameterization methods, the resultant model captures experimental metabolite measurements across two experimental conditions for nine metabolites at timepoints between 0 and 24 hours. We present analysis of the model predictions, provide recommendations for butanol optimization, and identify the aldehyde/alcohol dehydrogenase as the primary bottleneck in butanol production. Sensitivity analysis further reveals the extent to which various parameters are constrained, and our approach for probing valid parameter ranges can be applied to other modeling efforts.

## 1. INTRODUCTION

Widespread commercial success of metabolic engineering projects – from the production of biofuels (Atsumi et al., 2008; Fackler et al., 2021; Liew et al., 2022) to pharmaceuticals (Galanie et al., 2015; Ro et al., 2006) to polymer precursors (Arvay et al., 2021; Chung et al., 2015) – remains a challenge due to sub-optimal titers and yields (Biggs et al., 2021) and difficulty in optimizing biosynthetic pathways for production (Naseri & Koffas, 2020). This challenge is due to several factors, including the competition in living cells between non-native biosynthesis and cell viability (Dudley et al., 2015; Rasor et al., 2021; Wu et al., 2016), the highly connected nature of metabolic networks which obscures simple design choices for optimization (Tsiantis & Banga, 2020), and the difficulty of obtaining high-quality, kinetic data from living cells (Costa et al., 2016; Karim & Jewett, 2016). Cell-free systems addresses some of these issues, as cell viability no longer interferes with pathway flux, and time-course metabolomics are readily available without the need to extract intracellular species (Horvath et al., 2020; Karim et al., 2020; Miguez et al., 2019, 2021; Silverman et al., 2020; Vogeli et al., 2022). However, unlike purified systems, cell extract-based cell-free systems maintain much of the complexity of living metabolic networks (Bowie et al., 2020), which allows for native energy and cofactor regeneration to fuel biosynthesis (Jewett et al., 2008) but also preserves the complexity arising from network connectivity, making optimization nontrivial. While this complexity might seem problematic, it provides a means of collecting large data sets to understand and unravel metabolic complexity (Bowie et al., 2020; Bujara et al., 2011; Dudley et al., 2015, 2020).

Various computational models have been used to explain and analyze complex metabolic phenomena that would be difficult if not impossible to understand by manual inspection alone. For example, constraint-based models (CBMs), such as flux balance analysis (FBA)(Orth, Thiele, et al., 2010) and its many extensions (Jenior et al., 2020; Lewis et al., 2010; Sánchez et al., 2017), have been used to simulate and explain cellular behaviors. However, CBMs can only simulate steady-state solutions with limited exceptions (Mahadevan et al., 2002), must assume cell-growth or other reasonable cellular objectives for optimization, and do not explicitly represent metabolites or enzyme abundances necessary to capture effects of enzyme saturation and allosteric regulation.

Kinetic models, which use a system of ordinary differential equations (ODEs) where metabolites are tracked through time by explicitly formulated reaction rates, can overcome these limitations (Strutz et al., 2019; Suthers et al., 2021). The primary challenge in creating kinetic models is finding parameter values for each reaction rate, as there are many more model parameters than experimental data points and parameters are often unobservable, making the calculation a single “correct” or “best” set of parameters impossible. To address this problem, an ensemble modeling (EM) approach ca be applied to kinetic models, where several independently parameterized models, or parameter sets, are used simultaneously as an ensemble of models (Tran et al., 2008) to make predictions from the available data without overly relying on any single set of parameters.

Many kinetic modeling formalisms exist which highly depend on the type of system being modeled and the type and amount of data (Saa & Nielsen, 2017). For example, when modeling living-cell metabolism at steady-state, kinetic models have used “top-down” approaches for parameterization. First, known steady-state fluxes – either experimentally measured (Tran et al., 2008) or predicted from tools like flux balance analysis (J. Greene et al., 2019) – can be used as a “reference state” to lower the degrees of freedom of parameterization, though this tends to produce solutions which are more accurate nearer the reference state (J. L. Greene et al., 2017; St John et al., 2019). Second, steady-state models can forgo simulations of absolute metabolite concentrations in favor of tracking “relative” concentrations, again lowering the degrees of freedom (Tran et al., 2008) (**Supplementary Methods S1**). Many methods are then available to estimate the remaining parameters in steady-state models, including Monte Carlo methods using random sampling (Tran et al., 2008), Bayesian estimation (Saa & Nielsen, 2016; St John et al., 2019), local optimization (Gopalakrishnan et al., 2020), and global parameter optimization (Khodayari et al., 2014).

In contrast, dynamic kinetic models, which can capture transient metabolism found in non-steady state systems, typically require a “bottom-up” approach in which model parameters are initially set at experimentally measured values if available (or at reasonable estimates) and later optimized (Kim et al., 2018). These models have been used for smaller-scale systems biology such as studying signaling or small isolated biosynthetic pathways, where there are fewer model parameters and the primary hurdle is network inference as opposed to parameter estimation (Min Lee et al., 2008). These pathway-level models have frequently chosen to not explicitly model cofactors (Jia et al., 2012; van Eunen et al., 2012), a choice which simplifies parameterization but limits the scope of utility. Conversely, some dynamic metabolic models have included significant portions metabolism but have “lumped” many reactions together into a single term or reduced model structure (Buffing et al., 2018; Kurata & Sugimoto, 2018). While these strategies make dynamic metabolic models more accessible, they limit the ability of the model to fit more complex metabolic interactions, such as those found in unpurified cell-free systems, and so more general methods were needed for this work. Recently, a large-scale, high-resolution, and fully dynamic kinetic model of metabolism in an *Escherichia coli-*derived cell-free system was developed using a literature-based ensemble with Markov Chain Monte Carlo parameter optimization to study metabolic impacts on cell-free protein synthesis (Horvath et al., 2020). While this work fit data from a single experiment, this effort nonetheless demonstrated the feasibility of large-scale, bottom-up parameterization of dynamic models in biological systems.

In this work, we aimed to develop a dynamic kinetic model of *E. coli*-based cell-free metabolism for the study of biosynthetic capacity that can capture complex cell-free metabolic behavior across multiple experimental conditions. To do this, we chose to model the heterologous butanol production pathway, comprising five enzymatic steps to catalyze the transformations from acetyl-CoA to butanol (Atsumi et al., 2008), which has been successfully implemented in extract-based cell-free systems (Karim et al., 2019, 2020; Karim & Jewett, 2016). We chose this pathway in part because butanol is highly reduced and requires several cofactors, and so was expected to be highly connected with the larger cell-free metabolic network. Our model contains a large-scale representation of core *E. coli* metabolism and the heterologous butanol pathway, and model reactions are formulated with rate laws derived from mass-action kinetics of elementary enzyme mechanisms. To parameterize these models, we combined a Monte Carlo-based ensemble screening approach with a second step of local parameter optimization, which were used in series to successfully fit measurements of butanol and several species from various pathways of metabolism. Notably, both steps incorporated thermodynamic and literature constraints that have not previously been used in the bottom-up parameterization of large-scale dynamic models. The tools of metabolic control analysis (MCA) were then used to give insight into the driving factors of cell-free metabolism, as well as to give predictions regarding the optimization of butanol in this cell-free system. We anticipate our model along with the cell-free platform will facilitate both high-throughput prototyping of biosynthetic pathways for cellular design and cell-free biomanufacturing.

## 2. MATERIALS AND METHODS

### 2.1. CELL-FREE METABOLIC EXPERIMENTS AND DATA

Metabolically active cell extracts were prepared according to (Grubbe et al., 2020) from BL21-Star(DE3) *E. coli*. First, during cell-free protein synthesis (CFPS), each enzyme in the butanol pathway was separately expressed in cell-free extracts and quantified via ^14^C-leucine incorporation (Rasor et al., 2022). Next, cell-free metabolic engineering (CFME) experiments were conducted as described in (Karim et al., 2020; Rasor et al., 2022), wherein each separately expressed butanol enzyme was added to fresh cell-free extract, along with 120 mM glucose and several other cofactors, salts, and other compounds. For the concentrations of each compound in the experiment present from CFPS or added at the beginning of the CFME experiment, see **Supplementary Table S7**.

Metabolite concentrations were measured by either high-performance liquid chromatography (flowing 5 mM sulfuric acid at 0.6 mL/min on an Aminex Rezex™ ROA-Organic Acid H+ (8%) Column) or gas chromatography – mass spectrometry (using N,O-Bis(trimethylsilyl)trifluoroacetamide derivatization and an Agilent HP-5MS column with helium carrier gas). These measurements were taken every half hour from 0 hours (defined when glucose is first added) through 7.5 hours and triplicate measurements were taken at 8 hours and 24 hours. Two experimental conditions were tested: a butanol-positive (production) condition where all enzymes in the butanol pathway were present, and a butanol-negative condition where the second enzyme in the butanol pathway, 3-hydroxybutryryl-CoA dehydrogenase (HBD), was not added, thus preventing butanol production. All metabolite measurements are given in **Supplementary Table S11.**

### 2.2. NETWORK CONSTRUCTION

The metabolic network used to represent cell-free *E. coli* metabolism was adapted from the model presented by Greene and coworkers (J. L. Greene et al., 2017), which itself contained all metabolites and reactions in the BiGG *E. coli* core model (Norsigian et al., 2020; Orth, Palsson, et al., 2010), as well as reactions described in a previous model of *E. coli* metabolism (Khodayari et al., 2014). To simulate the non-steady state cell-free environment, our model removed any reactions allowing the transport of species between compartments as well as exchange reactions allowing species to enter or leave the system. Duplicate reactions from isozymes were also removed due to the lack of flux or literature data needed to distinguish each enzyme. All enzyme-inhibitor pairs described in recent kinetic models of *E..coli* metabolism were used (J. L. Greene et al., 2017; Horvath et al., 2020; Khodayari et al., 2014), as well as *E. coli* enzyme inhibition described in BRENDA (Chang et al., 2021) or MetaCyc (Caspi et al., 2014). All model regulation was either competitive or uncompetitive inhibition as reported in the literature, and noncompetitive inhibition and allosteric inhibition and activation were not considered due to lack of literature data. A full list of reactions and inhibition is given in **Supplementary Table S6**. Lastly, the reactions for the engineered pathway from acetyl-CoA to butanol (Atsumi et al., 2008) were included. The final model contained 63 enzymatic reactions, 63 metabolites, and 100 inhibitor-enzyme pairs, and is shown in **Figure 1**, as well as a more detailed representation in **Supplementary Fig. S1**.

**Figure 1:**
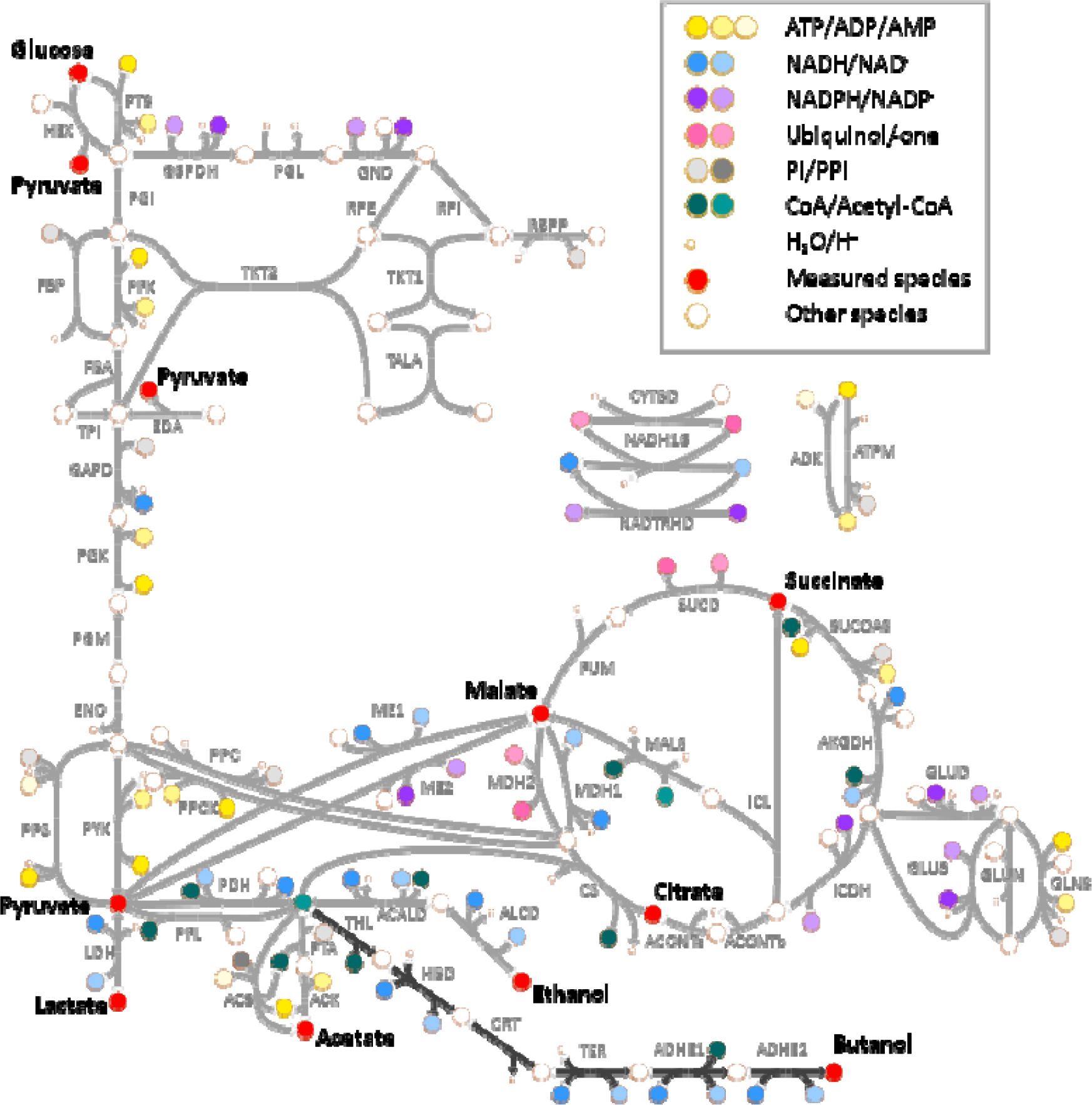
Metabolic Network Map. The utilized network includes *E. coli* core metabolism (gray arrows), as well as the 6-step butanol pathway (black arrows at the bottom of the map). Measured metabolites are labeled in black text and shown as red circles. Cofactors are shown as colored circles according to the legend, and water and protons (not kinetically included in the model) are shown as smaller circles. All other species are shown as unfilled circles.

To construct a kinetic model, an approximate rate law was derived and applied to each reaction. This equation to describe the rate of each enzymatic reaction expands the random-order ternary complex enzyme mechanism and rate law described in (Cornish-Bowden, 1979) with the additional assumption that the dissociation constant of each substrate in multi-substrate reactions is unaffected by binding order. This assumption halves the number of parameters per substrate compared to an order-dependent rate form (Cornish-Bowden, 1979). The resulting rate law is mechanistically grounded and is able to use experimental Michaelis constants as estimates for dissociation constants, yet does not require manual curation to specific enzyme mechanisms, is expandable to an arbitrary number of substrates, products, and inhibitors, and reduces the hurdle of parameter estimation. In the case of two substrates and two products:

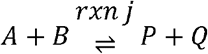

The resultant net rate equation for reaction 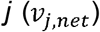, or the rates of the forward and reverse elementary steps (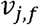 and 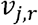), is given below:

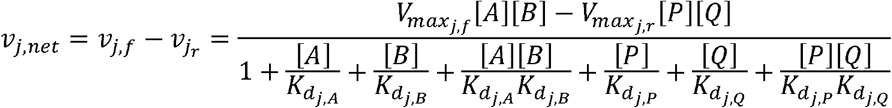

The parameters 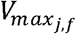 and 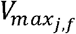 represent the forward and reverse maximal rates, or rate constants, and 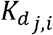 represents the dissociation constant for species *i*. For details, see **Supplementary Methods S2**.

A few unique metabolites and reactions were implemented for this model. First, because the concentration of water is assumed to be constant, no dissociation constants for water are used in any rate law, and instead is implicitly included in the rate constant. Hydrogen ions are assumed to be buffered and are similarly not included in any rate laws. Gas exchange for carbon dioxide and oxygen between the aqueous and gas phases was included in the model, as CFME experiments were performed in a small volume of liquid with a large headspace. Preliminary partial differential equation (PDEs) modeling predicted spatial gradients of aqueous gas concentrations were negligible and so were not further considered. Gas exchange at the surface of the liquid was assumed to be quasi-equilibrated and was modeled kinetically using Henry’s law constants as constraints. See **Supplementary Methods S11** for more details.

### 2.3. MODEL PARAMETERIZATION

Due to the difficulty of parameter estimation for large-scale kinetic models and the model equations being inherently underdetermined by the data, this work employed an ensemble modeling approach in which many models, known as an “ensemble” of models, each with the same structure but with independent parameter sets, are used simultaneously. This approach not only alleviates the difficulty of finding a single set of parameters that accurately describes the experimental data, but also acknowledges the inability to determine a single optimal parameter value, known as parameter identifiability or observability, that is a hallmark of large-scale kinetic models and provides a metric by which uncertainty in both model parameter values and model behavior can be quantified.

Initial values, or parameter priors, for model dissociation constants were set to experimentally measured Michaelis constants from BRENDA (Chang et al., 2021) and MetaCyc (Caspi et al., 2014) when available. Of the 319 dissociation constants in the model, 116 of these enzyme-substrate pairs had at least one Michaelis constant available in these databases. When multiple experimental values were available, the geometric mean was used. Where literature values were not available, the corresponding dissociation constant was set to 0.1 mM, which was the median value of all measured Michaelis constants (see **Supplementary Table S9 and S10** for all initially available literature values). Of the 100 inhibition constants in the model, 13 values were available on either BRENDA or MetaCyc, and the remainder were initially set to 0.5 mM, which was the median value of the available constants.

Because turnover numbers are typically reported as specific quantities relative to enzyme levels, the use of literature values for rate constants would require accurate and broad proteomics for the cell-free experimental system, which were not available. Instead, rate constants in a previously published model of cell-free protein synthesis from Horvath, *et al.* (Horvath et al., 2020), which contained many of the same reactions in central *E. coli* metabolism, were used as parameter priors. For reactions used in this work that were not in the Horvath model, the initial parameter prior was set to the geometric mean of all rate constants in the Horvath model. The direct use of the Horvath rate constants in our model gave very poor fit to our data, as the data fit by the Horvath model qualitatively differed from this study due to various experimental differences. Nonetheless, the Horvath parameter values provided an initial point from which parameters in this study could be further tuned.

Once priors for all parameters were obtained (**Figure 2a**), a single parameter set was generated by sampling all model parameters within one order of magnitude in each direction of each prior. Within each reaction, these parameters were further constrained by the Haldane relationship, which provided physiological feasibility by ensuring consistency with known thermodynamics. This equation related the equilibrium constant in reaction *j* (*K_eq,j_*) with the forward and reverse rate constants (*V_max,f_*, *V_max,r_*) and the dissociation constants for each substrate or product *i* in reaction 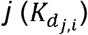 along with their stoichiometry (*S_i,j_*) and is given below in Eq. 1 (see **Supplementary Methods S3** for details). Equilibrium constants were calculated using eQuilibrator 3.0 (Beber et al., 2022).

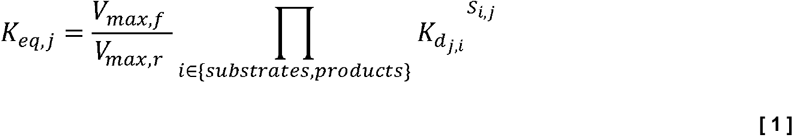

**Figure 2:**
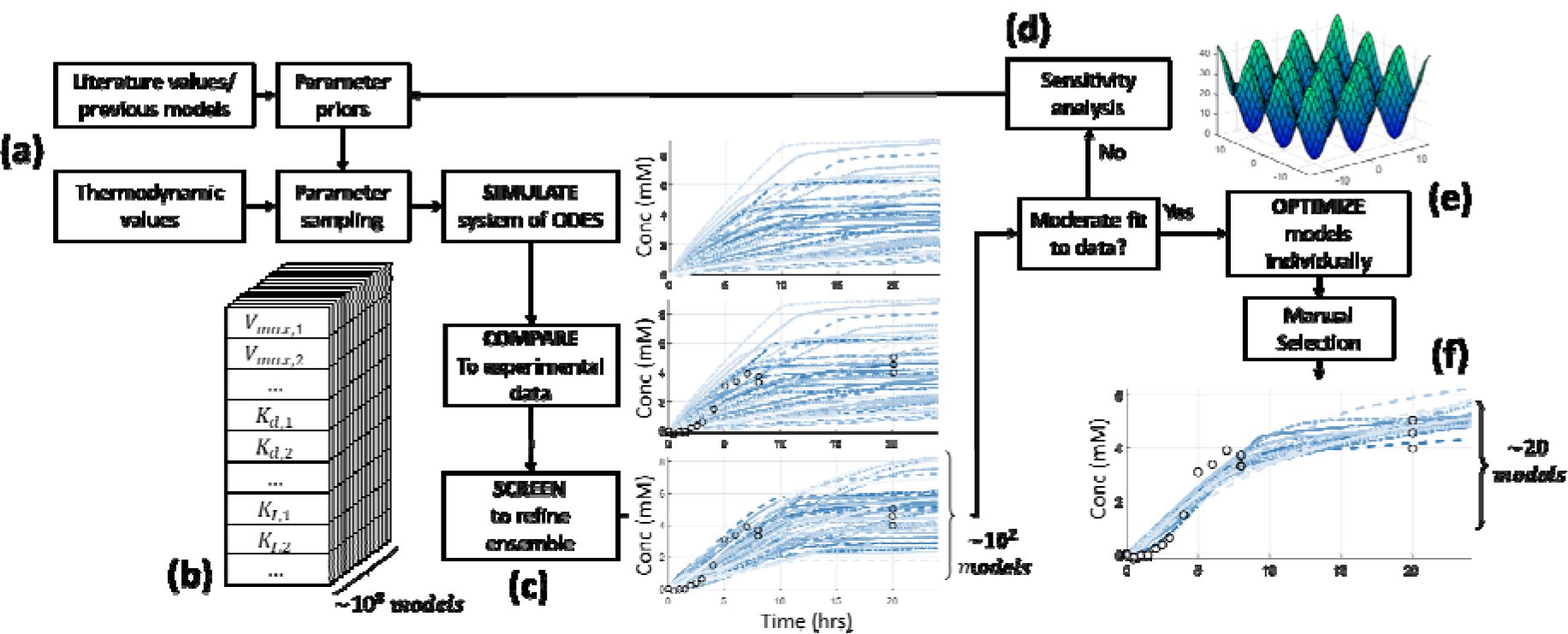
Model parameterization pipeline. (a) Parameter priors are initially constructed from literature values and similar previous models. (b) Individual models, or parameter sets, are sampled from parameter priors while simultaneously applying thermodynamic values as constraints. This is repeated to create ensembles of 10^8^ models. (c) The initial ensemble is simulated via ODEs and then compared to experimental data; the ensemble is then screened to the best ~10^2^ models based on agreement to this data. (d) When the agreement to experimental data after screening was still insufficient, parameter sensitivity analysis was performed and used to update parameter priors for additional rounds of ensemble sampling and screening. (e) Once screened models showed adequate agreement to experimental data, each individual model was further refined via local parameter optimization. (f) This full pipeline generated an ensemble of 20 models which gave excellent fit to the experimental data and were further used for analysis and predictions.

This procedure was repeated to generate an ensemble of 10^7^ independent parameter sets (**Figure 2b**). Each parameter set was simulated by integrating the system of ordinary differential equations (ODEs) which describe the change of each metabolite concentration over time (**Figure 2c**) and were solved in MATLAB 2020b (The MathWorks, Natick, MA) (**Supplementary Methods S7**). Initial ODE metabolite concentrations were set, in order of priority, first by available measurements, then by experimentally added concentrations, and finally by known *E. coli* intracellular concentrations as measured by (Bennett et al., 2009), corrected for dilution. Any metabolite not in any of these categories was set to 0 mM. Initial concentrations of gas species were set to atmospheric conditions, and aqueous gases were set to equilibrium concentrations via Henry’s Law. See **Supplementary Table S7** for values for each model metabolite.

The performance of each parameter set was quantified by the weighted root mean squared errors (RMSE) of the simulated metabolite profiles to experimental metabolite measurements across both experimental conditions. The RMSE values for succinate were weighted 6-fold compared to other metabolites due to poor fit to succinate measurements in early models, and the RMSE for butanol were weighted 20-fold because butanol production was emphasized in this model (**Supplementary Methods S8**). The ensemble was “screened” by keeping only the 100 parameter sets with the lowest weighted RMSE, which were used for further optimization and analysis.

Because models generated from the initial parameter priors failed to adequately fit the experimental data, sensitivity analysis was performed to adjust parameter priors. The finite differences method was used to calculate the sensitivity of model fitness with respect to each parameter value (**Supplementary Methods S6**) and was applied to all models in the final screened ensemble. The priors for any parameters that were highly sensitive across most models in the ensemble were then adjusted in the direction of improved fitness to the data (**Figure 2d**).

After iterating this procedure to update parameter priors, the fit of all models to the data remained poor. To remedy this, we applied a local parameter optimization to each of the top 200 models in the screened ensemble (**Figure 2e**). In this procedure, each screened model was used as the initial point in the MATLAB pattern search algorithm, which attempts to find a local optimum fitness value by using heuristics to traverse the parameter space. This algorithm showed better performance than both particle swarm and genetic algorithm global search methods as well as simpler local search methods, such as MATLAB’s “fmincon”. Individual parameter constraints were applied to ensure rate constants, dissociation constants, and inhibition constants stayed in physiologically realistic ranges, and parameters were log-transformed to allow the Haldane relationship to be applied as a set of linear inequality constraints within the optimization algorithm (see **Supplementary Methods S4** for details on optimization constraints).

Local parameter optimization was carried out on MATLAB 2020b on the Northwestern Quest High Performance Computing Cluster. Each of the 200 optimization runs employed parallelization within the “pattern search” toolbox and ran on 250 CPUs with a wall time of 4 hours. After optimization, each parameter set was re-ranked according to the weighted RMSE fitness of the optimized point. The top 20 optimized models were chosen for the final ensemble and further analysis, as these models showed the strongest agreement to the experimental timecourse metabolite measurements in addition to capturing a distinct shift in acetate and succinate production in the butanol-negative condition.

### 2.4. METABOLIC CONTROL ANALYSIS

Metabolic control analysis (MCA) is a means of measuring how network features, such as metabolite concentrations, reaction fluxes, and model parameters, impact each other. This analysis can be either local, such as concentration elasticities 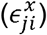, which describe the change in reaction flux caused by those concentrations or parameters within that reaction, or they can be global, such as flux control coefficients 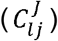, concentration control coefficients 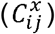, and the Jacobian (*J_ii′_*), which describe how changes in each concentration or flux affect distant network features through shared metabolites. The definitions of each normalized value, in terms of metabolite concentrations *x_i_* and *x_i′_*, and reaction fluxes 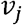 and 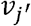, as given in Equations 2–5 below:

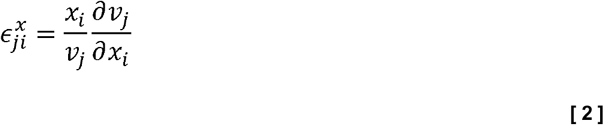

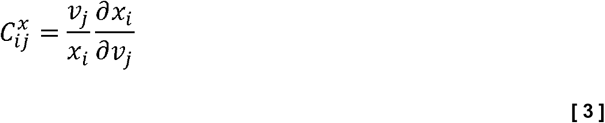

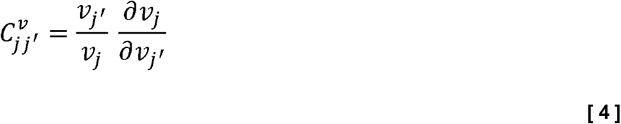

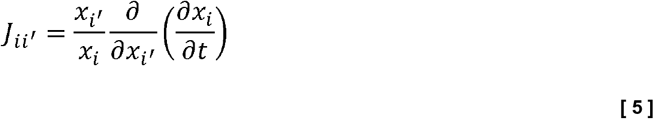

The above MCA features were calculated analytically, as described in (Hofmeyr, 2001) and detailed in **Supplementary Methods S5**. Because the system in this model is dynamic, these values were calculated at discrete timepoints for every 0.5 hours between 0 and 24 hours in the model simulation. In all analyses presented in this work, only values calculated between 0.5 hours and 7.5 hours are presented, since values calculated at 0 hours caused numerical instability from initial concentrations of zero, and values after 7.5 hours showed wide variation due to the lack of experimental data between 8 and 20 hours to constrain model behavior.

## 3. RESULTS

### 3.1. DYNAMIC MODEL FITTING AND GOODNESS OF FIT

To train our dynamic model of cell-free metabolism, we iterated the process of sampling around parameter priors to generate an ensemble, screening the ensemble against the experimental data, and updating the parameter priors using parameter sensitivity analysis fifteen times until the model adequately fit the metabolite data (see **Supplementary Table S9** for each iteration of parameter priors). From this iteration, the top 200 screened parameter sets (of 10^8^ sampled) showed significantly improved fit to the nine measured metabolites in the butanol-positive condition, with a 32% reduction in the median weighted RMSE compared to screened parameter sets from early iterations **(Supplementary Table S1)**. However, the experimental data from the butanol-negative condition showed a large increase in acetate accumulation and a decrease in succinate accumulation as compared to the butanol-positive condition, which were not captured by any of the top 200 parameter sets **(Supplementary Fig. S3)**. These shifts in metabolism were difficult to capture in part because the measured increase in acetate could not simply be accounted for from the measured decrease in butanol. Additionally, previous experimental data showed that changes in acetate and succinate titers – as well as ethanol and lactate, to a lesser extent – were correlated with changes in butanol titers (**Supplementary Fig. S2**), which implied that system-level effects were coupling the production of all these metabolites.

We next employed automated parameter optimization algorithms to fine-tune each parameter set and found that the local optimization “pattern search” algorithm (The Mathworks, Inc.) significantly improved the weighted RMSE of nearly all models, reducing the weighted RMSE a further 46% **(Supplementary Table S1)**. We found that the top 20 parameter sets (in terms of post-optimization weighted RMSE) captured the experimentally observed shifts in acetate and succinate as well as all other experimentally measured metabolites **(Figure 3)**. Importantly, the combination of ensemble screening and local parameter optimization was found to be necessary to successfully fit both positive and negative butanol production conditions, as each method alone was not sufficient to fit the data, even when sampling around the final updated parameter priors (**Supplementary Fig. S4**).

**Figure 3:**
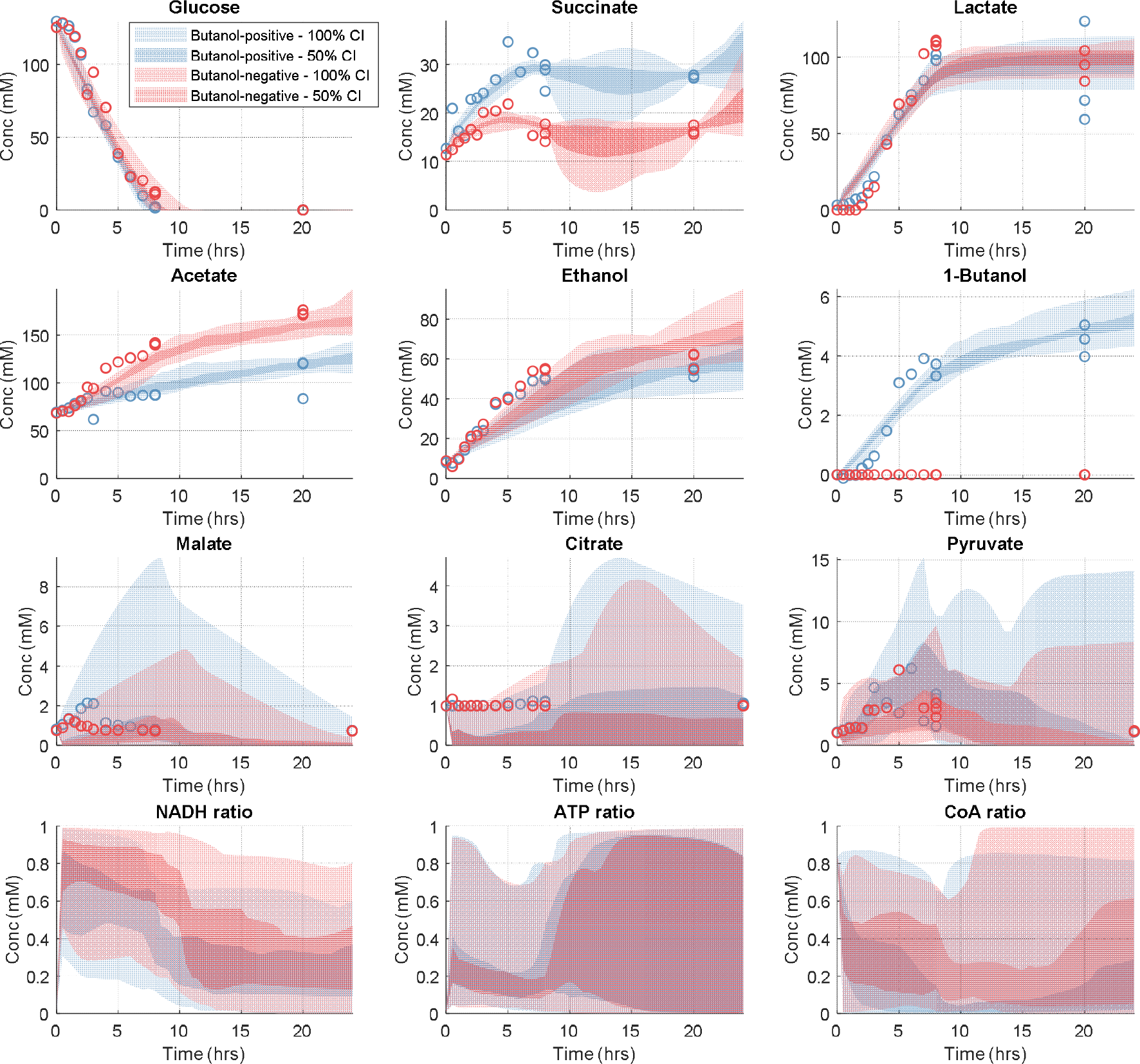
Final ensemble simulations compared to experimental data. Model simulated metabolite time courses are shown for the nine metabolites with experimental measurements, as well as predicted time courses for the ratios of NADH to all NAD species, ATP to all adenosine species, and free CoA to all CoA species. Experimental measurements are shown as circles. The simulations of the top 20 models in the ensemble are shown as confidence intervals (CI); the dark region is a 50% CI and therefore contains the middle 50% of models, and the lighter region is a 95% CI. Blue data represent the butanol-positive condition, and red data represent the butanol-negative condition.

### 3.2. ANALYSIS OF CELL-FREE METABOLISM AND THE BUTANOL PATHWAY

With a trained ensemble of 20 parameter sets, we wanted to understand how simulated metabolism was able to fit experimental metabolite measurements. The most interesting behaviors of this data were the shifts in acetate and succinate concentrations in the butanol-negative condition compared to the butanol-positive condition. Our models suggest that acetate accumulation in the butanol-negative condition is due to increased acetyl-CoA concentration (**Supplementary Fig. S5**), which itself is caused by the lack of acetyl-CoA flux into the butanol pathway; these conclusions are supported by the high control of acetyl-CoA concentration by butanol concentration in MCA, as well as pathway flux analysis (**Supplementary Fig. S6**; **Supplementary Methods S9**). Next, by decomposing the reactions for succinate production into “elementary” (forward and reverse) fluxes, we observed that the decrease in succinate production in the butanol-negative condition was not from lowered “forward” reaction fluxes but instead from increased “reverse” flux of succinyl-CoA synthase (in the direction of succinyl-CoA from succinate) (**Supplementary Fig. S7a**). This reverse flux – equivalent to product inhibition – was attributable to an increase in free Coenzyme A (CoA) (**Supplementary Fig. S7b**) and was confirmed with MCA showing strong negative control over this reaction by CoA (**Supplementary Fig. S7c**). The increase in CoA, in turn, is caused by the increase in phosphotransacetylase (PTA) flux in the acetate pathway from the butanol pathway knockout, which can be seen both in the high concentration control coefficient of CoA from PTA (**Supplementary Fig. S7c**) and the reactions that contribute to changes in acetyl-CoA concentration (**Supplementary Fig. S6b**).

Significantly, the increases in acetyl-CoA and CoA in the butanol-negative condition – which are responsible for the increase in acetate and decrease in succinate, respectively – are not trivial consequences of the butanol pathway knockout in the negative condition, as the butanol pathway uses acetyl-CoA and releases CoA in the same stoichiometric ratios as the acetate pathway. Instead, our model analysis found that these shifts are only possible due to a bottleneck in the butanol pathway wherein intermediates were accumulated, causing butanol pathway flux to have a net decrease in the quasi-steady state concentrations of both acetyl-CoA and CoA, as compared to acetate pathway flux (**Supplementary Fig. S8a**). This prediction of butanol pathway intermediate accumulation was conserved among all 20 parameter sets and will be discussed further in the context of predictions for pathway optimization.

Beyond acetate and succinate shifts, carbon metabolism within the model remains a highly connected, balanced network. Strong control of butanol production is maintained among all models in the final ensemble, with consistent control by acetaldehyde dehydrogenase (ACALD), NAD dehydrogenase (NADH16pp), malate dehydrogenase (MDH), and NAD-dependent malic enzyme (ME1) (**Supplementary Fig. S8b**). However, this control varies in direction (i.e., some models have parameters where this control is positive and others where the control is negative) suggesting several possible control mechanisms. Ethanol production is even more tightly controlled, which allows the model to fit experimental observations of unchanged ethanol production between conditions despite changing acetyl-CoA concentration. MCA shows that this effect is due to saturation of the first step of ethanol production (ACALD) by acetyl-CoA, as well as further distributed control over ethanol production across ACALD, glyceraldehyde-3-phosphate dehydrogenase (GAPD), fructose bisphosphatase (FBP), pyruvate dehydrogenase (PDH), and NADH16pp fluxes (**Supplementary Fig. S12**). In contrast, the production of lactate is not strongly controlled by other reactions, as can be seen in the very low flux control coefficients for lactate dehydrogenase (LDH) (**Supplementary Fig. S9a**), but is instead more strongly controlled by NADH than by pyruvate or other metabolites (**Supplementary Fig. S9b**). This finding suggesting that lactate production stops at the same time as glycolytic flux not because of lack of pyruvate, but instead due to lack of NADH regeneration in glycolysis, which thereby lowers the NADH:NAD^+^ ratio and brings LDH flux into equilibrium. Lastly, the consumption of glucose itself shows a large amount of metabolic control from NADH16pp, GAPD, TPI, FBP, PTA, and HEX fluxes (**Supplementary Fig. S10**), which are important reactions in the control of the other measured metabolites discussed above. GAPD and TPI are both interesting cases, as they are thermodynamic bottlenecks with large positive Gibbs free energies (Beber et al., 2022). While some previous literature has suggested that glycolysis in *E. coli* is predominately controlled by the rate of ATP consumed outside of glycolysis itself (Solem et al., 2003), other work has identified GAPD as a key controlling and limiting step of glycolysis, both in *E. coli* (Centeno-Leija et al., 2013; Cho et al., 2012) and in other organisms (Shestov et al., 2014). In the context of metabolism in the cell-free system described here, this finding would indicate that GAPD flux, and the NADH generated by this reaction, is the dominant factor controlling glycolytic flux.

### 3.3. Model Predictions for Optimization of Butanol Production

Kinetic models offer advantages in their ability to extrapolate and make predictions about unseen experimental conditions (J. L. Greene et al., 2017; Strutz et al., 2019). Here, we simulate several new experimental conditions (e.g., altering butanol pathway enzyme concentrations and the initial concentrations of metabolites) that aim to increase butanol production. In the cell-free reaction, the concentrations of heterologous enzymes are easily perturbed while the enzymes in core metabolism are fixed. We therefore first used our model to adjust the levels of all five butanol pathway enzymes at 0.5x, 2x, 5x, and 10x fold-changes from the baseline butanol-positive condition (**Figure 4**). Within the model, these conditions were simulated by adjusting the forward and reverse rate constant parameters (V_max_) of the corresponding reactions. When changing all enzymes simultaneously, we found that final 20-hour butanol titers were predicted to increase monotonically with increasing butanol pathway enzyme levels but all models show diminishing returns at the highest levels of enzyme overexpression, implying that heterologous pathway enzymes become no longer limiting. When increasing each butanol pathway enzyme individually, all models in the final ensemble predicted that increasing the bi-functional aldehyde-alcohol dehydrogenase, which carries out the final two reactions in the butanol pathway, ADHE1 and ADHE2, gave a much larger improvement in butanol titer than increasing any other butanol pathway enzyme (**Figure 4**).

**Figure 4:**
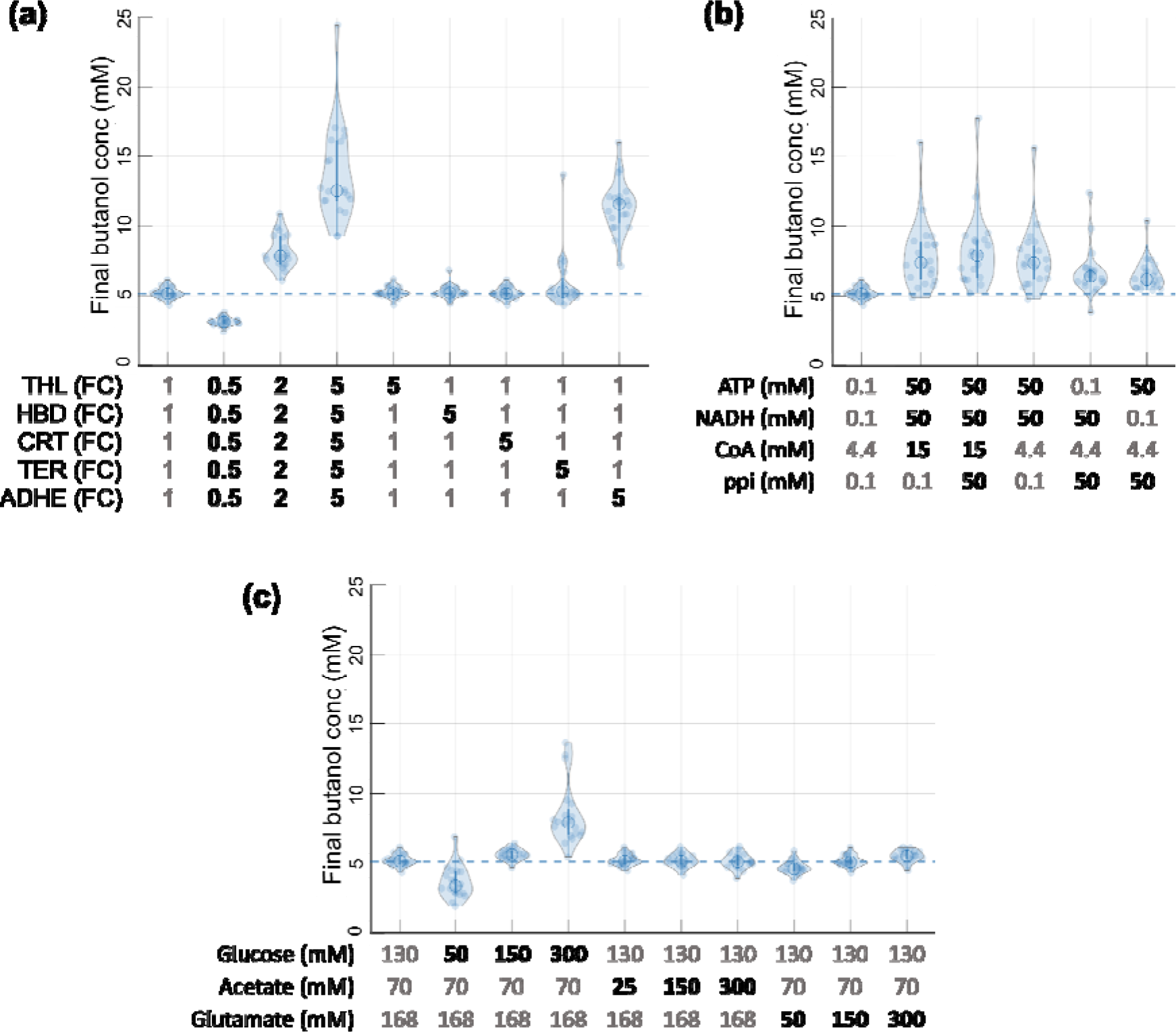
Predictions for butanol optimization. (a) Enzyme level changes were simulated to optimize 24-hour butanol titer. The left-most column shows the simulated butanol at 24 hours in the base butanol-positive condition, which was fit to experimental data. Each light blue dot represents one of the 20 models in the final ensemble, and their distribution is summarized by each violin plot with medians given by unfilled dark blue circles and interquartile ranges given by dark blue lines. The horizontal dashed line shows the base condition median prediction as a baseline. The next three columns show the predicted butanol titer from simultaneously adjusting all five enzymes in the butanol pathway by either 0.5x, 2x, or 5x their respective levels in the base production condition. The next five columns show the predicted butanol titer from adjusting each of the five enzymes one at a time by 5x their respective levels in the base production condition. (b) Initial cofactor level changes were simulated to optimize 24-hour butanol titer. The left-most column shows the simulated butanol at 24 hours in the base butanol-positive condition and the base cofactor concentrations in gray. The next five columns show the predicted butanol titer from raising the concentrations of the cofactors predicted to improve butanol titers, which included NADH, ATP, CoA, and pyrophosphate (ppi), either individually or in groups. (c) Predicted butanol titers from adjusting the initial concentrations of either glucose, acetate, or glutamate are shown. Initial concentrations of each species in the base condition (left-most column) are shown in gray.

As previously mentioned, all parameter sets predicted that a bottleneck to accumulate intermediates in the butanol pathway was needed to fit the observed shifts in acetate and succinate production. Interestingly, despite the lack of enzyme-specific training data, all parameter sets further agreed that this bottleneck would accumulate butyryl-CoA and butyraldehyde, the substrates to ADHE1 and ADHE2, respectively (**Supplementary Fig. S8a**). To understand this model behavior, we performed a thermodynamic analysis of the butanol pathway. This analysis showed that the first step of the pathway, THL (ΔG’^m^ = +23 kJ/mol), and the second step, HBD (ΔG’^m^ = −20 kJ/mol), must be kinetically fast and therefore near equilibrium to establish flux into the butanol pathway. Similarly, the third step of the pathway, CRT, is only slightly exergonic (ΔG’^m^ = −1.0 kJ/mol), and so the first three steps of the pathway are at quasi-equilibrium, and the fourth step (TER) is again quite favorable (ΔG’^m^ = −61 kJ/mol). The quasi-equilibrium of the first three reactions means that any bottleneck in these reactions or the subsequent fourth reaction would not result in accumulation of intermediates but would equilibrate back to acetyl-CoA, preventing the shifts seen in the data. Therefore, bottlenecks responsible for accumulation were only possible in the last two steps of ADHE1 and ADHE2 (ΔG’^m^ = −10 kJ/mol and ΔG’^m^ = −25 kJ/mol, respectively) once flux was effectively irreversible. For this reason, the alcohol-aldehyde dehydrogenase represents the primary enzyme bottleneck in the butanol pathway, and its overexpression is consistently predicted to have the only significant impact on increased butanol production, which is validated by previous experimental observations (Karim & Jewett, 2016). These results provide valuable knowledge that can be leveraged experimentally when expression and solubility of the enzyme are not limiting.

We next used the models to predict the effect of changing the initial cofactor and substrate concentrations. First, titrations were simulated for the initial concentration of acetyl-CoA, CoA, ATP, ADP, AMP, NADH, NAD^+^, NADPH, NADP^+^, pyruvate, phosphoenolpyruvate, inorganic phosphate, and pyrophosphate (**Supplementary Fig. S11**). While parameter sets did not have the same consensus as for enzyme level predictions, the simulated initial concentrations of 50 mM NADH (compared to 0.1 mM in the base condition of butanol-positive condition), 50 mM ATP (0.1 mM in the base condition), 15 mM CoA (4.4 mM in the base condition), and 50 mM pyrophosphate (0 mM in the base condition) all predicted increases in butanol titer to varying degrees. We then simulated butanol production with these new combinatorial conditions which predicted the highest median butanol production (**Figure 5**). These cofactors have previously been found experimentally to be the most important additions for controlling cell-free metabolism (Karim et al., 2018). For high initial NADH, butanol titer is increased both by competitively inhibiting PTA and providing reduction potential to relieve the ADHE1 and ADHE2 bottleneck. The increase in CoA increases butanol titers primarily by increasing pyruvate flux to acetyl-CoA via pyruvate dehydrogenase and indirectly increasing NADH through increased flux in AKGDH. High pyrophosphate was predicted to decrease acetyl-CoA synthase flux and minimize futile cycling. Lastly, high initial ATP was predicted to increase butanol production by inhibiting PTA and LDH flux (**Supplementary Fig. S11b**), though it should be noted that this trend for ATP in particular was not observed in previous experimental studies of related cell-free systems (Karim et al., 2018; Karim & Jewett, 2016). Next, we varied initial concentrations of glucose, acetate, and glutamate (**Figure 5**). While glucose was seen to seen to have the largest effect on final butanol titer, the correlation was not linear. For example, a 62% decrease in initial glucose resulted in a 34% decrease in final butanol titer in the median across parameter sets, with one parameter set even showing an increase in butanol titer from this decreased glucose.

**Figure 5:**
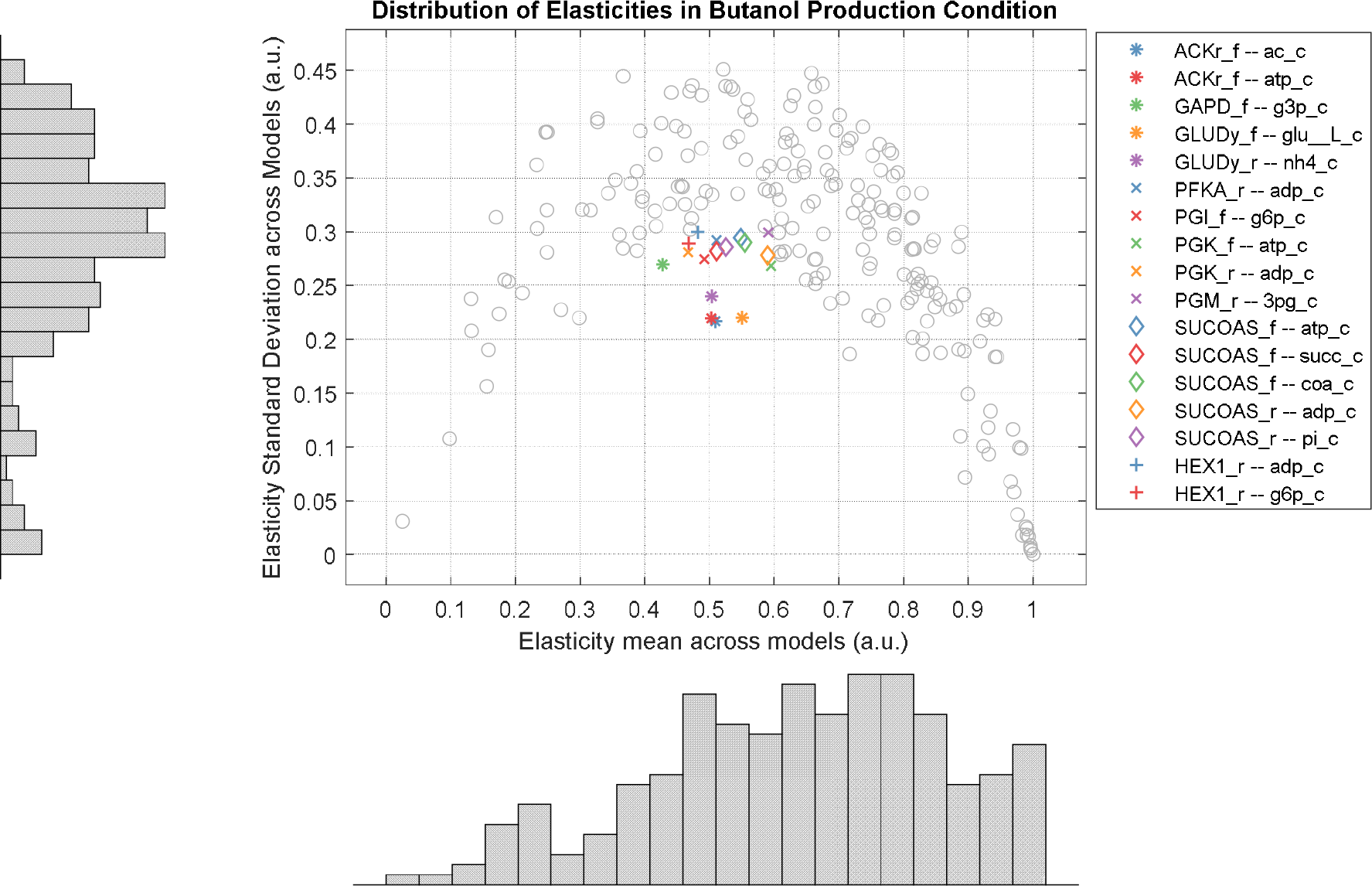
Analysis of parameter variation across models shows small number of highly conserved parameters. Elasticity values for each parameter, which serve as a more informative proxy for absolute parameter values, are plotted with their mean value across the top 20 models in the final ensemble (x axis) versus the standard deviation across the top 20 models (y axis). Each point represents a single elasticity or model parameter, and marginal histograms for both mean and standard deviation show distributions across these elasticities. Elasticity values near 0 represent elasticities for metabolites that are fully saturated (zeroth order kinetics), and values near 1 represent metabolites far below saturation in a linear regime (first order kinetics). Those parameters which a mean elasticity between 0.4 and 0.6, representing a high degree of control over the reaction rate, and a standard deviation below 0.3, representing highly conserved behavior among all models in the ensemble, are highlighted with names given in the legend, denoted by the reaction name (i.e., ACKr for acetate kinase), then the reaction direction (“_f” for forward or “_r” for reverse), then the reaction substrate (i.e., “ac_c” for acetate). A complete list of reaction, substrate, and elasticity names given Elasticity values are calculated as the derivative of reaction rate with respect to metabolite concentration and are normalized with respect to both rate and concentration. as well as the stoichiometry of the metabolite in each reaction. To ensure all values were between 0 and 1, metabolites acting as inhibitors are not shown in this figure.

### 3.4. ANALYSIS OF PARAMETER OPTIMIZATION AND CONSERVATION OF PARAMETER VALUES

Virtually all large-scale kinetic models are vastly underdetermined and individual parameters are not identifiable, thus limiting the interpretability of individual parameter values. Because we used an ensemble approach, we have an opportunity to study how parameter values and network features are constrained – or unconstrained – as a result of undertaking this type of model fitting. For a given parameter, the conserved prediction of similar parameter values across the ensemble may support several conclusions, including increased confidence in a true physiological parameter value, increased structural identifiability due to network structure or reaction mechanisms, or increased practical identifiability due to proximity to observed data. In all cases, these are important factors in model building and analysis, and these results may help increase understanding in these areas.

Here, we measure how tightly or loosely constrained individual parameter values are between the parameter sets in the final ensemble to understand how the data might constrain certain parameter values, or conversely, how different parameter values might give similar model behaviors. However, because individual parameter values are highly context-dependent, and the effect of a parameter on model responses cannot be determined from the absolute parameter value alone, it is instead preferrable to use proxy values for each parameter which are more representative of network behavior.

We first used as a proxy the metric of concentration elasticity, which relates the response in reaction flux to the value of each dissociation constant parameter and normalizes all parameters for better comparison. This metric further allows analysis of the degree to which parameters control various reactions, as elasticity values near 0 signify saturated substrates, elasticities near 1 signify substrate concentrations far below saturation in a first-order regime, and intermediate elasticities represent substrates with more control over reaction rates. After performing this analysis on the ensemble of 20 models, we saw that there were several parameters whose corresponding elasticity values were highly conserved. This result in summarized in **Figure 5**, which plots the mean value of each elasticity against the standard deviation across the 20 models in the final ensemble. Further discussion of specific examples of conserved elasticities is given in **Supplementary Table S2**, and the full presentation of elasticities for all reactions across all models are shown in **Supplementary Table S4** and **Supplementary Fig. S12**.

Next, we used as a proxy the metric of parameter sensitivity with respect to NADH oxidative flux (**Supplementary Methods S10**). Because this value is physiologically relevant and yet was not directly constrained or favored by the model fitting process, this provides an additional unbiased metric to analyze the extent to which parameter values are constrained. Again, there were several parameters whose sensitivity to this behavior were relatively conserved, which is summarized in **Supplementary Table S3**, with all values given in **Supplementary Table S5** and shown in Supplementary Fig. S13.

## 4. DISCUSSION

### 4.1. SIGNIFICANCE OF WORK

In this work, we successfully fit a large-scale dynamic model to multiple conditions of experimental time course metabolite measurements. The resulting model captured complex, unintuitive interactions between various branches of metabolism and was able to further provide predictions as to the metabolic phenomena underlying these effects. The ensemble approach was leveraged to provide quantitative measurements of the uncertainty of these predictions; somewhat surprisingly, the predictions around both the metabolic behaviors in the training data, as well as the predicted effects of hypothetical enzyme level changes, were much more conserved within the ensemble than initially expected, demonstrating high certainty in these predictions. The final model produced several other optimization strategies with varying degrees of ensemble uncertainty that may be used in future work to increase butanol production and further retrain this model.

### 4.2. ADVANTAGES AND LIMITATIONS

The most significant advantage of the methodology of thiswork was the use of the ensemble approach. First, this approach allows for a thorough sampling of parameter space, which was seen to be necessary to provide adequate initial points for subsequent optimization (**Supplementary Fig. S4**). Second, it allows quantification of the uncertainty in both model parameters and model predictions, such as the increased confidence in the predicted effect of butanol enzyme level changes. Third, the ensemble approach allows for uncertain predictions to be hedged, such that even predictions which vary between models can provide utility. For example, lower-confidence predictions can be tested to improve butanol production, and in the worst-case scenario will still provide valuable training data to refine the ensemble. This ease of re-optimization by local optimization is also a significant advantage of this work which can increase the speed of model-building in the design-build-test-learn cycle.

Another advantage of this work is the scalability and utility of additional experimental data. The above analysis found that the most highly constrained parameters were in reactions near experimentally measured metabolites, which implies that currently under-constrained parameters can be fine-tuned simply by obtaining more experimental measurements. The resulting tightening of parameter constraints should give more accurate and conserved predictions across models. Simultaneously, a well-fit model was obtained in this work despite measurements from relatively few metabolites, so while additional measurements may have benefits, they should not be seen as a reason not to undertake similar model-fitting efforts.

One primary limitation in this work, common to many large-scale models, is the manual process of model selection. While there have been early efforts to use automated model reconstruction tools on kinetic models (van Rosmalen et al., 2021), their widespread use remains limited to constraint-based models with steady-state data (Mendoza et al., 2019), and while parameter identifiability tools have seen great improvements, they still scale poorly for larger models, especially those with complex rate forms (Villaverde et al., 2019). Therefore, model selection remains an iterative process which must balance reducing the number of parameters with leaving enough reactions to allow the model sufficient flexibility to fit the data. For the “bottom-up” parameterization in this work, the choice of reactions was further complicated by the choice of parameter priors, especially where data could be fit in multiple ways. For example, our model included both hexokinase (HEX) and the glucose phosphotransferase system (PTS) as reactions to consume glucose. While we chose higher rate constants for HEX, as we believed it unlikely that the multi-step, membrane-bound PTS was highly active in cell-free extracts, a different choice may have led to a different final model. Similarly, model predictions of succinate production from glutamate versus from TCA flux may have been biased by parameter priors. Despite this potential uncertainty, in this work we saw that these two decisions had minimal impact: in the first case, the primary reaction for glucose consumption did not significantly affect model predictions surrounding butanol optimization (**Supplementary Fib. S14**), and in the second case, nearly all parameter sets, once optimized, utilized glutamate as the primary source of succinate flux. Regardless, this fortunate result is not guaranteed for future efforts, and so careful consideration of model selection, along with the potential use of automated model construction or parameter identifiability methods, model reduction methods (Strutz et al., 2019) and increasingly approximate rate forms (Du et al., 2016; St John et al., 2019) should all be considered.

In conclusion, while there are certainly improvements to be made in future modeling efforts, this work demonstrates the successful parameterization of a large-scale dynamic kinetic model to capture complex metabolic data. The resulting model has increased realism and confidence due to improved literature value parameter priors and rigorously applied thermodynamic constraints. This model was used to gain new causal understanding of metabolism in cell-free systems and how this metabolism interacts with the engineered butanol pathway. This understanding was further leveraged to provide strategies to optimize butanol titers, including the prediction that the alcohol/aldehyde enzyme was the primary bottleneck of the butanol pathway, which itself was unique and unanticipated due to its high conservation across models despite the lack of direct training data. Lastly, the final trained model was used for a large-scale analysis of the degree to which individual parameter values are constrained during model fitting, which has broad implications in kinetic models in general. We hope that the success, applicability, and general ease of use of these methods and results will inspire additional experiments to measure dynamic behaviors around engineered pathways in metabolism for the purpose of continued model building and improved metabolic understanding.

## CONTRIBUTION STATEMENT

Jacob Martin: Conceptualization, Methodology, Software, Formal analysis, Writing – Original Draft.

Blake J. Rasor: Data Collection, Writing – review & editing.

Jon DeBonis: Methodology, Software, Writing – review & editing. Ashty Karim: Data Collection, Writing – review & editing.

Michael Jewett: Supervision, Funding acquisition, Writing – review & editing. Keith Tyo: Conceptualization, Supervision, Writing – review & editing.

Linda Broadbelt: Conceptualization, Supervision, Writing – review & editing.

## ACKNOWLEDGEMENTS

We would like to thank Prof. Niall Mangan, Prof. Matthew Plumlee, and Prof. Andreas Waechter for project guidance and helpful discussions. Thanks to Jon Strutz and Kevin Shebek for helpful discussions and assistance with software and eQuilibrator 3.0.

This research was supported in part through the computational resources and staff contributions provided for the Quest high performance computing facility at Northwestern University which is jointly supported by the Office of the Provost, the Office for Research, and Northwestern University Information Technology. This work was funded by the U.S. Department of Energy Office of Science, Biological and Environmental Research Division (BER), Genomic Science Program (GSP) under Contract No. DE-SC0018249. M.C.J. gratefully acknowledges the David and Lucile Packard Foundation and the Camille Dreyfus Teacher–Scholar Program. J.P.M. was supported by a National Institutes of Health Training Grant (T32GM008449) through Northwestern University’s Biotechnology Training Program. B.J.R. was supported by a National Defense Science and Engineering Graduate Fellowship (Award ND-CEN-017-095).

