## Supplementary material for "Dynamic Kinetic Models Capture Cell-Free Metabolism for Improved Butanol Production": cell_free_kinetic_model-Supplementary_Figures_Final.docx

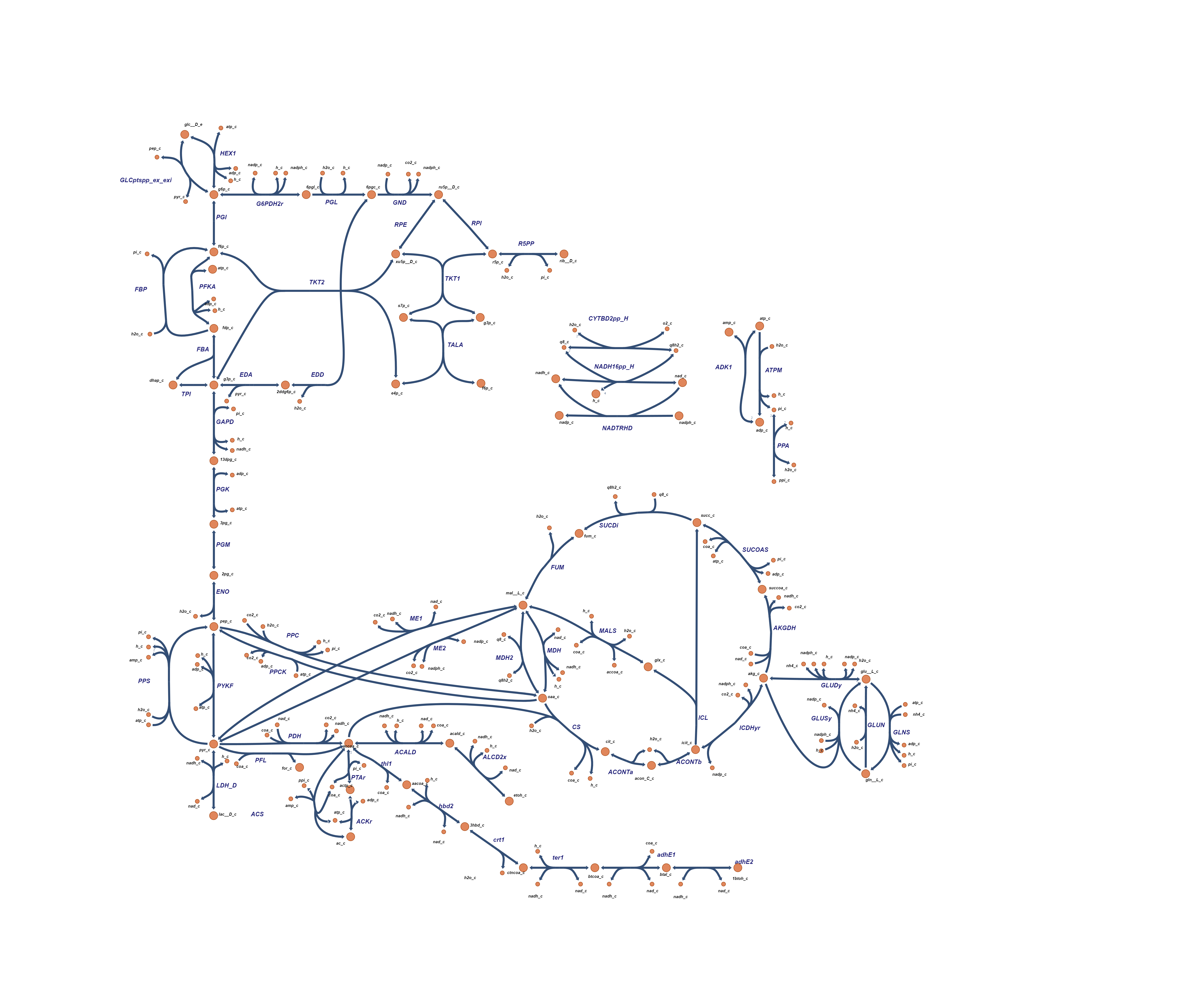

**Supplementary Figure S1: Detailed metabolic network map.** The full metabolic network is shown here. The network includes the BiGG *E. coli* core model, as well as additional reactions in the pentose phosphate pathway (EDD, EDA, and R5PP),acetyl-CoA synthase (ACS), ubiquinone-dependent malate dehydrogenase (MDH2), and the butanol pathway reactions (thl1, hbd2, crt1, ter1, adhE1, and adhE2). Reactions with redundant stoichiometry (FRD7) or those made redundant by the lack of compartments (THD2, ATPS4r) were removed, as well as transport reactions between compartments and exchange reactions removing species from the system. Escher was used to generate the network image and for visualization (King et al., 2015).

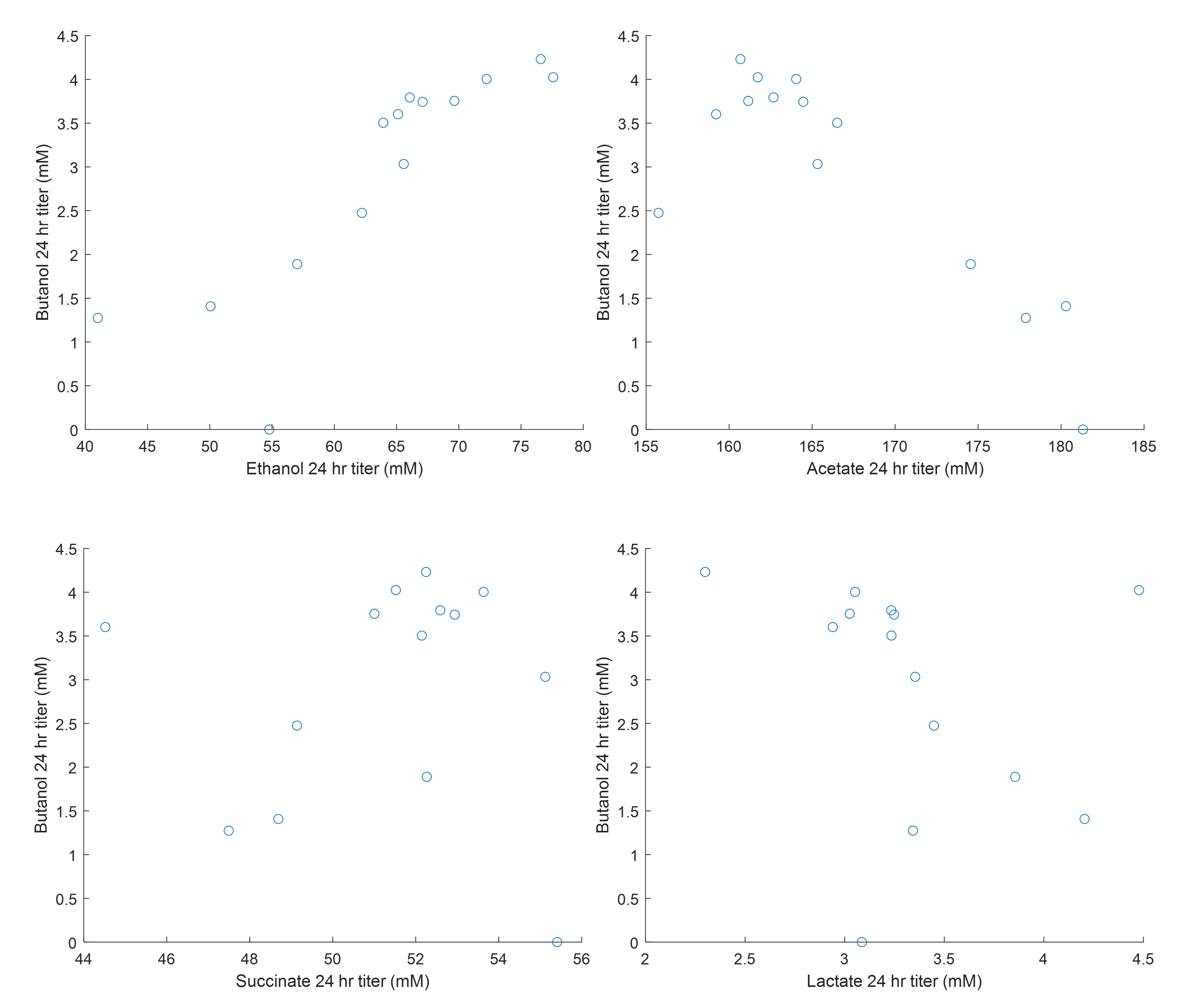

**Supplementary Figure S3: Correlation of product titers across preliminary experiments.** Unpublished data show moderate correlations in 24-titers of ethanol, acetate, succinate, and lactate with respect to 24-hours butanol titers. Each of the 17 points is an experimental condition in which only the enzymes in the butanol pathway were adjusted.

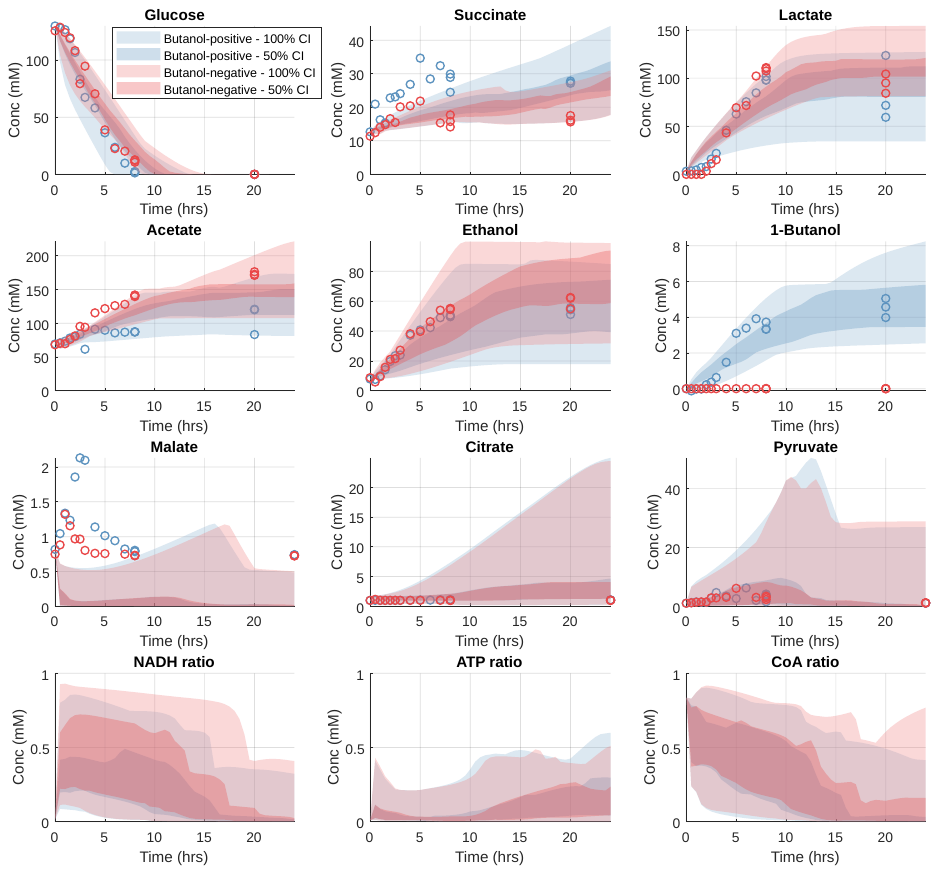

**Supplementary Figure S3: Model fit from ensemble screening alone.** By creating an ensemble of 10^7^ parameter sets and screening to the top 20 by weighted RMSE, the model obtained adequate fit to metabolite data in the butanol-positive condition (blue circles). However, shifts in the butanol-negative condition (red circles), especially in measurements for succinate and acetate, were not all able to be captured by any of the top 20 parameter sets. Light and dark shaded regions denote the model-simulated concentrations in which 100% and 50% of parameter sets fall, respectively.

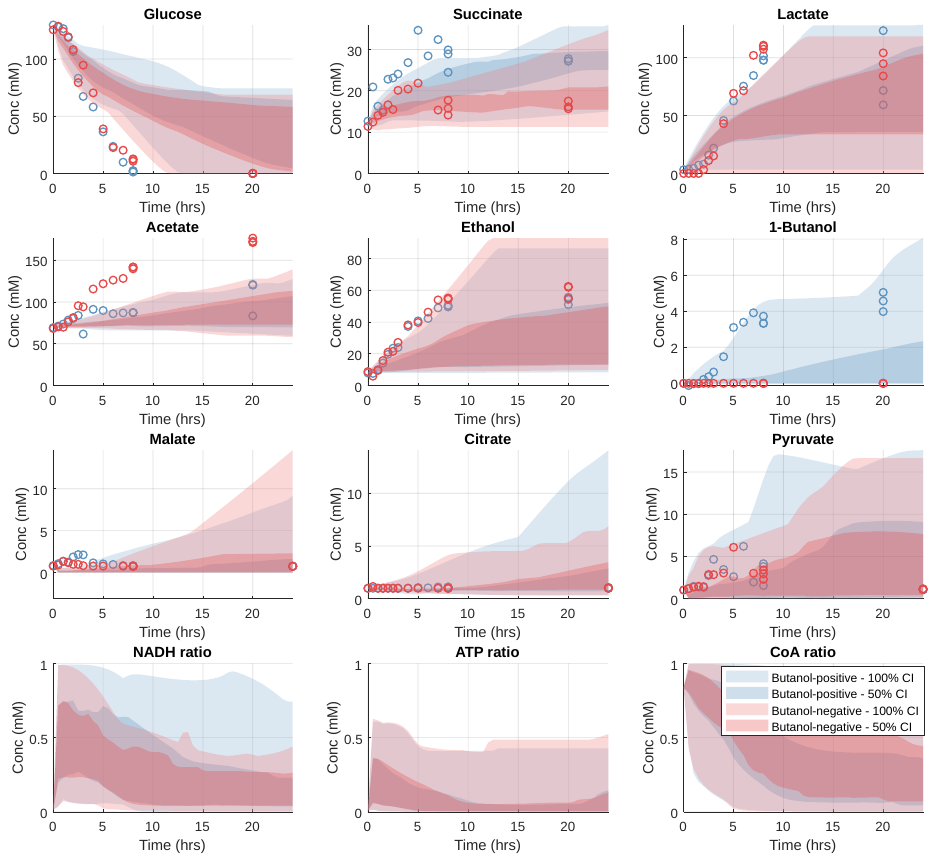
**Supplementary Figure S4: Model fit from optimization alone.** 200 parameter sets were randomly sampled (from the final, updated parameter priors) without any additional screening and fed into the patternsearch algorithm for local optimization. These unscreened initial points were unable to find optimized parameter sets that adequately fit the shifts in acetate and succinate seen in the butanol-negative condition (red circles).

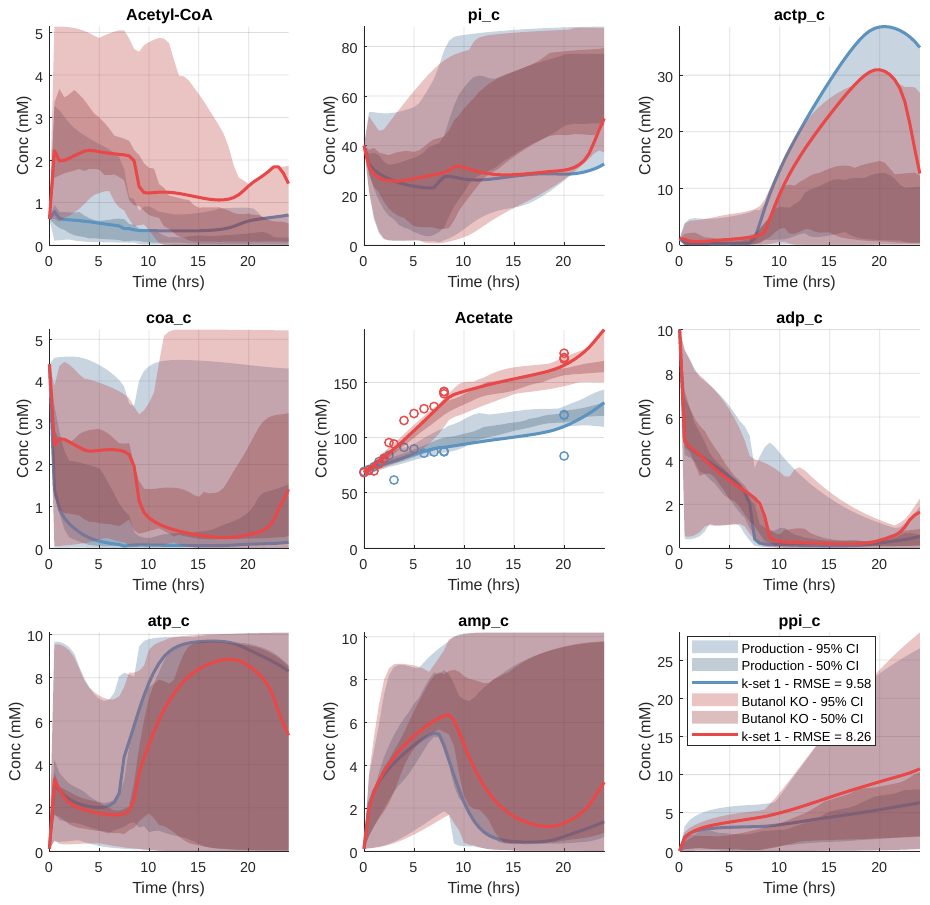

**Supplementary Figure S5: Acetate pathway metabolites.** The concentration of each metabolite in the acetate pathway was analyzed to understand what factors led to increased acetate production in the butanol-negative pathway. In the direction of acetate production, the only substrate whose concentration increased (or product whose concentration decreased) was acetyl-CoA, which was therefore found to be responsible for the increased acetate production.

**(a)**
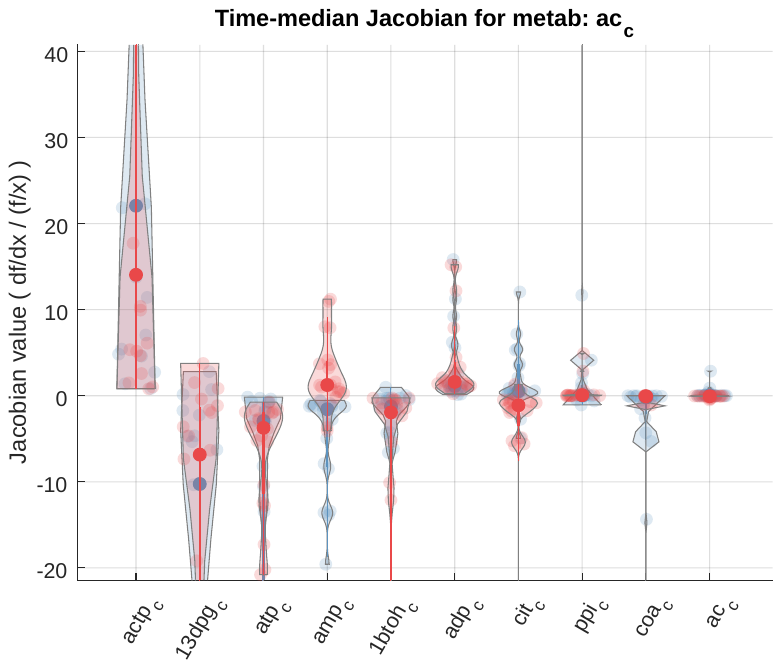

**(b)**
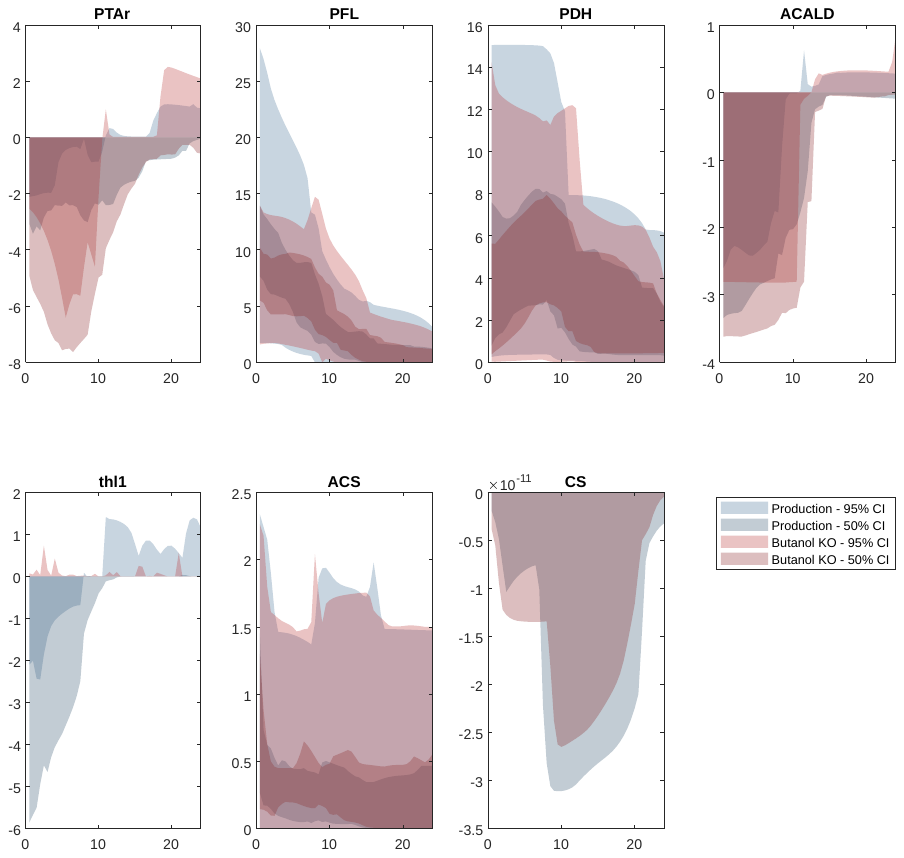

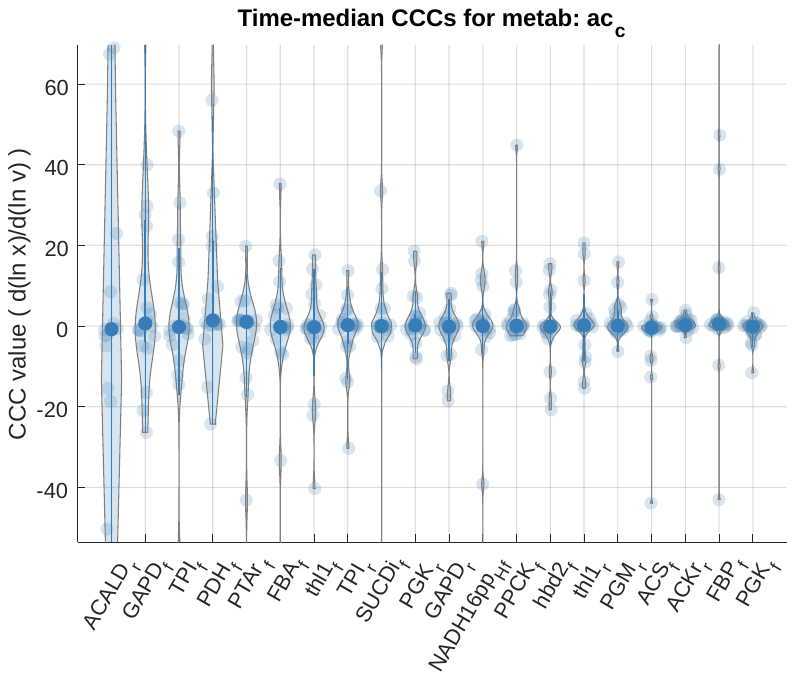

**(c)**

**Supplementary Figure S6: Analysis of acetate production.** (a) Jacobian values are averaged in the 0.5 hour to 7.5 hour window. Each dot shows the time-averaged Jacobian for each model, and violins summarize the distribution across models for both the production condition (blue) and the knockout condition (red). Within the top species (ranked by median absolute value across models) for control of acetate is butanol (1btoh_c_), showing how butanol pathway flux influences the acetyl-CoA pool and ultimately acetate production. (b) This shift in acetyl-CoA levels is also shown by calculating the instantaneous rate of change of acetyl-CoA concentration, broken down by each reaction producing or consuming acetyl-CoA; thiolase (thl1), the first step in the butanol pathway, is the only reaction which significantly raises the availability of acetyl-CoA from the production (blue) to the knockout (red) condition, and therefore is responsible for the increase in acetyl-CoA which leads to increased acetate production. (c) The top 20 (across models) concentration control coefficients include several reactions within acetate metabolism, as well as the first two steps of the butanol pathway (thl1 and hbd2).

(a)
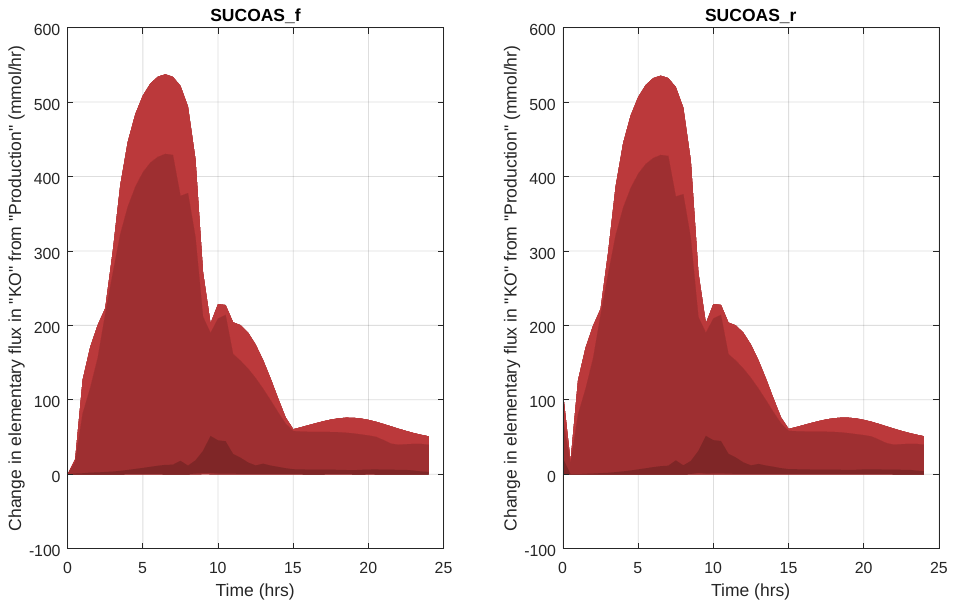

(b)
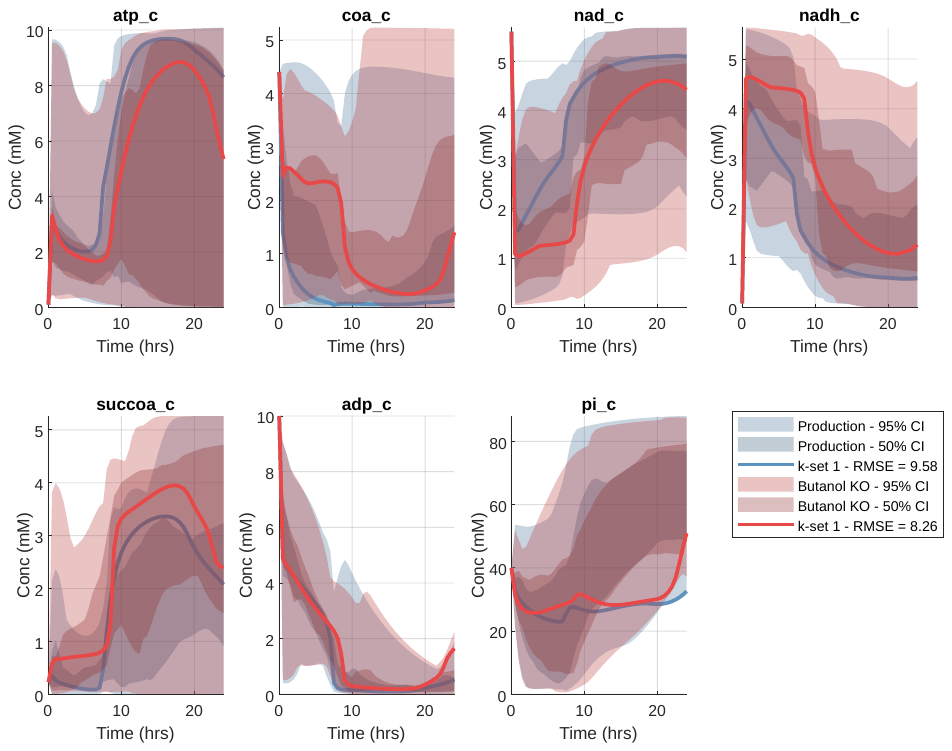

**(c)**
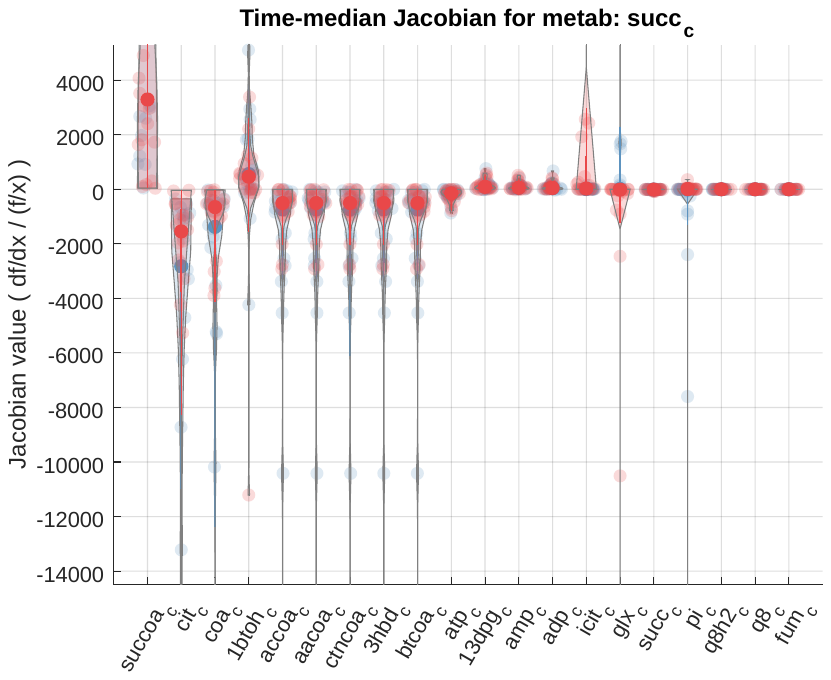

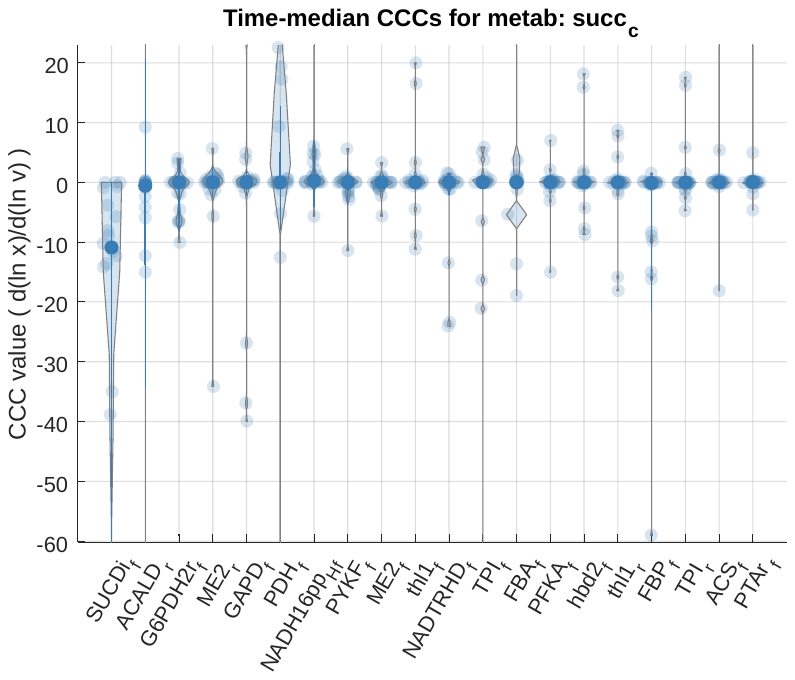

**(d)**

**Supplementary Figure S7: Analysis of succinate production.** (a) Changes in elementary flux in the knockout condition are shown. Decomposing SUCOAS flux into elementary forward and reverse steps shows that the decrease in succinate production in the butanol knockout condition comes from increased elementary flux from succinate to succinyl-CoA, labeled here as SUCOAS_f, and not from decreased flux from (SUCOAS_r). The darkest region the changes in flux for the median 50% of models, the middle region contains 90% of models, and the lightest region contains 100% of models (b) This increase in SUCOAS_f, whose substrates are succinate, CoA, and ATP, is only attributable to an increase in CoA in the knockout condition, as ATP and succinate do not increase, nor do the metabolites used in AKGDH (c) The Jacobian of succinate, which shows how small perturbations in other metabolites will affect succinate concentration, show that succinate is very sensitive to free CoA levels. While citrate (cit_c_) is also found to be sensitive, this is partially due to the normalized Jacobian being inflated from low baseline levels of citrate, and flux from citrate to succinate is low in nearly all models. (d) Concentration control coefficients over succinate show that succinate, compared to acetate, shows relatively low levels of direct control by the butanol pathway reactions, and is instead largely controlled by the levels of free CoA.

**(a)**

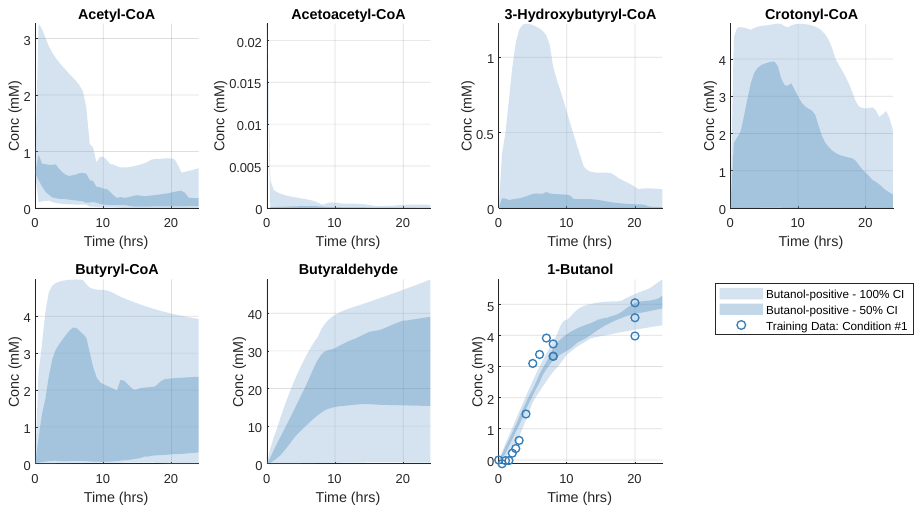

**(b)**

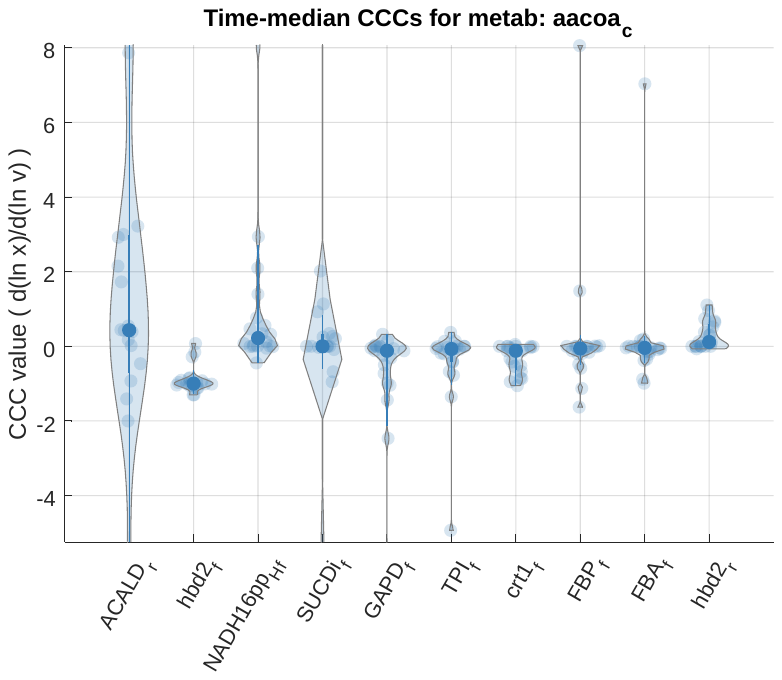

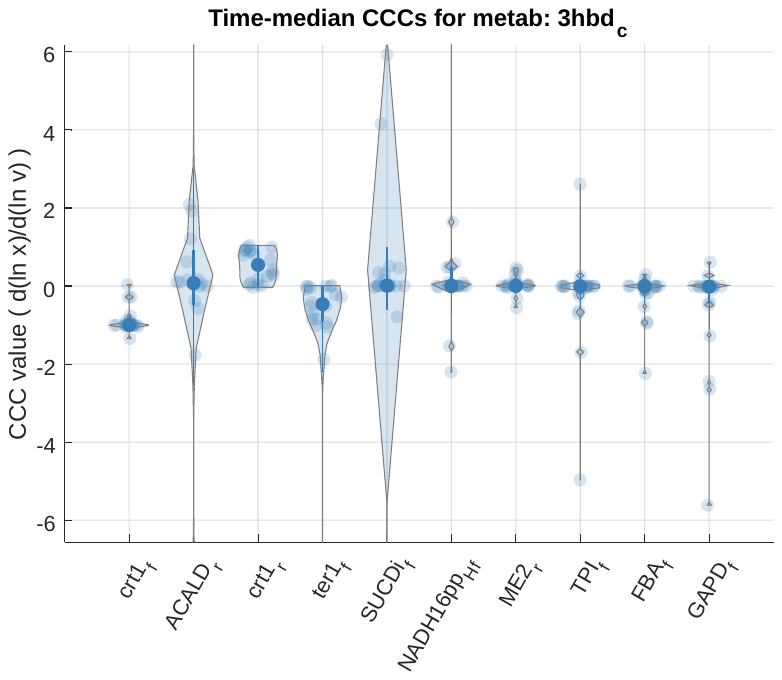

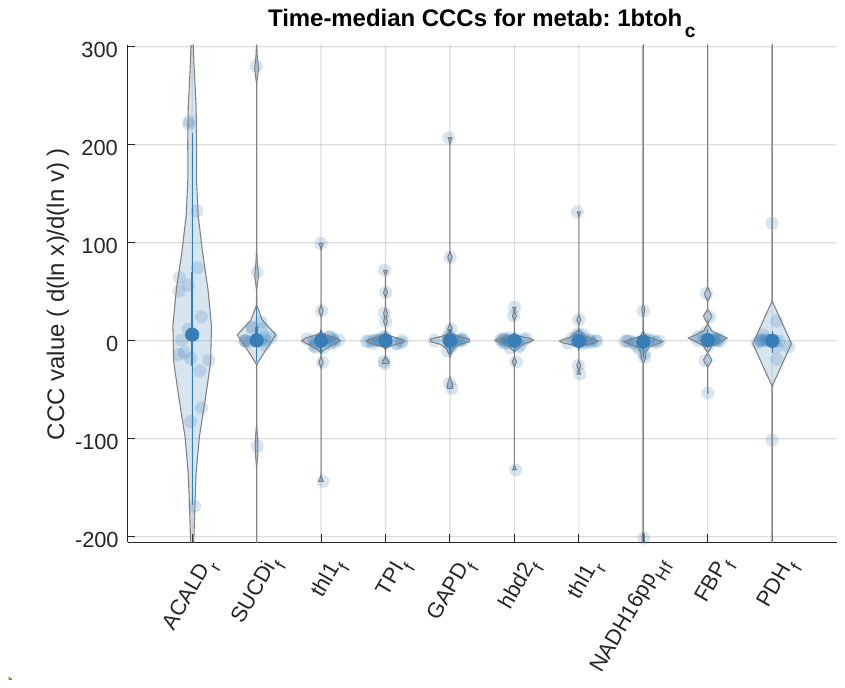

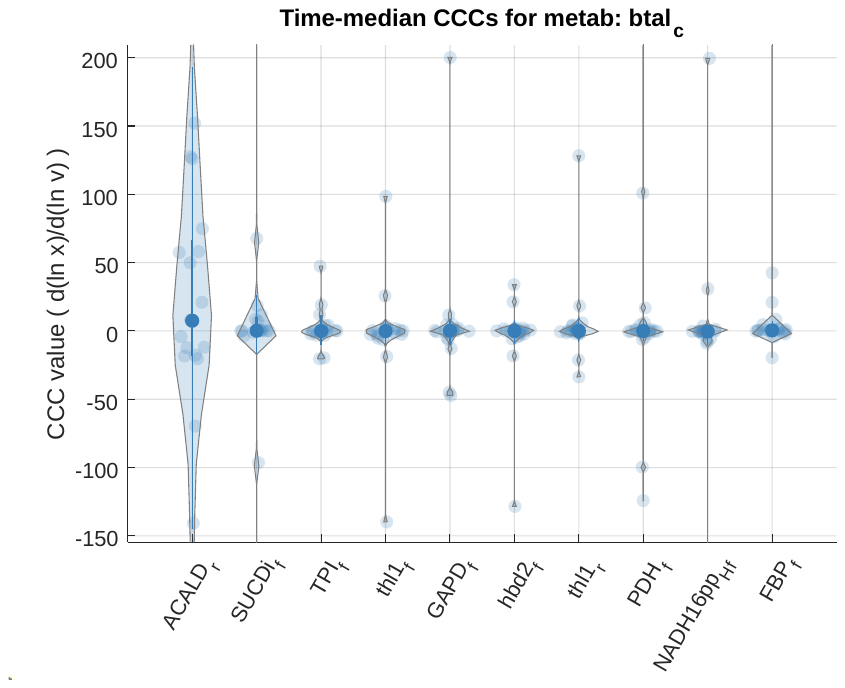

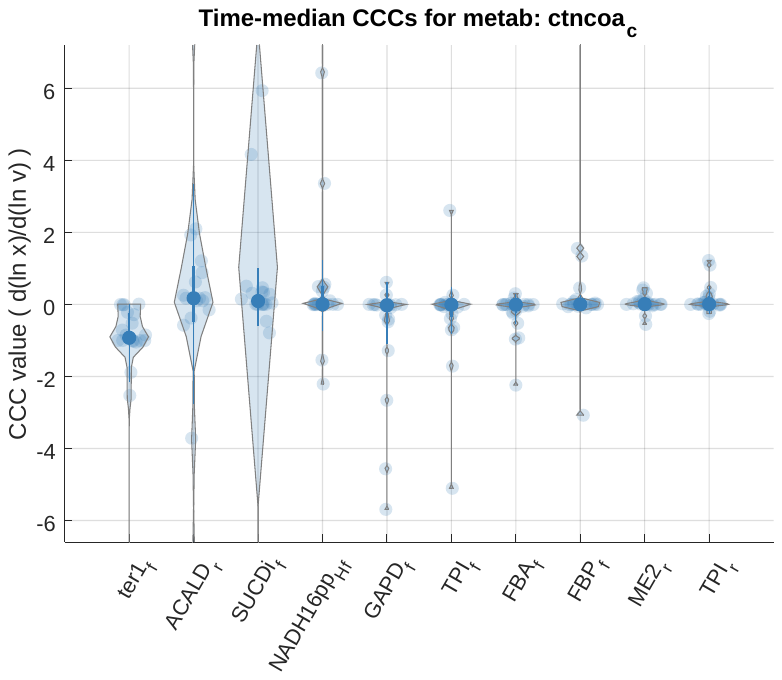

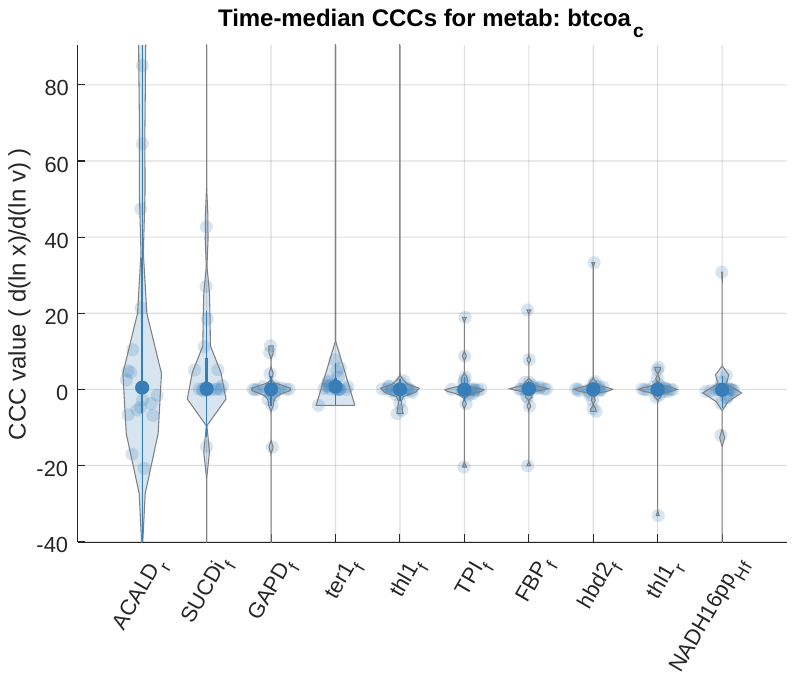

**Supplementary Figure S8: Analysis of the butanol pathway.** (a) Model-predicted timecourses of each of the butanol pathway intermediates are given by the top 20 optimized parameter sets. Significant accumulation of butyryl-CoA and butyraldehyde are predicted. (b) Calculated concentration control coefficients for each metabolite in the butanol pathway are given, where columns are the top 10 reactions ranked by median magnitude of control over each metabolite. Full descriptive metabolite names are given in Supplementary Table S7.

**(a)**

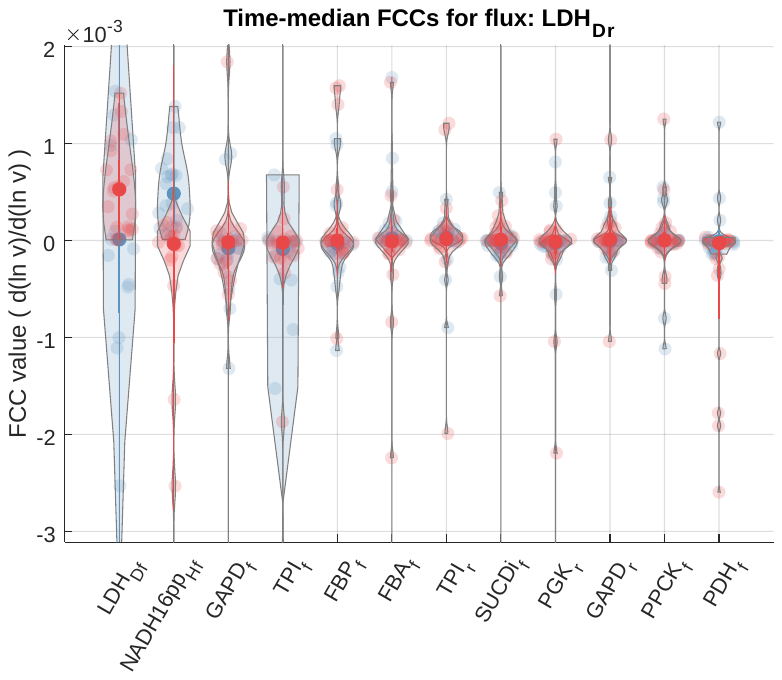

**(b)**

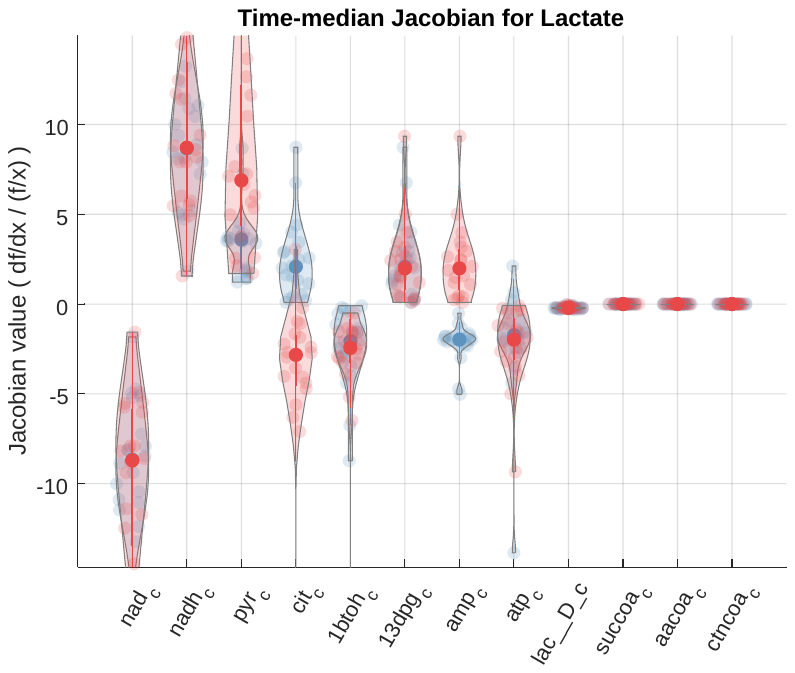

**Supplementary Figure S9: Analysis of lactate production.** (a) Flux control coefficients for lactate dehydrogenase (LDH) (in the direction of lactate production) show that LDH flux is not strongly affected by changes in other model fluxes. (b) Jacobian values for lactate show that changes in lactate concentration are more strongly impacted by changes in NADH or NAD^+^ concentrations than changes in pyruvate concentration. For each panel, the top 12 most important factors shown, as ranked by median absolute value across the final 20 optimized parameter sets.

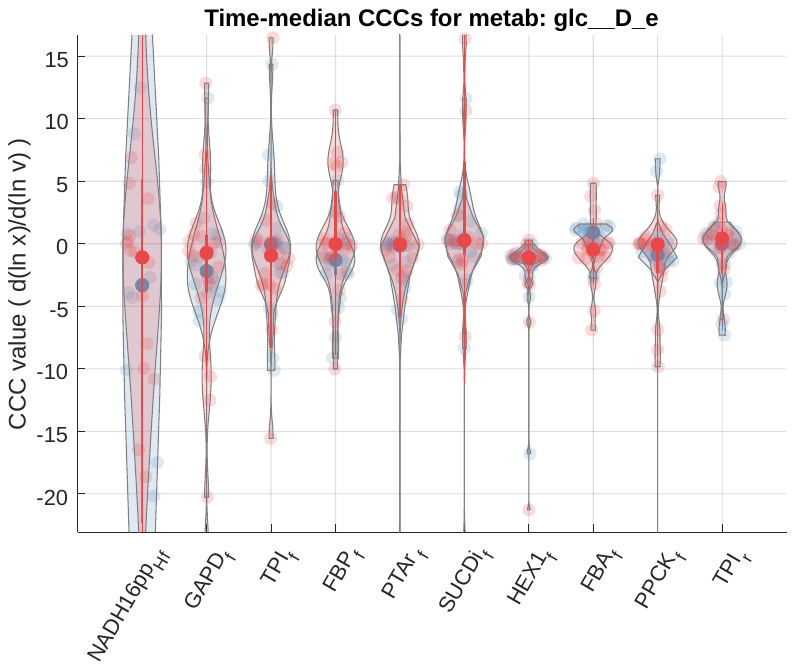

**Supplementary Figure S10: Analysis of glycolysis.** Distribution of concentration control coefficients for control of glucose by the top 10 reactions. Blue violins indicate the range of control coefficients across the final 20 parameter sets in the butanol-positive condition, and red violins indicate the butanol-negative condition.

**(a)**
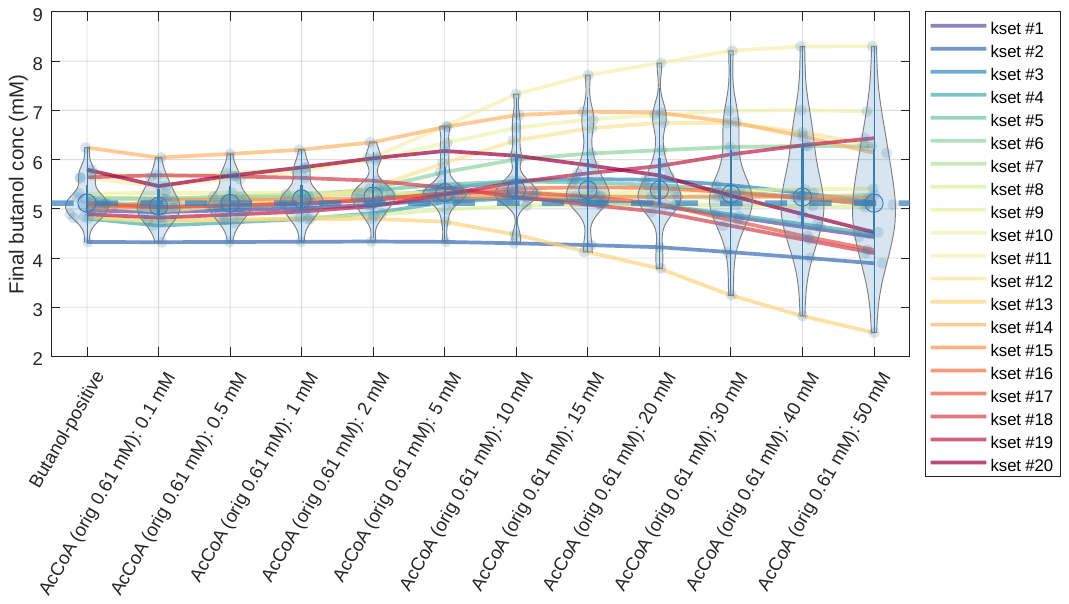

**(b)**
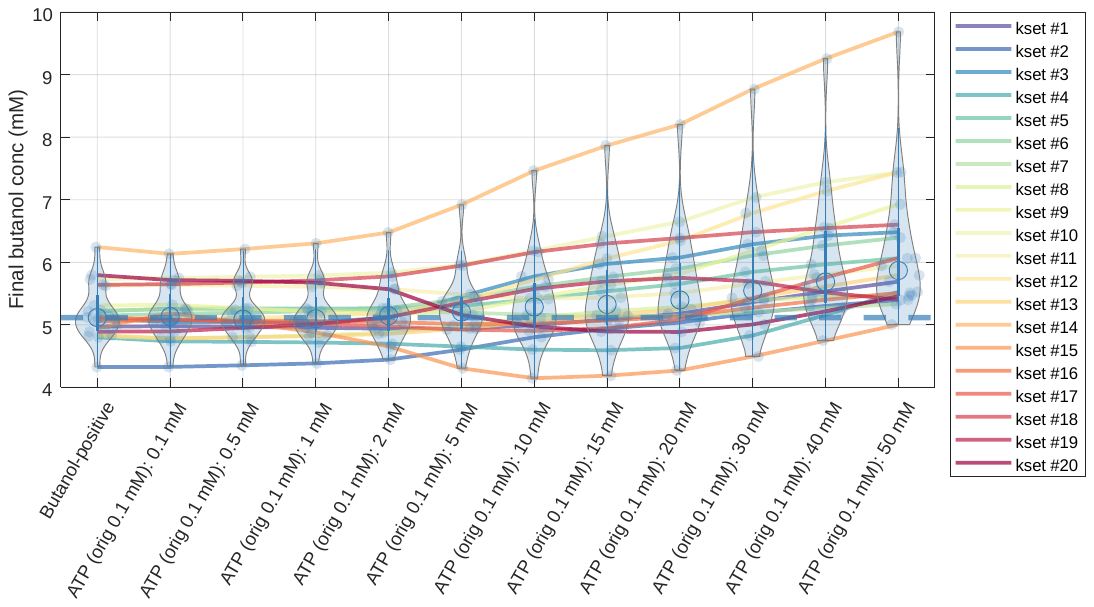

**(c)**
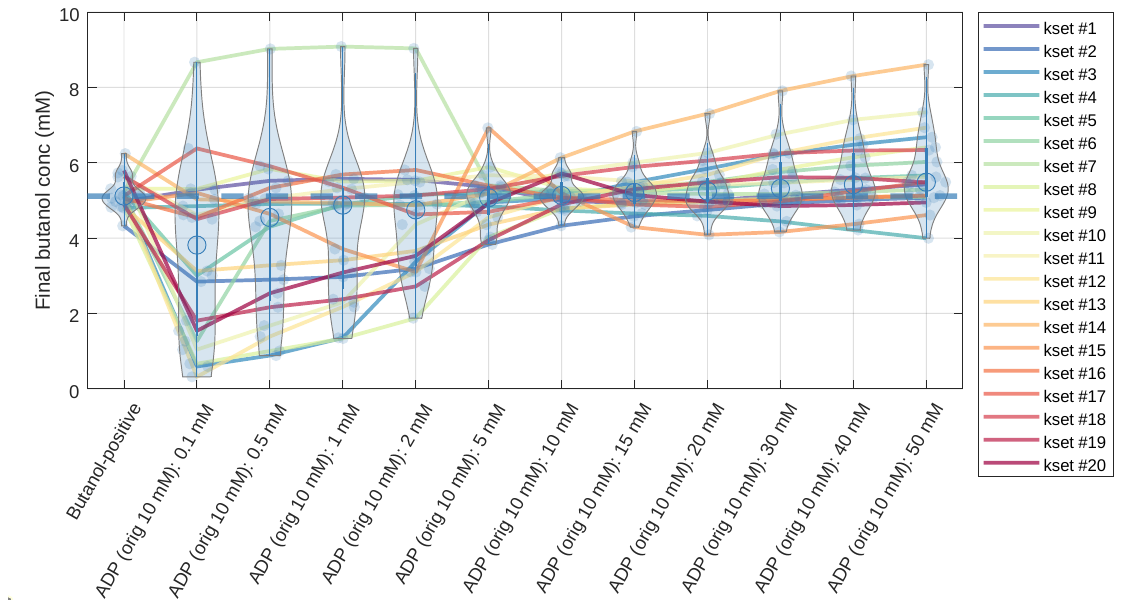

**(d)**
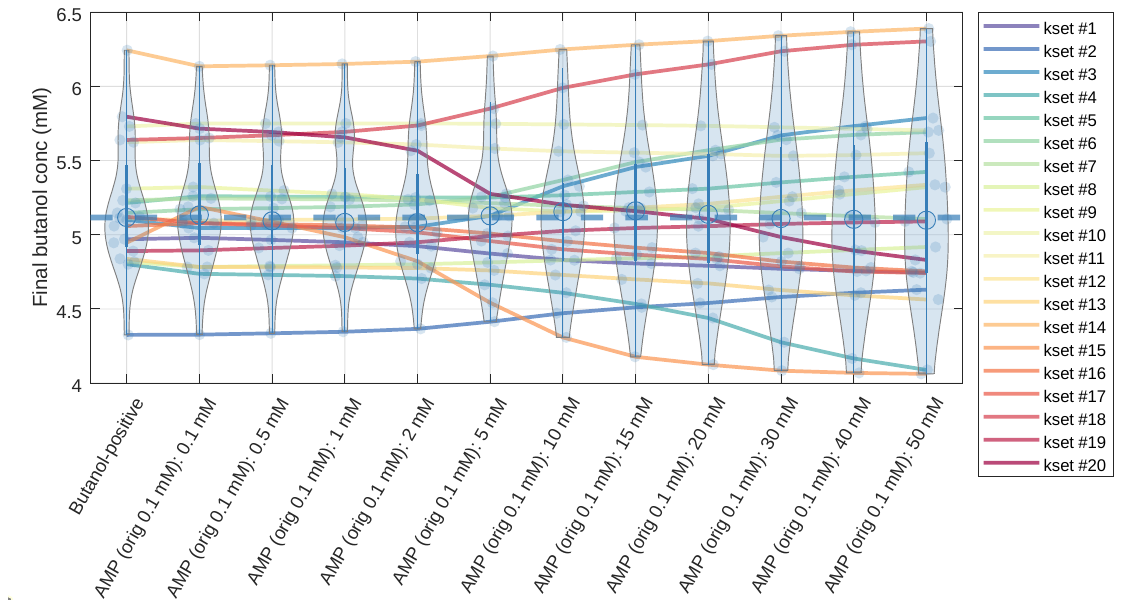

**(e)**
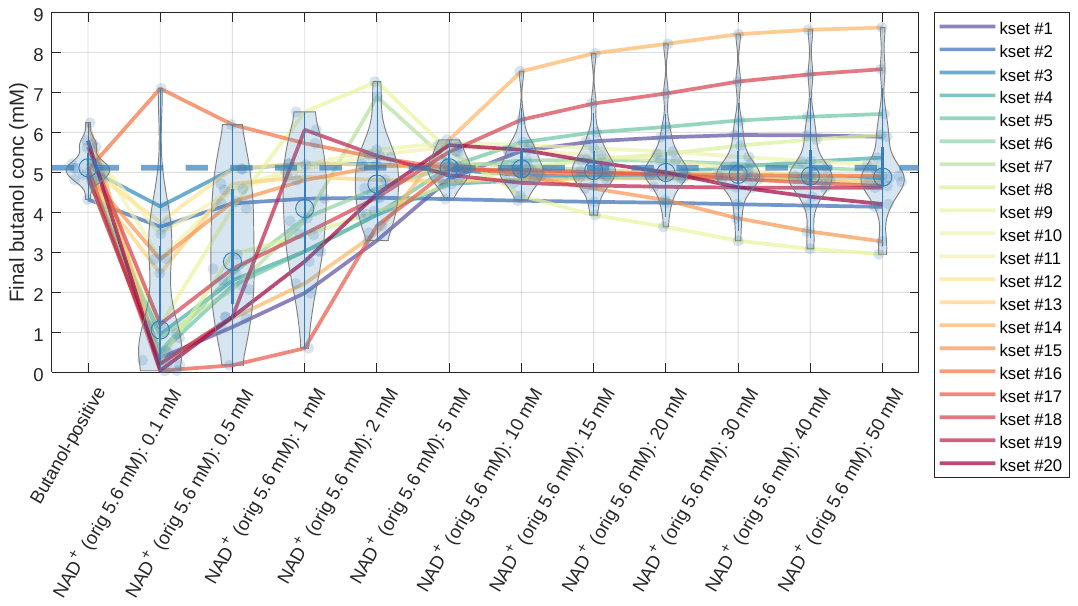

**(f)**
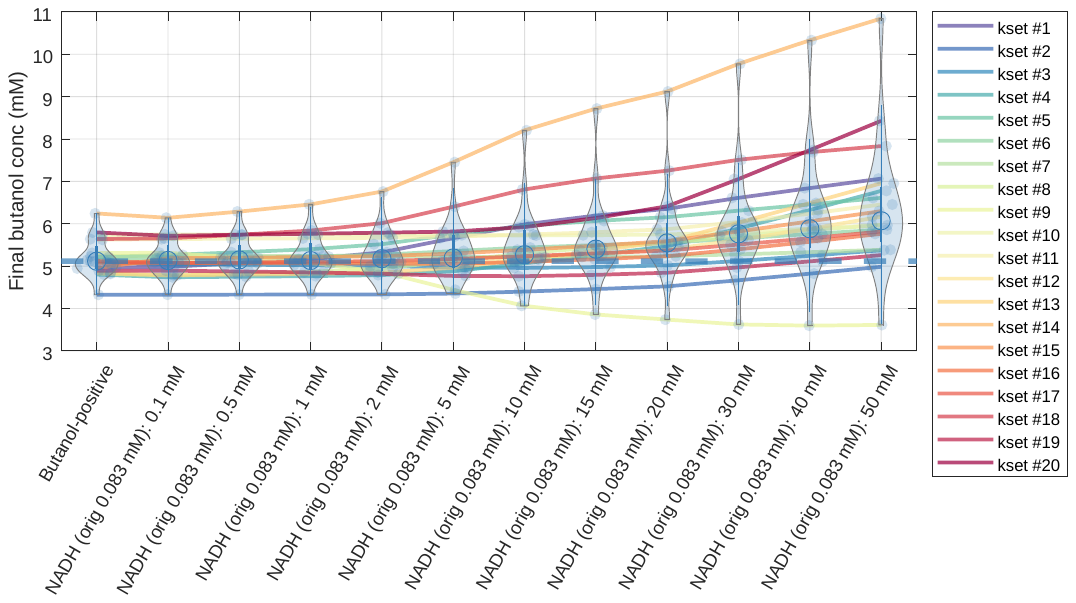

**(g)**
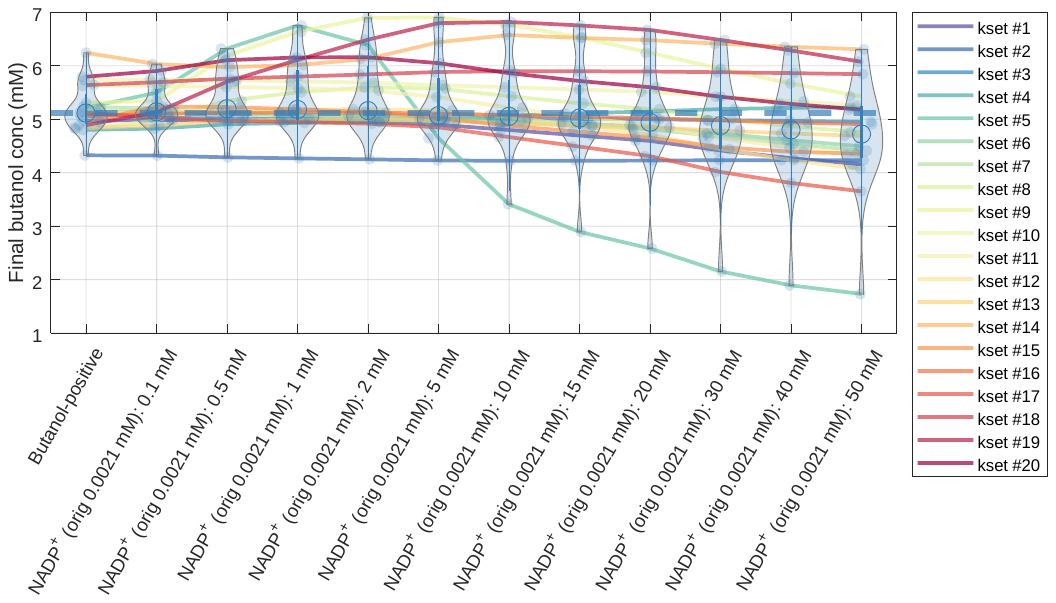

**(h)**
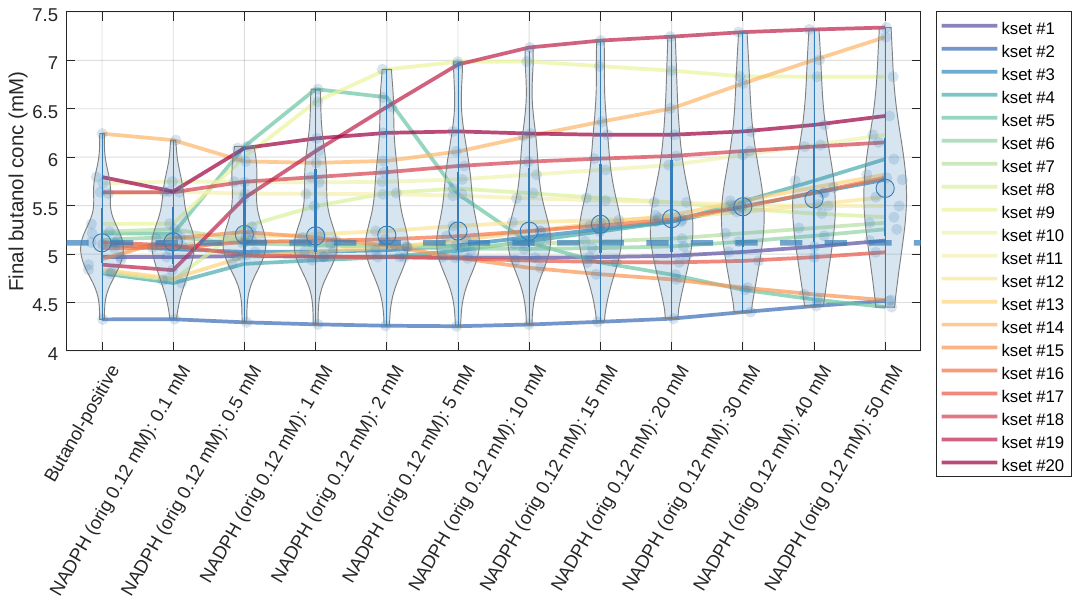

**(i)**
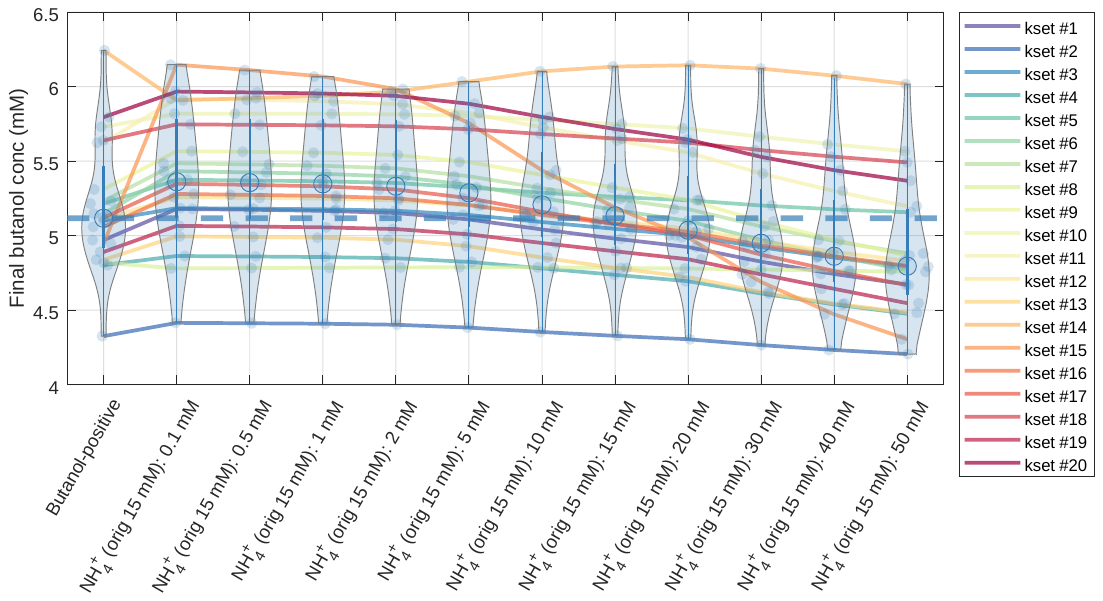

**(j)**
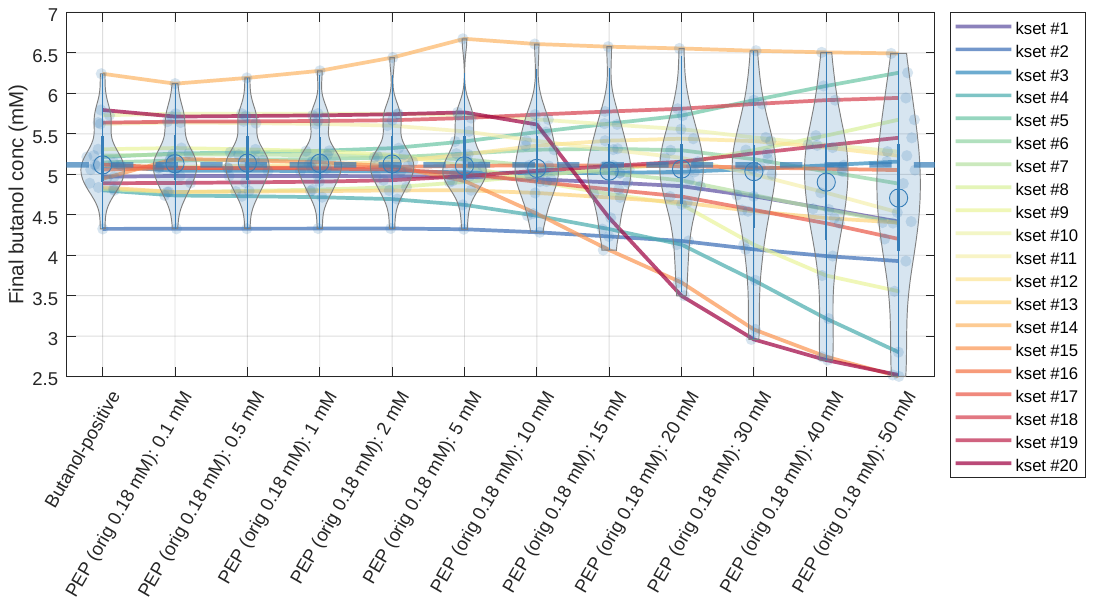

**(k)**
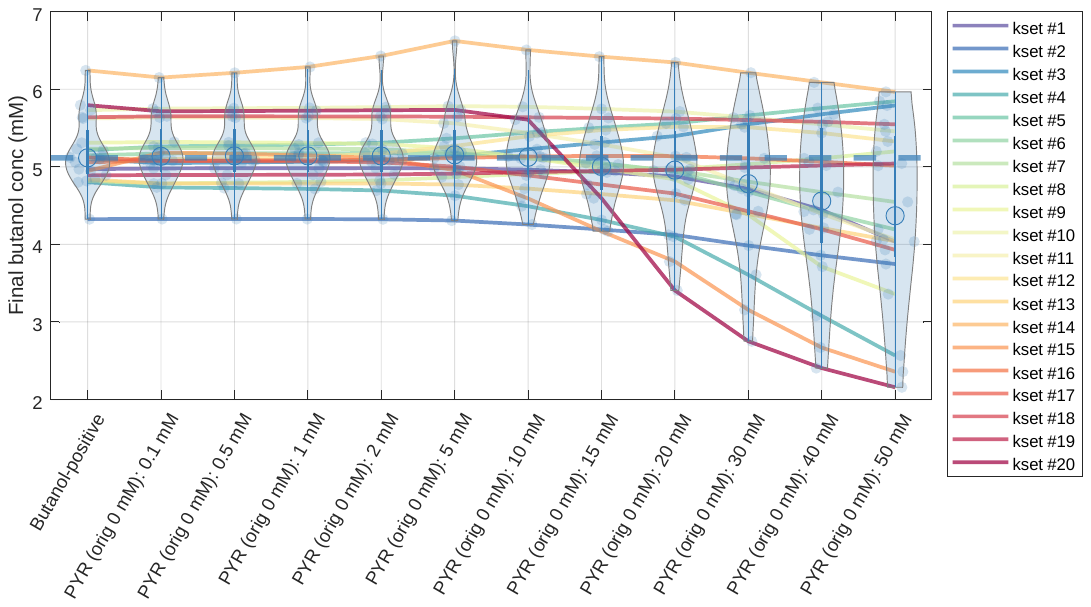

**(l)**
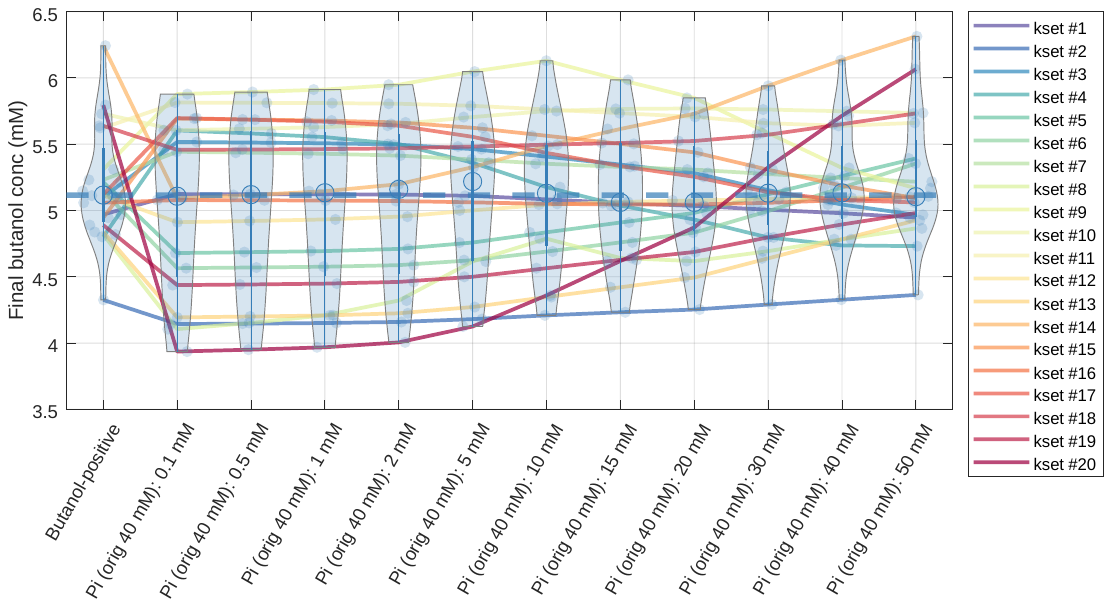

**(m)**
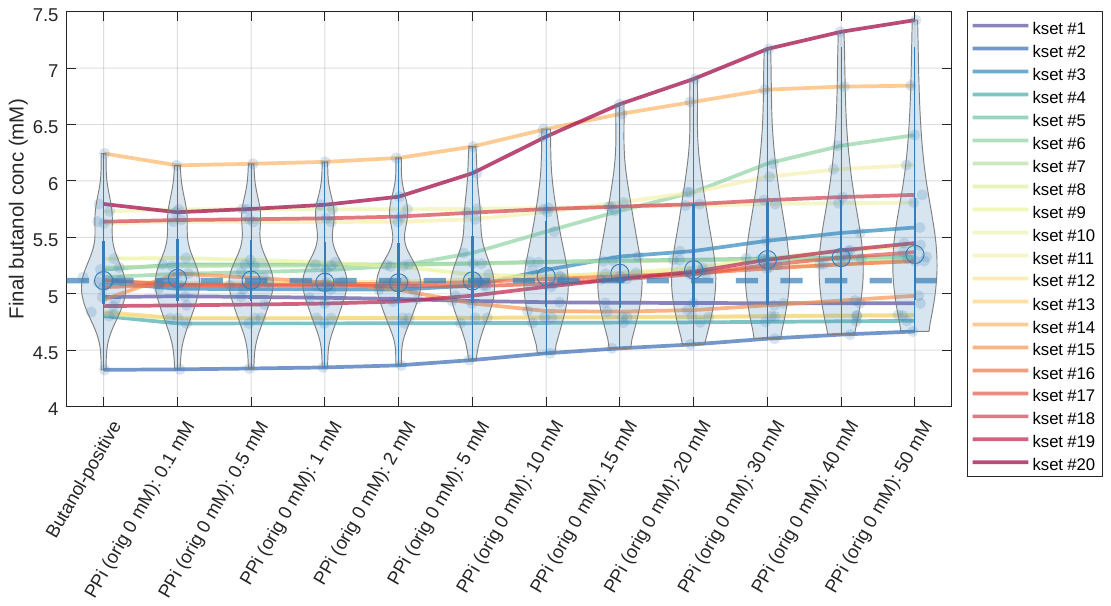

**(n)**
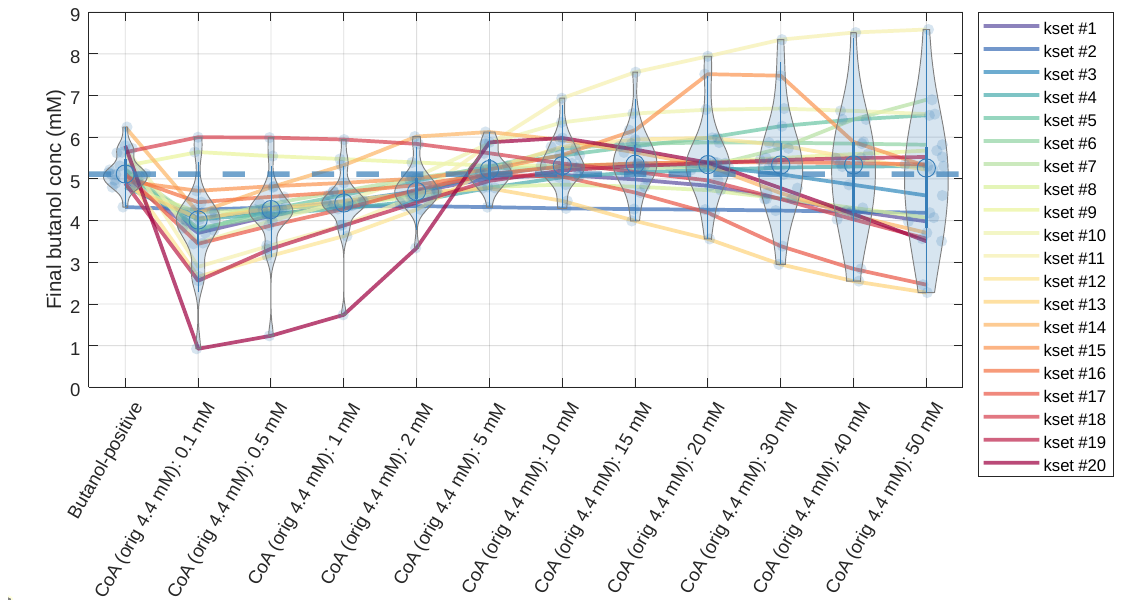

**Supplementary Figure S11: Model predictions of cofactor perturbations.** Conditions with increasing initial concentrations of each model cofactor were simulated and the resulting 20-hour butanol titer was plotted. Violins were used to show the distribution across all 20 parameter sets, with a light blue dot denoting each parameter set prediction and the median prediction shown as an unfilled blue circle. The median production in the baseline butanol-positive condition is shown as a dotted blue line. Initial concentrations in the simulations were titrated at 0.1, 0.5, 1, 2, 5, 10, 15, 20, 30, 40, and 50 mM. These simulations were performed on a single species at a time, and were tested on (a) acetyl-CoA, (b) ATP, (c) ADP, (d) AMP, (d) NAD^+^, (e) NADH, (f) NADP^+^, (g) NADPH, (h) NH_4_^+^, (i) PEP, (j) pyruvate, (k) phosphate, (m) pyrophosphate, and (n) CoA. The predictions of each parameter set (k-set) between conditions is shown as a solid colored line to better show the behavior of individual models as each initial concentration is increased. The concentration of each titrated species in the butanol-positive condition is given in parentheses in the x-axis labels.

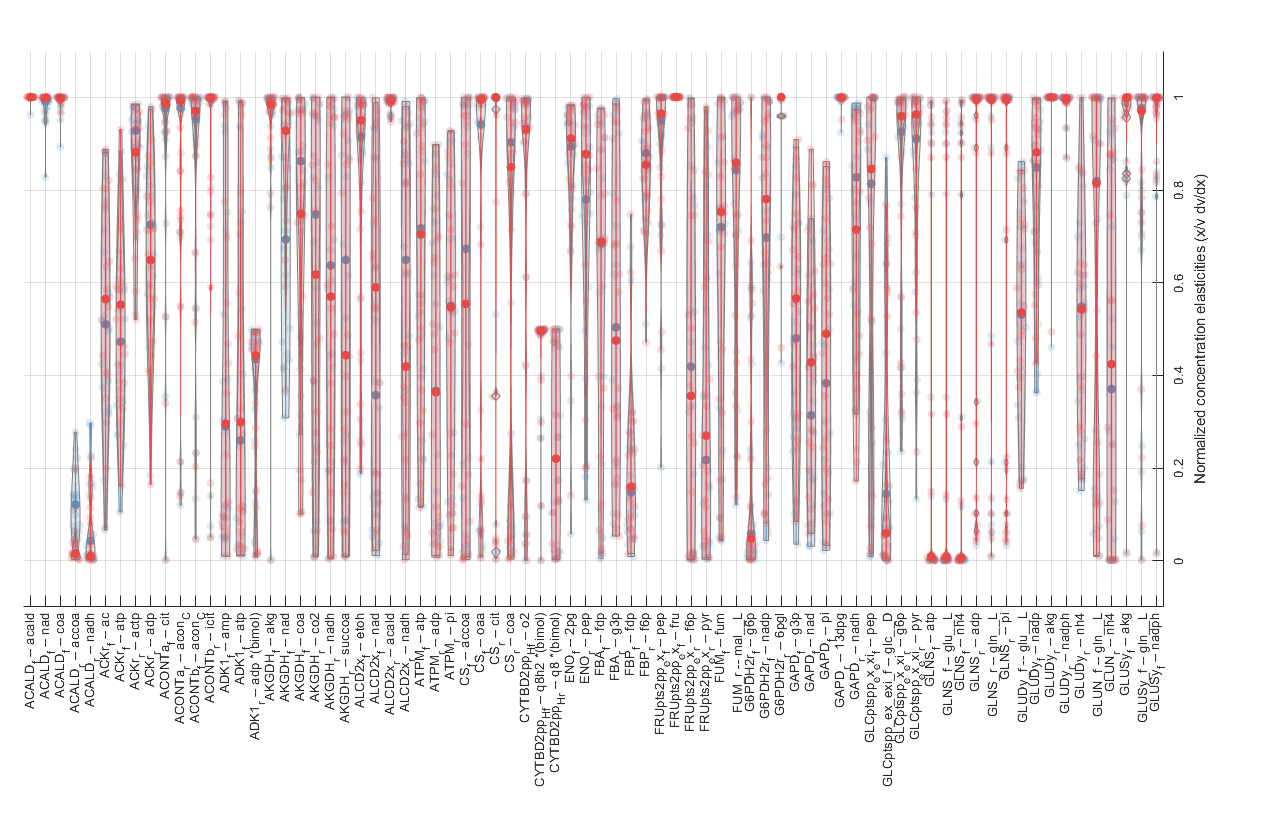

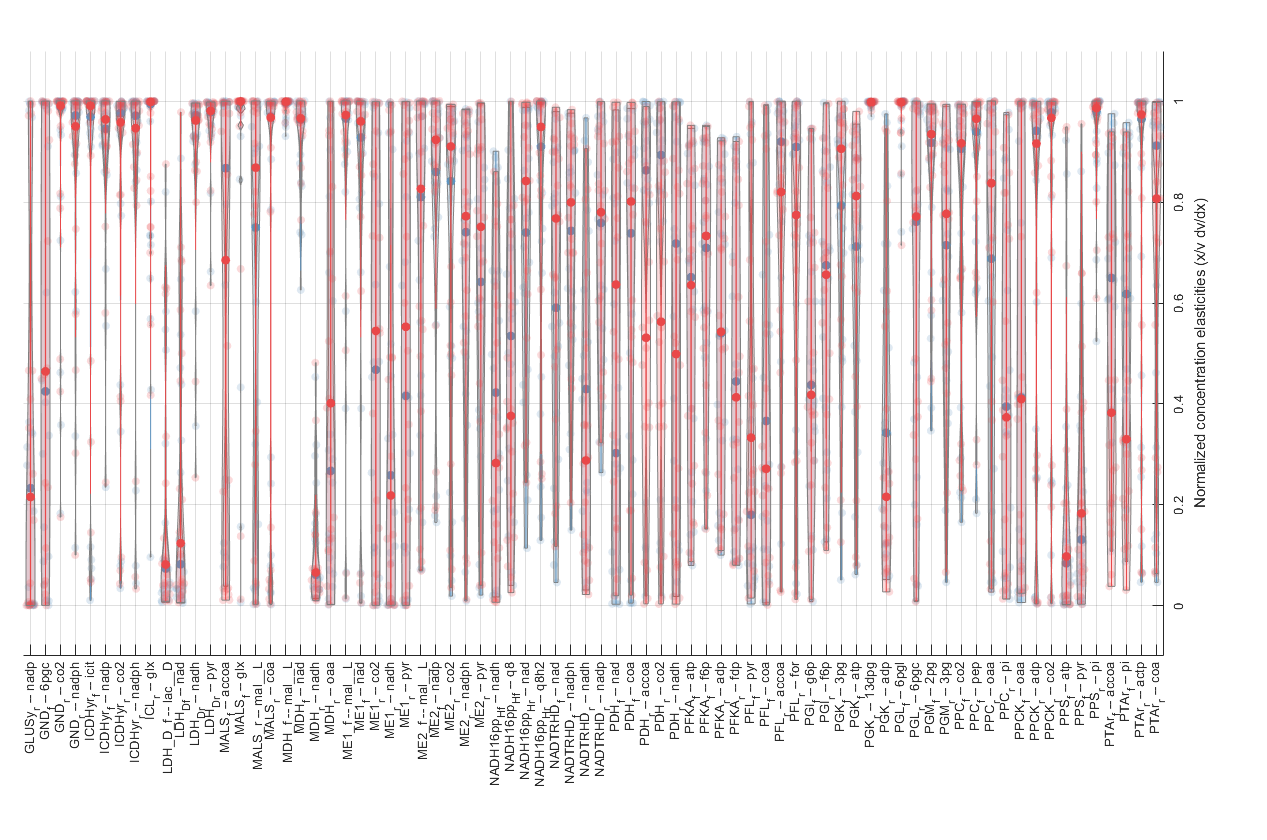

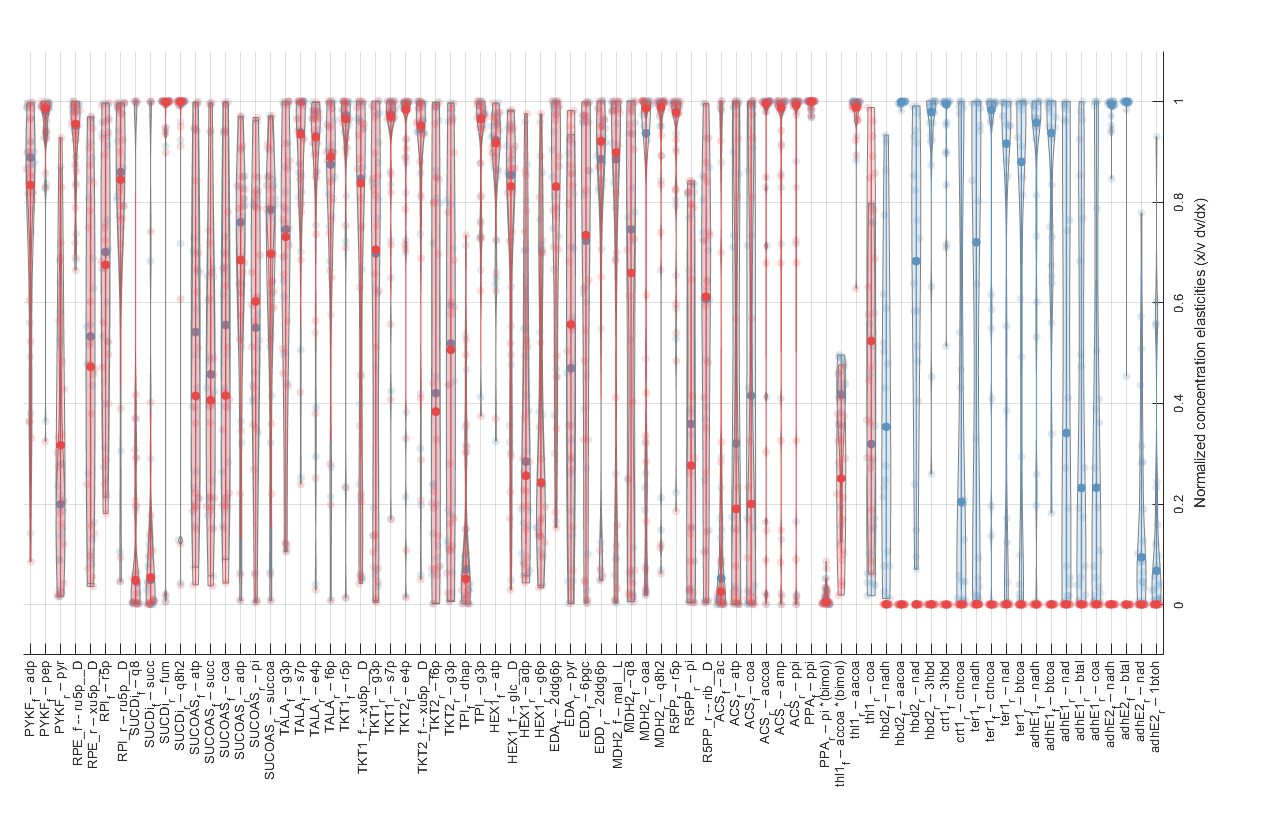

**(b)**
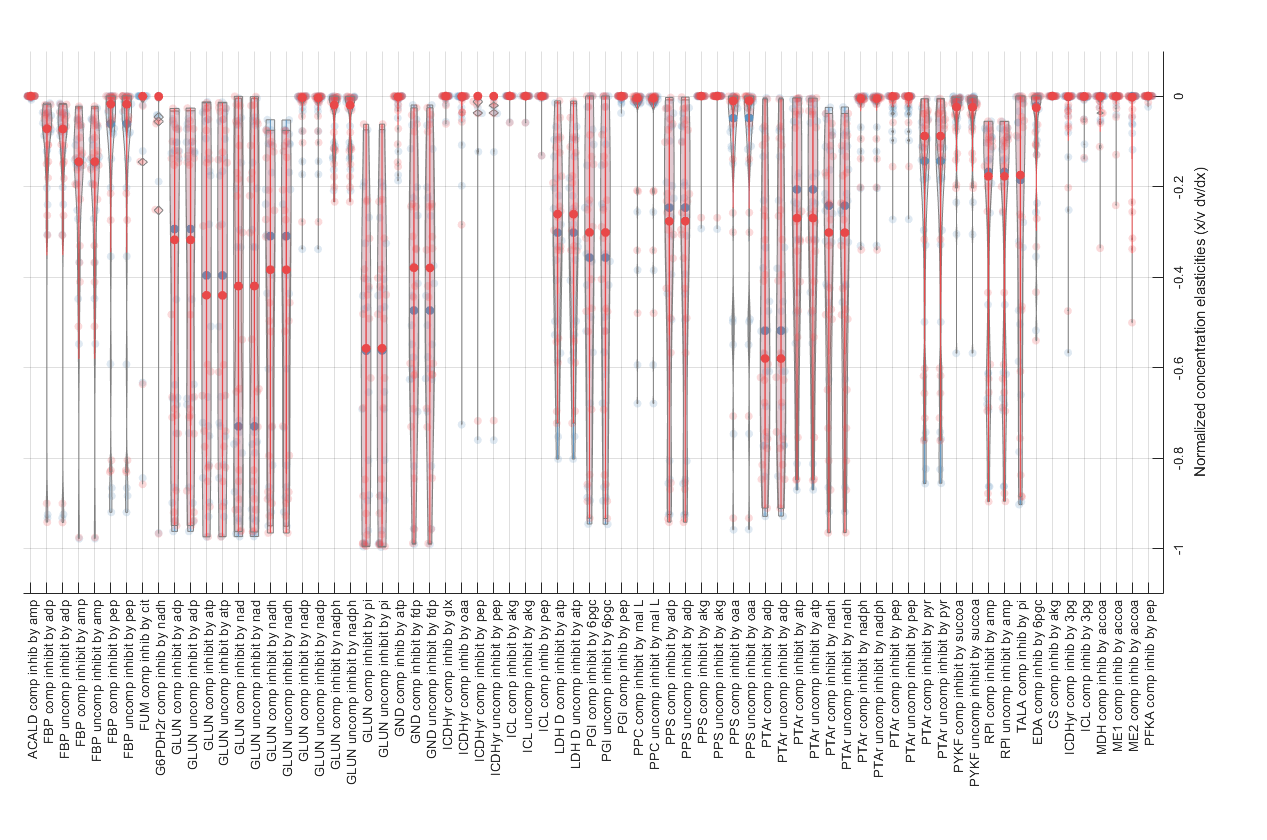

**(c)**
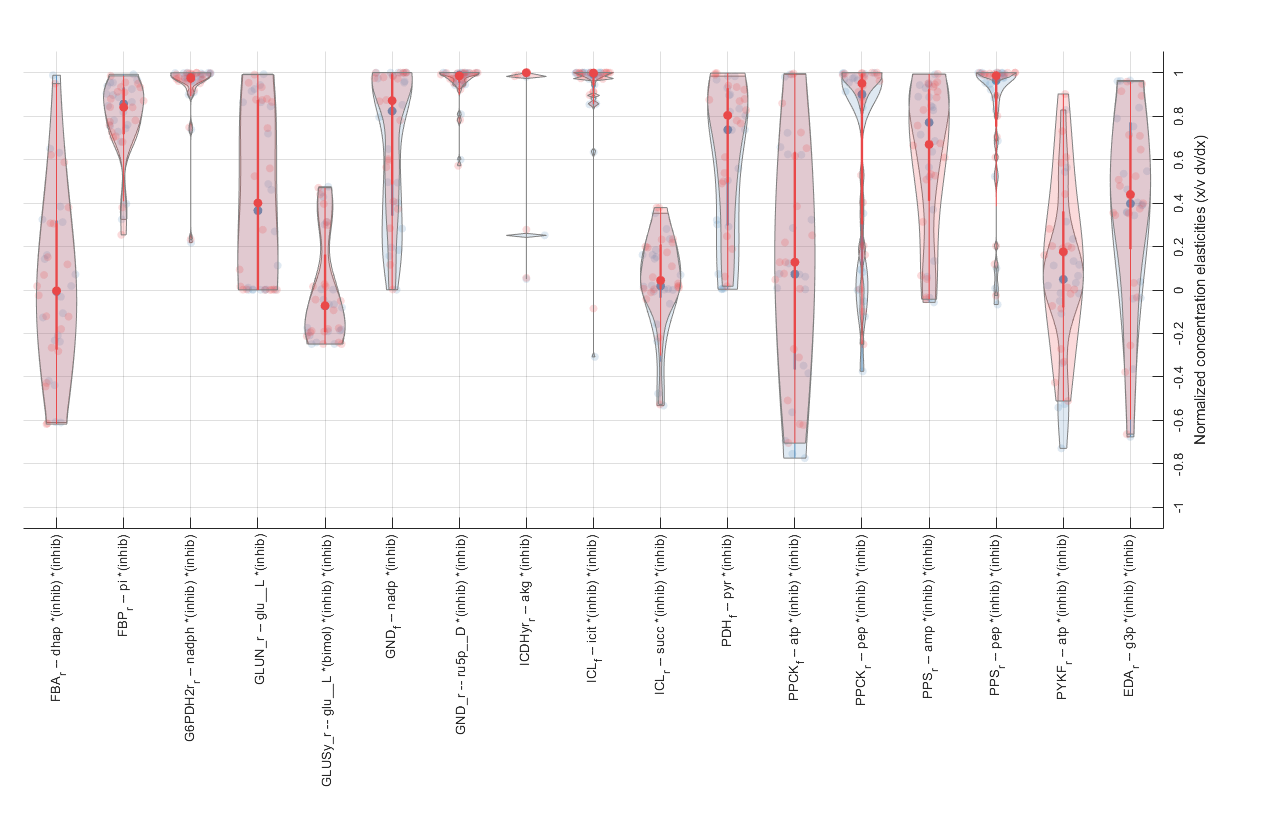

**Supplementary Figure S12: Model-wide concentration elasticities.** Concentration elasticity values for (a) all substrate-reaction pairs (as proxies for each dissociation constant) (b) inhibitor-reaction pairs are given, as well as metabolites that act as both a substrate and an inhibitor in the same reaction (c). Violins show the distribution across the final 20 models in the ensemble for both the butanol-positive (blue) and butanol-negative (red) conditions, and dark circles indicate median values across models.

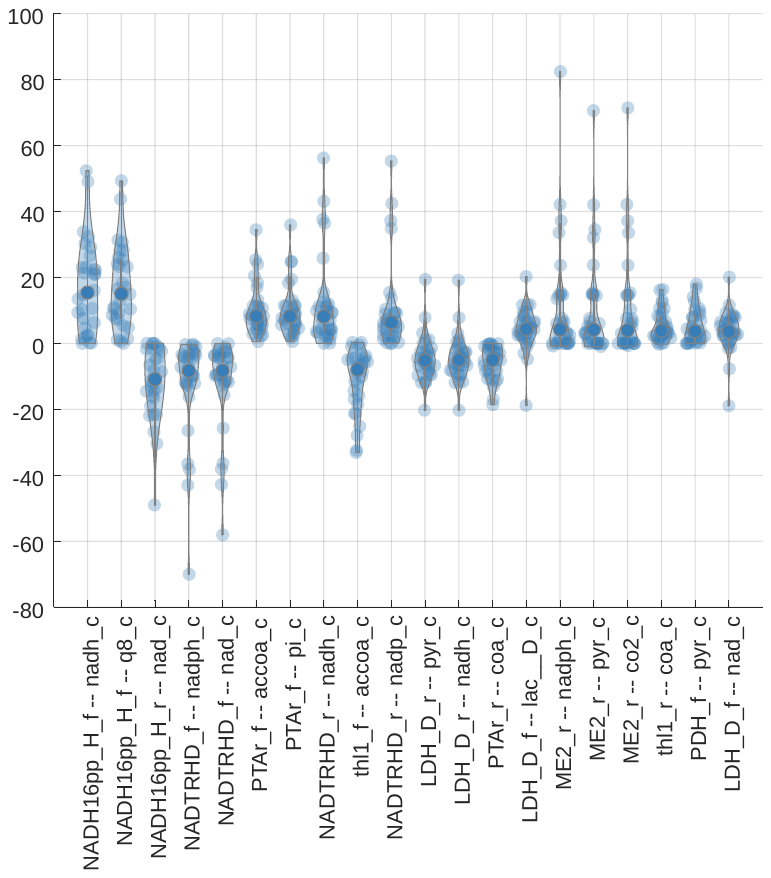

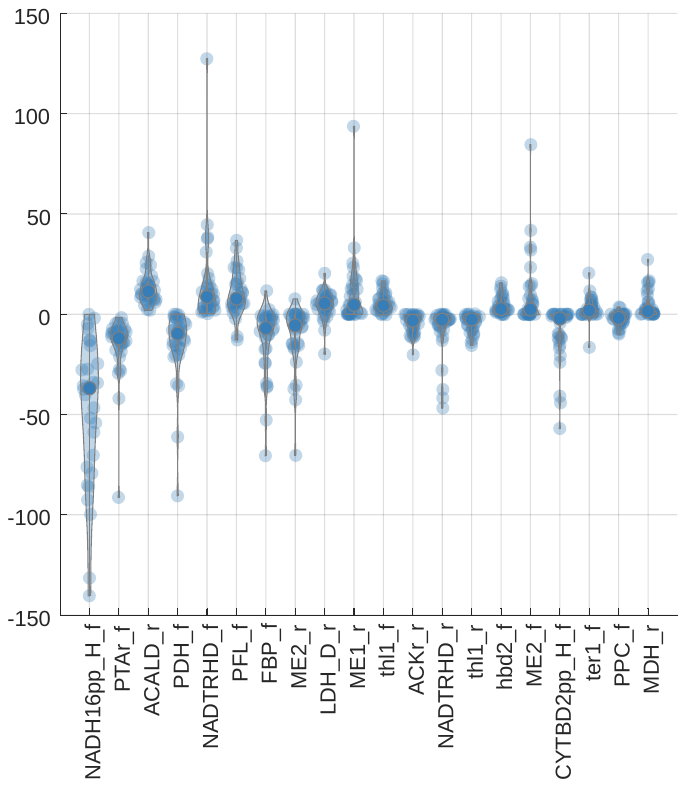
**(a)**

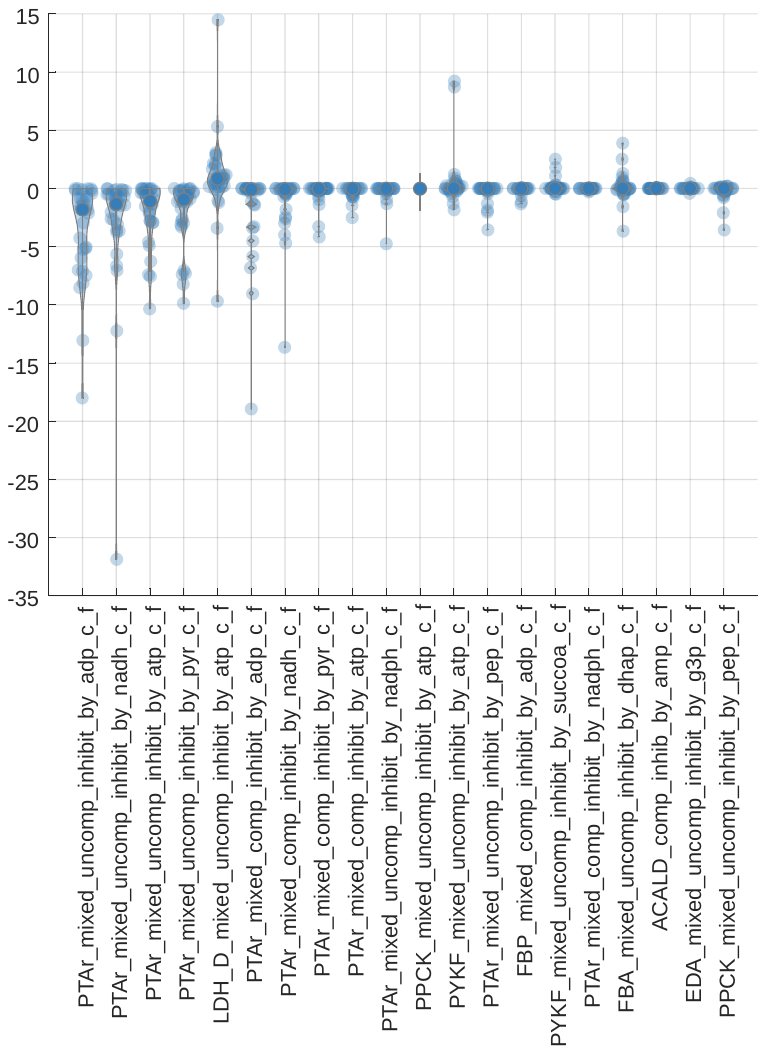

**(b)**
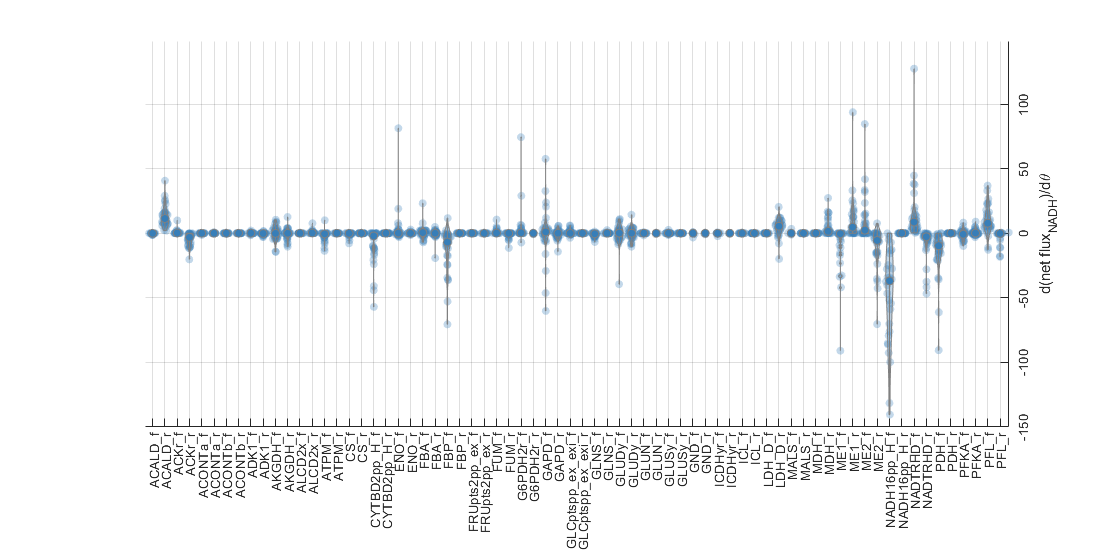

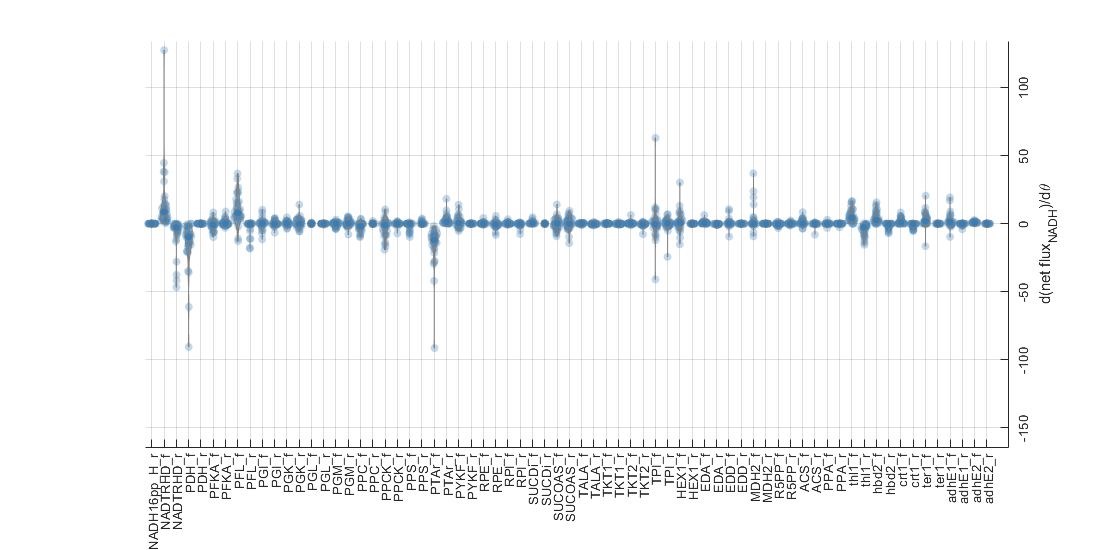

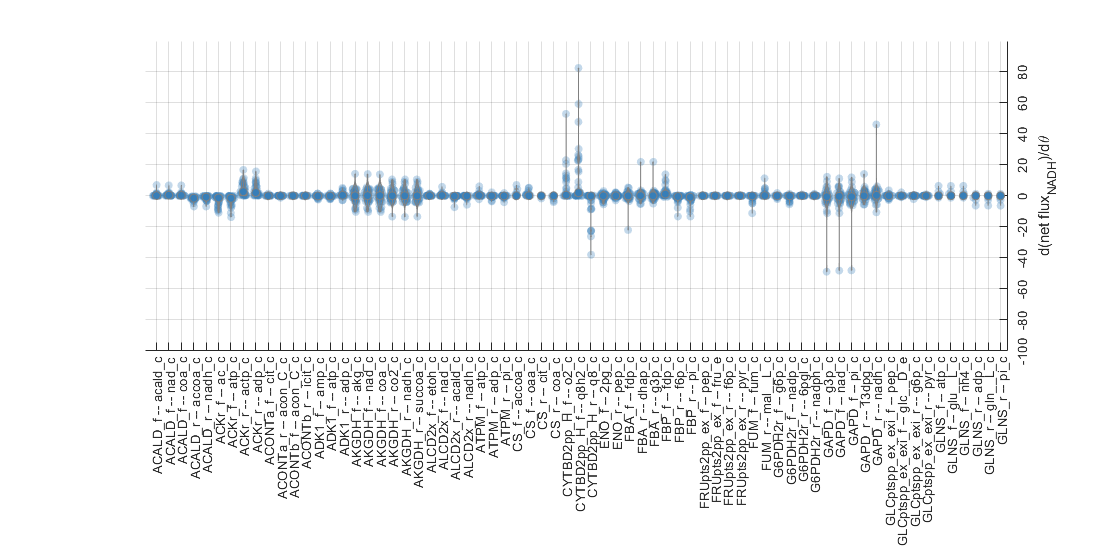

**(c)**
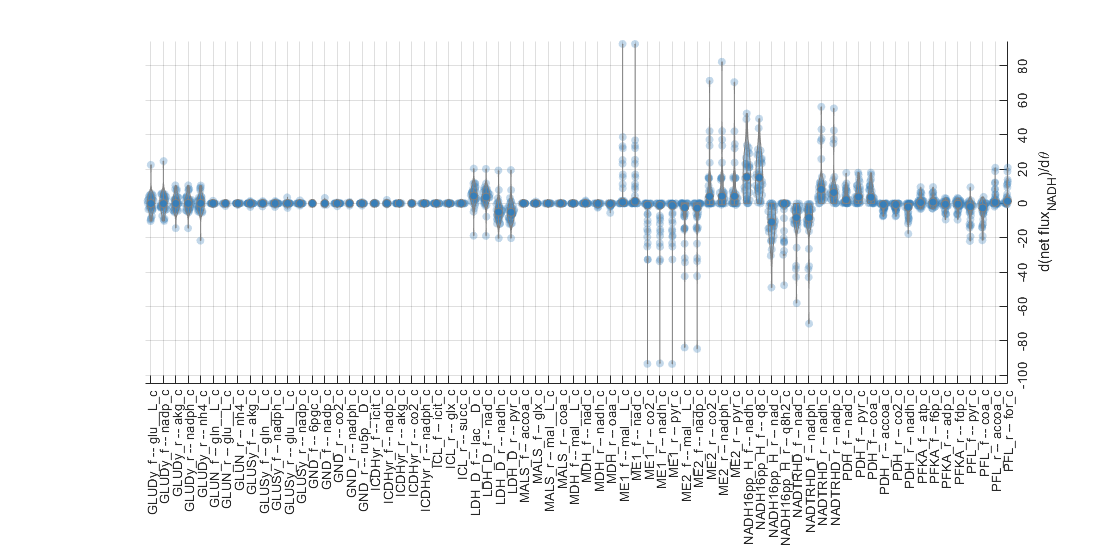

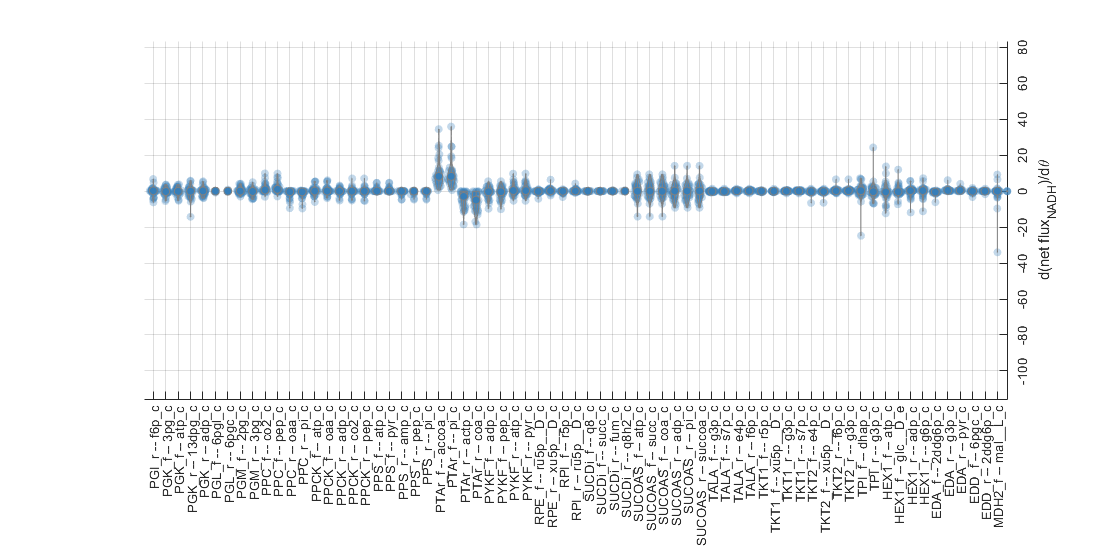

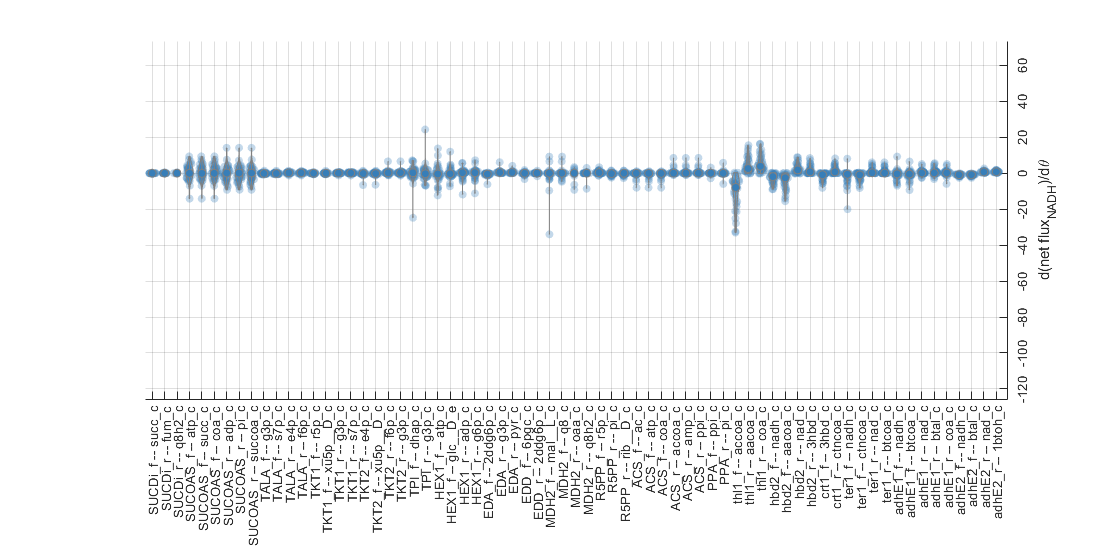

**(d)**
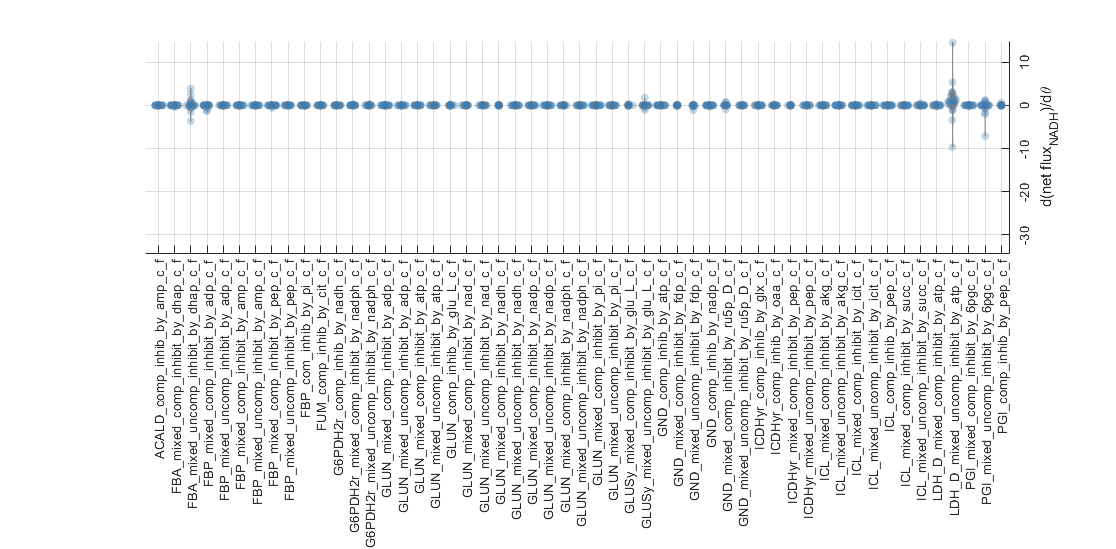

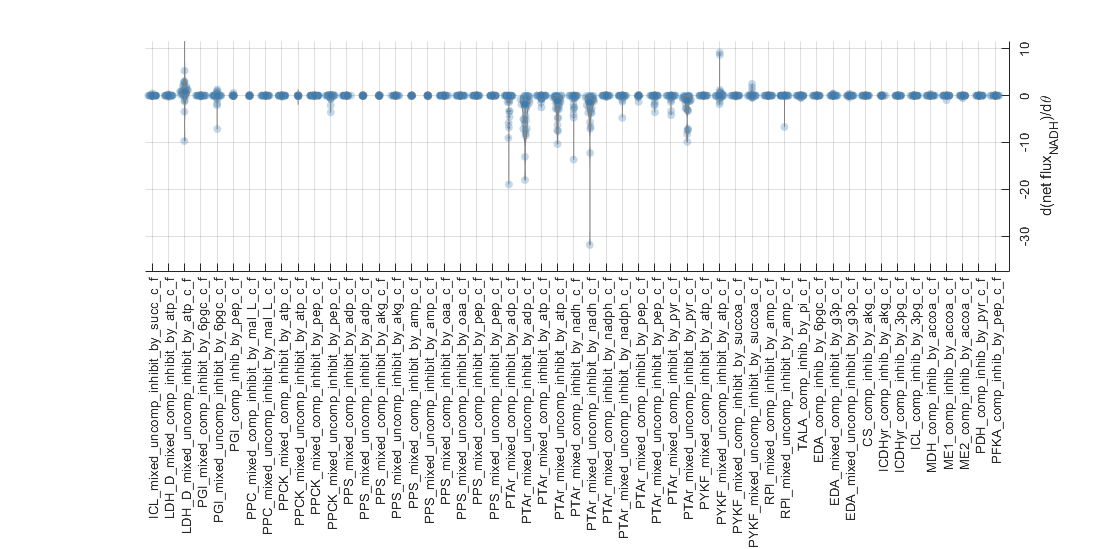

**Supplementary Figure S13: Sensitivity of parameters to NAD flux.** The sensitivities of parameters with respect to cumulative model NAD regeneration are shown. (a) The top 20 of each group of rate constants, dissociation constants, and inhibition constants are shown, as ranked by the median absolute value of sensitivity across all 20 parameter sets. Sensitivities for all parameters all also shown, and are again split by (b) rate constants, (c) saturation/dissociation constants, and (d) inhibition constants.

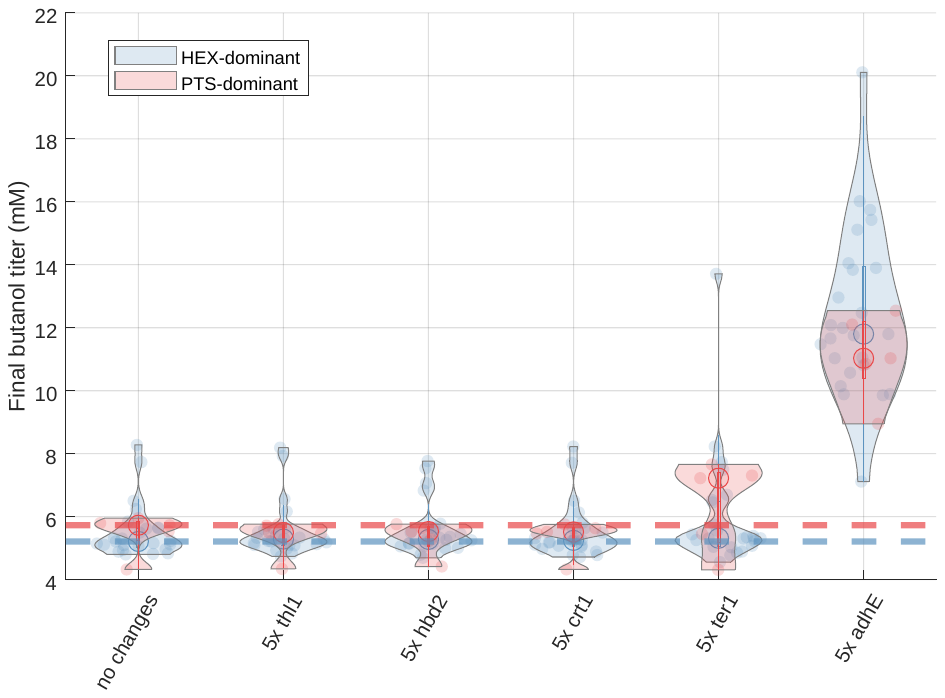

**Supplementary Figure S14: Comparison of butanol predictions between models with differing glucose-consumption behaviors.** Models are binned into those which carry more flux through hexokinase (HEX) than through glucose-phosphotransferase system (PTS), shown in blue, or those which carry more flux through PTS than HEX, shown in red. Despite these differences, these populations give qualitatively similar predictions regarding the 20-hour butanol titer from the overexpression of each of the five enzymes in the butanol pathway (baseline levels of butanol titer for each population are shown as a dotted line).
