## Supplementary material for "Dynamic Kinetic Models Capture Cell-Free Metabolism for Improved Butanol Production": cell_free_kinetic_model-Supplementary_Methods_Final.docx

Running Title:

Cell-Free Dynamic Model

Jacob Martin^1,2,3^, Blake Rasor^1,2,3^, Jon DeBonis^1^, Ashty Karim^1,2,3^, Linda J. Broadbelt^1,2^*, Keith E.J. Tyo^1,2,3^*

^1^Department of Chemical and Biological Engineering, Northwestern University, Evanston, IL, USA 60208

^2^Center for Synthetic Biology, Northwestern University, Evanston, IL, USA 60208

^3^Chemistry of Life Processes Institute, Northwestern University, Evanston, IL, USA 60208

Manuscript in preparation for:

Metabolic Engineering Journal

*Corresponding author:

### S1: Normalization of parameters and metabolites against reference state

For reaction *j* with an arbitrary numbers of substrates & products:

$$A+B\to P+Q$$

a general rate equation for of the form

$$v_{net,j}=\frac{V_{max,f}\frac{\left[ A \right]}{K_{mA}}\frac{\left[ B \right]}{K_{mB}}-V_{max,r}\frac{\left[ P \right]}{K_{mP}}\frac{\left[ Q \right]}{K_{mQ}}}{1+\frac{\left[ A \right]}{K_{mA}}+\frac{\left[ B \right]}{K_{mB}}+\frac{\left[ P \right]}{K_{mP}}+\frac{\left[ Q \right]}{K_{mQ}}+\{additional terms\}}$$

can be given. The vector of all rates $\boldsymbol{v}_{\boldsymbol{net}}$ is used to calculate the system of ordinary differential equations, so that the vector of time-derivatives of all metabolite concentrations, $\boldsymbol{x=}\left\{ \left[ A \right],\left[ B \right],\left[ P \right],\left[ Q \right],\ldots\right\}$, is given by

$$\frac{d\boldsymbol{x}}{dt}=\boldsymbol{S}\boldsymbol{v}_{\boldsymbol{net}}$$

where $\boldsymbol{S}$ is the stoichiometric matrix.

If we divide each metabolite concentration or Michaelis-type constant by a fixed reference concentration of each metabolite $\boldsymbol{x}_{\boldsymbol{ref}}=\left\{ \left[ A_{ref} \right],\left[ B_{ref} \right],\left[ P_{ref} \right],\left[ Q_{ref} \right],\ldots\right\}$:

$$v_{net,j}=\frac{V_{max,f}\frac{\left[ A \right]/[A_{ref}]}{K_{mA}/[A_{ref}]}\frac{\left[ B \right]/\left[ B_{ref} \right]}{K_{mB}/[B_{ref}]}-V_{max,r}\frac{\left[ P \right]/[P_{ref}]}{K_{mP}/[P_{ref}]}\frac{\left[ Q \right]/[Q_{ref}]}{K_{mQ}/[Q_{ref}]}}{1+\frac{\left[ A \right]/[A_{ref}]}{K_{mA}/[A_{ref}]}+\frac{\left[ B \right]/[B_{ref}]}{K_{mB}/[B_{ref}]}+\frac{\left[ P \right]/[P_{ref}]}{K_{mP}/[P_{ref}]}+\frac{\left[ Q \right]/[Q_{ref}]}{K_{mQ}/[Q_{ref}]}+\{additional terms\}}$$

we get the similar rate form:

$$v_{net,j}=\frac{V_{max,f}\frac{\tilde{\left[ A \right]}}{\tilde{K_{mA}}}\frac{\tilde{\left[ B \right]}}{\tilde{K_{mB}}}-V_{max,r}\frac{\tilde{\left[ P \right]}}{\tilde{K_{mP}}}\frac{\tilde{\left[ Q \right]}}{\tilde{K_{mQ}}}}{1+\frac{\tilde{\left[ A \right]}}{\tilde{K_{mA}}}+\frac{\tilde{\left[ B \right]}}{\tilde{K_{mB}}}+\frac{\tilde{\left[ P \right]}}{\tilde{K_{mP}}}+\frac{\tilde{\left[ Q \right]}}{\tilde{K_{mQ}}}+\{additional terms\}}$$

where $\tilde{\boldsymbol{x}}=\frac{\boldsymbol{x}}{\boldsymbol{x}_{\boldsymbol{ref}}}\boldsymbol{=}\left\{ \tilde{\left[ A \right]},\tilde{\left[ B \right]},\tilde{\left[ P \right]},\tilde{\left[ Q \right]},\ldots\right\}$ are now relative metabolite concentrations

If the steady-state concentrations $\boldsymbol{x}_{\boldsymbol{ref}}$ are taken to be the concentrations at a “reference state” at which steady-state fluxes were measured, then at that reference state, the normalized concentrations of all species are 1 at that steady-state. This then reduces the rate equation to:

$$v_{net,j}=\frac{V_{max,f}\frac{1}{\tilde{K_{mA}}}\frac{1}{\tilde{K_{mB}}}-V_{max,r}\frac{1}{\tilde{K_{mP}}}\frac{1}{\tilde{K_{mQ}}}}{1+\frac{1}{\tilde{K_{mA}}}+\frac{1}{\tilde{K_{mB}}}+\frac{1}{\tilde{K_{mP}}}+\frac{1}{\tilde{K_{mQ}}}+\{additional maybe\}}$$

where $v_{net,j}$ is known and each $V_{max}$ and $\tilde{K_{m}}$ can be calculated or sampled with fewer degrees of freedom.

The system of ODEs being solved is now equivalent to:

$$\frac{d\tilde{\boldsymbol{x}}}{dt}\boldsymbol{=}\frac{1}{\boldsymbol{x}_{\boldsymbol{ref}}}\frac{d\boldsymbol{x}}{dt}=\frac{1}{\boldsymbol{x}_{\boldsymbol{ref}}}\boldsymbol{Sv}$$

However, because $\boldsymbol{x}_{\boldsymbol{ref}}$ is not typically known, the above equation is only correct (using $x_{ref,i}\boldsymbol{=}1$**)** when $\frac{d\tilde{\boldsymbol{x}}}{dt}\boldsymbol{=}\frac{d\boldsymbol{x}}{dt}=0$, i.e., at the steady-state solution. Therefore, this method of normalization requires a known steady-state solution to use as a reference state and is only strictly accurate at the steady-state solution, and should not be used to capture transient behavior.

### S2: Derivation of Enzyme Rate Law

The derivation of this rate equation is similar to that described by Cornish-Bowden for reactions with multiple substrates (Cornish-Bowden, 1979), but differs in that we assume that the order of substrate binding does not change a given dissociation constant.

Consider a general reversible enzymatic reaction,

$$A+B{E \atop\begin{aligned} \rightleftharpoons\\ \end{aligned}}P+Q$$

Defined by elementary binding steps and rate constants as the following five reversible steps:

$$A+E{k_{Af} \atop\begin{aligned} \rightleftharpoons\\ k_{Ar} \end{aligned}}EA ; EA+B{k_{Bf} \atop\begin{aligned} \rightleftharpoons\\ k_{Br} \end{aligned}}EAB ; EAB{k_{cat,f} \atop\begin{aligned} \rightleftharpoons\\ k_{cat,r} \end{aligned}}EPQ ; EPQ{k_{Qr} \atop\begin{aligned} \rightleftharpoons\\ k_{Qf} \end{aligned}}EP+Q ; EP{k_{Pr} \atop\begin{aligned} \rightleftharpoons\\ k_{Pf} \end{aligned}}P+E$$

If we assume that substrates or products can bind to the enzyme in either order, and that binding order does not affect the dissociation constant of that binding event, then both of the following are true:

$$A+E{k_{Af} \atop\begin{aligned} \rightleftharpoons\\ k_{Ar} \end{aligned}}EA, K_{dA}=\frac{\left[ A \right]\left[ E \right]}{\left[ EA \right]}=\frac{k_{Ar}}{k_{Af}}$$

$$A+EB{k_{Af} \atop\begin{aligned} \rightleftharpoons\\ k_{Ar} \end{aligned}}EAB, K_{dA}=\frac{\left[ A \right]\left[ EB \right]}{[EAB]}=\frac{k_{Ar}}{k_{Af}}$$

for substrate *A*, or

$$B+E{k_{Bf} \atop\begin{aligned} \rightleftharpoons\\ k_{Br} \end{aligned}}EB, K_{dB}=\frac{\left[ B \right]\left[ E \right]}{\left[ EB \right]}=\frac{k_{Br}}{k_{Bf}}$$

$$B+EA{k_{Bf} \atop\begin{aligned} \rightleftharpoons\\ k_{Br} \end{aligned}}EAB, K_{dB}=\frac{\left[ B \right]\left[ EA \right]}{[EAB]}=\frac{k_{Br}}{k_{Bf}}$$

for substrate *B* (and likewise for products). The amount of free enzyme available for binding is given by:

$$E=E_{free}=E_{tot}-\left[ EA \right]-\left[ EB \right]-\left[ EAB \right]-\left[ EP \right]-\left[ EQ \right]-\left[ EPQ \right]$$

By substituting equations, the equation becomes:

$$[E_{free}]=\left[ E_{tot} \right]-\frac{\left[ A \right]\left[ E \right]}{K_{dA}}-\frac{\left[ B \right]\left[ E \right]}{K_{dB}}-\frac{\left[ A \right]}{K_{dA}}\left( \frac{\left[ B \right]\left[ E \right]}{K_{dB}} \right)-\frac{\left[ P \right]\left[ E \right]}{K_{dP}}-\frac{\left[ Q \right]\left[ E \right]}{K_{dQ}}-\frac{\left[ P \right]}{K_{dP}}\left( \frac{\left[ Q \right]\left[ E \right]}{K_{dQ}} \right)$$

$$\left[ E_{free} \right]=\left[ E_{tot} \right]-\left[ E \right]\left( \frac{\left[ A \right]}{K_{dA}}+\frac{\left[ B \right]}{K_{dB}}+\frac{\left[ A \right]\left[ B \right]}{K_{dA}K_{dB}}+ \frac{\left[ P \right]}{K_{dP}}+\frac{\left[ Q \right]}{K_{dQ}}+\frac{\left[ P \right]\left[ Q \right]}{K_{dP}K_{dQ}} \right)$$

$$\left[ E_{free} \right]=\frac{E_{tot}}{1+\frac{\left[ A \right]}{K_{dA}}+\frac{\left[ B \right]}{K_{dB}}+\frac{\left[ A \right]\left[ B \right]}{K_{dA}K_{dB}}+ \frac{\left[ P \right]}{K_{dP}}+\frac{\left[ Q \right]}{K_{dQ}}+\frac{\left[ P \right]\left[ Q \right]}{K_{dP}K_{dQ}}}$$

Assuming that all binding steps are quasi-equilibrated and so the net rate of the reaction (in each direction) to be controlled by the catalytic step, the rate – as defined by mass action –is the product of the ultimate complex and the catalytic rate constant:

$$EAB {k_{cat,f} \atop\begin{aligned} \rightleftharpoons\\ k_{cat,r} \end{aligned}}ESP, v_{f}=k_{cat,f}\left[ EAB \right], v_{r}=k_{cat,r}[ESP]$$

Solving for the concentration of the final complex of the enzyme and all substrates,

$$EAB=\frac{\left[ A \right]\left[ EB \right]}{K_{dA}}=\frac{\left[ A \right]}{K_{da}}\left( \frac{\left[ B \right]\left[ E_{free} \right]}{K_{dB}} \right)$$

$$=\frac{\frac{\left[ A \right]\left[ B \right]\left[ E_{tot} \right]}{K_{dA}K_{dB}}}{1+\frac{\left[ A \right]}{K_{dA}}+\frac{\left[ B \right]}{K_{dB}}+\frac{\left[ A \right]\left[ B \right]}{K_{dA}K_{dB}}+ \frac{\left[ P \right]}{K_{dP}}+\frac{\left[ Q \right]}{K_{dQ}}+\frac{\left[ P \right]\left[ Q \right]}{K_{dP}K_{dQ}}}$$

And finally, substituting this into the forward and reverse rates, the final rate equations are given:

$$v_{f}=\frac{\frac{k_{cat,f}\left[ E_{tot} \right]\left[ A \right]\left[ B \right]}{K_{dA}K_{dB}}}{1+\frac{\left[ A \right]}{K_{dA}}+\frac{\left[ B \right]}{K_{dB}}+\frac{\left[ A \right]\left[ B \right]}{K_{dA}K_{dB}}+ \frac{\left[ P \right]}{K_{dP}}+\frac{\left[ Q \right]}{K_{dQ}}+\frac{\left[ P \right]\left[ Q \right]}{K_{dP}K_{dQ}}}$$

$$v_{r}=\frac{\frac{k_{cat,r}\left[ E_{tot} \right]\left[ P \right]\left[ Q \right]}{K_{dP}K_{dQ}}}{1+\frac{\left[ A \right]}{K_{dA}}+\frac{\left[ B \right]}{K_{dB}}+\frac{\left[ A \right]\left[ B \right]}{K_{dA}K_{dB}}+ \frac{\left[ P \right]}{K_{dP}}+\frac{\left[ Q \right]}{K_{dQ}}+\frac{\left[ P \right]\left[ Q \right]}{K_{dP}K_{dQ}}}$$

We here repeat the derivation for a single substrate (or product) to show that this form is generalizable to an arbitrary number of substrates and products. For the reaction:

$$A{E \atop\begin{aligned} \rightleftharpoons\\ \end{aligned}}P+Q$$

elementary binding steps and rate constants are given by these four reversible steps:

$$A+E{k_{Af} \atop\begin{aligned} \rightleftharpoons\\ k_{Ar} \end{aligned}}EA ; EA{k_{cat,f} \atop\begin{aligned} \rightleftharpoons\\ k_{cat,r} \end{aligned}}EPQ ; EPQ{k_{Qr} \atop\begin{aligned} \rightleftharpoons\\ k_{Qf} \end{aligned}}EP+Q ; EP{k_{Pr} \atop\begin{aligned} \rightleftharpoons\\ k_{Pf} \end{aligned}}P+E$$

If we assume that substrates or products can bind to the enzyme in either order, and that binding order does not affect the dissociation constant of that binding event, then I can have either:

$$A+E{k_{Af} \atop\begin{aligned} \rightleftharpoons\\ k_{Ar} \end{aligned}}EA, K_{dA}=\frac{\left[ A \right]\left[ E \right]}{\left[ EA \right]}=\frac{k_{Ar}}{k_{Af}}$$

for substrate *A*, or

$$P+E{k_{Pf} \atop\begin{aligned} \rightleftharpoons\\ k_{Pr} \end{aligned}}EP, K_{dP}=\frac{\left[ P \right]\left[ E \right]}{\left[ EP \right]}=\frac{k_{Pr}}{k_{Pf}}$$

$$P+EQ{k_{Pf} \atop\begin{aligned} \rightleftharpoons\\ k_{Pr} \end{aligned}}EPQ, K_{dP}=\frac{\left[ P \right]\left[ EQ \right]}{[EPQ]}=\frac{k_{Pr}}{k_{Pf}}$$

for product *P*. The amount of free enzyme available for binding is given by:

$$E=E_{free}=E_{tot}-\left[ EA \right]-\left[ EP \right]-\left[ EQ \right]-\left[ EPQ \right]$$

By substituting equations, the equation becomes:

$$[E_{free}]=\left[ E_{tot} \right]-\frac{\left[ A \right]\left[ E \right]}{K_{dA}}-\frac{\left[ P \right]\left[ E \right]}{K_{dP}}-\frac{\left[ Q \right]\left[ E \right]}{K_{dQ}}-\frac{\left[ P \right]}{K_{dP}}\left( \frac{\left[ Q \right]\left[ E \right]}{K_{dQ}} \right)$$

$$\left[ E_{free} \right]=\left[ E_{tot} \right]-\left[ E \right]\left( \frac{\left[ A \right]}{K_{dA}}+ \frac{\left[ P \right]}{K_{dP}}+\frac{\left[ Q \right]}{K_{dQ}}+\frac{\left[ P \right]\left[ Q \right]}{K_{dP}K_{dQ}} \right)$$

$$\left[ E_{free} \right]=\frac{E_{tot}}{1+\frac{\left[ A \right]}{K_{dA}}+ \frac{\left[ P \right]}{K_{dP}}+\frac{\left[ Q \right]}{K_{dQ}}+\frac{\left[ P \right]\left[ Q \right]}{K_{dP}K_{dQ}}}$$

Taking the net rate of the reaction (in each direction) to be controlled by the catalytic step (i.e., all enzyme binding or dissociation steps are at quasi-equilibrium), the rate – as defined by mass action – will be the product of the ultimate complex and the catalytic rate constant:

$$EA {k_{cat,f} \atop\begin{aligned} \rightleftharpoons\\ k_{cat,r} \end{aligned}}ESP, v_{f}=k_{cat,f}\left[ EA \right], v_{r}=k_{cat,r}[ESP]$$

Solving for the concentration of the final complex of the enzyme and all substrates,

$$EA=\frac{\left[ A \right]\left[ E_{free} \right]}{K_{dA}}$$

$$=\frac{\frac{\left[ A \right]\left[ E_{tot} \right]}{K_{dA}}}{1+\frac{\left[ A \right]}{K_{dA}}+ \frac{\left[ P \right]}{K_{dP}}+\frac{\left[ Q \right]}{K_{dQ}}+\frac{\left[ P \right]\left[ Q \right]}{K_{dP}K_{dQ}}}$$

And finally, substituting this into the forward and reverse rates, the final rate equations are given:

$$v_{f}=\frac{\frac{k_{cat,f}\left[ E_{tot} \right]\left[ A \right]}{K_{dA}}}{1+\frac{\left[ A \right]}{K_{dA}}+\frac{\left[ B \right]}{K_{dB}}+\frac{\left[ A \right]\left[ B \right]}{K_{dA}K_{dB}}+ \frac{\left[ P \right]}{K_{dP}}+\frac{\left[ Q \right]}{K_{dQ}}+\frac{\left[ P \right]\left[ Q \right]}{K_{dP}K_{dQ}}}$$

$$v_{r}=\frac{\frac{k_{cat,r}\left[ E_{tot} \right]\left[ P \right]\left[ Q \right]}{K_{dP}K_{dQ}}}{1+\frac{\left[ A \right]}{K_{dA}}+ \frac{\left[ P \right]}{K_{dP}}+\frac{\left[ Q \right]}{K_{dQ}}+\frac{\left[ P \right]\left[ Q \right]}{K_{dP}K_{dQ}}}$$

Inhibition can be included as either competitive, wherein the free enzyme is bound by a non-reactive inhibitor:

$$I_{c}+E{k_{Ic,f} \atop\begin{aligned} \rightleftharpoons\\ k_{Ic,r} \end{aligned}}EI_{c}$$

$$K_{Ic}=\frac{\left[ E \right]\left[ I_{c} \right]}{[EI_{c}]}=\frac{k_{Ic,r}}{k_{Ic,f}}$$

or noncompetitive, where the fully-bound ternary complex is bound and made non-reactive:

$$I_{u}+EAB{k_{Iu,f} \atop\begin{aligned} \rightleftharpoons\\ k_{Iu,r} \end{aligned}}EABI_{u}$$

$$K_{Iu}=\frac{\left[ EAB \right]\left[ I_{u} \right]}{[EABI_{u}]}=\frac{k_{Iu,r}}{k_{Iu,f}}$$

For competitive inhibition, the free enzyme term is altered:

$$E=E_{free}=E_{tot}-\left[ EA \right]-\left[ EP \right]-\left[ EQ \right]-\left[ EPQ \right]-[EI_{c}]$$

Again assuming quasi-equlibrium for the inhibitor and substituting the equilibrium expression, the equation becomes:

$$\left[ E_{free} \right]=\left[ E_{tot} \right]-\left[ E \right]\left( \frac{\left[ A \right]}{K_{dA}}+ \frac{\left[ P \right]}{K_{dP}}+\frac{\left[ Q \right]}{K_{dQ}}+\frac{\left[ P \right]\left[ Q \right]}{K_{dP}K_{dQ}}+\frac{\left[ I_{c} \right]}{K_{Ic}} \right)$$

$$\left[ E_{free} \right]=\frac{E_{tot}}{1+\frac{\left[ A \right]}{K_{dA}}+ \frac{\left[ P \right]}{K_{dP}}+\frac{\left[ Q \right]}{K_{dQ}}+\frac{\left[ P \right]\left[ Q \right]}{K_{dP}K_{dQ}}+\frac{\left[ I_{c} \right]}{K_{Ic}}}$$

which gives the final rates:

$$v_{f}=\frac{\frac{k_{cat,f}\left[ E_{tot} \right]\left[ A \right]\left[ B \right]}{K_{dA}K_{dB}}}{1+\frac{\left[ A \right]}{K_{dA}}+\frac{\left[ B \right]}{K_{dB}}+\frac{\left[ A \right]\left[ B \right]}{K_{dA}K_{dB}}+ \frac{\left[ P \right]}{K_{dP}}+\frac{\left[ Q \right]}{K_{dQ}}+\frac{\left[ P \right]\left[ Q \right]}{K_{dP}K_{dQ}}+\frac{\left[ I_{c} \right]}{K_{Ic}}}$$

$$v_{r}=\frac{\frac{k_{cat,r}\left[ E_{tot} \right]\left[ P \right]\left[ Q \right]}{K_{dP}K_{dQ}}}{1+\frac{\left[ A \right]}{K_{dA}}+\frac{\left[ B \right]}{K_{dB}}+\frac{\left[ A \right]\left[ B \right]}{K_{dA}K_{dB}}+ \frac{\left[ P \right]}{K_{dP}}+\frac{\left[ Q \right]}{K_{dQ}}+\frac{\left[ P \right]\left[ Q \right]}{K_{dP}K_{dQ}}+\frac{\left[ I_{c} \right]}{K_{Ic}}}$$

For uncompetitive inhibition, assuming the uncompetitive inhibitor can bind the active complex of either the substrate or the product (and assuming that the inhibition dissociation constant is the same for both), the amount of free enzyme is now lessened by:

$$E_{free}=E_{tot}-\left[ EA \right]-\left[ EB \right]-[EAB]-\left[ EP \right]-\left[ EQ \right]-\left[ EPQ \right]-[EABI_{u}]-[EPQI_{u}]$$

Rearranging Eq. <X.X >, we get

$$\left[ EABI_{u} \right]=\frac{\left[ EAB \right]\left[ I_{u} \right]}{K_{Iu}}, \left[ EPQI_{u} \right]=\frac{\left[ EPQ \right]\left[ I_{u} \right]}{K_{Iu}}$$

which substituting into Eq. <X.X> gives

$$E_{free}=E_{tot}-\left[ EA \right]-\left[ EB \right]-\left[ EAB \right]\left( 1+\frac{\left[ I_{u} \right]}{K_{iu}} \right)-\left[ EP \right]-\left[ EQ \right]-\left[ EPQ \right]\left( 1+\frac{\left[ I_{u} \right]}{K_{iu}} \right)$$

By following the same steps above, this leads to the final rates:

$$v_{f}=\frac{\frac{k_{cat,f}\left[ E_{tot} \right]\left[ A \right]\left[ B \right]}{K_{dA}K_{dB}}}{1+\frac{\left[ A \right]}{K_{dA}}+\frac{\left[ B \right]}{K_{dB}}+\frac{\left[ A \right]\left[ B \right]}{K_{dA}K_{dB}}\left( 1+\frac{\left[ I_{u} \right]}{K_{iu}} \right)+ \frac{\left[ P \right]}{K_{dP}}+\frac{\left[ Q \right]}{K_{dQ}}+\frac{\left[ P \right]\left[ Q \right]}{K_{dP}K_{dQ}}\left( 1+\frac{\left[ I_{u} \right]}{K_{iu}} \right)}$$

$$v_{r}=\frac{\frac{k_{cat,r}\left[ E_{tot} \right]\left[ P \right]\left[ Q \right]}{K_{dP}K_{dQ}}}{1+\frac{\left[ A \right]}{K_{dA}}+\frac{\left[ B \right]}{K_{dB}}+\frac{\left[ A \right]\left[ B \right]}{K_{dA}K_{dB}}\left( 1+\frac{\left[ I_{u} \right]}{K_{iu}} \right)+ \frac{\left[ P \right]}{K_{dP}}+\frac{\left[ Q \right]}{K_{dQ}}+\frac{\left[ P \right]\left[ Q \right]}{K_{dP}K_{dQ}}\left( 1+\frac{\left[ I_{u} \right]}{K_{iu}} \right)}$$

Combining competitive and uncompetitive inhibition, we get the full rate forms:

$$v_{f}=\frac{\frac{k_{cat,f}\left[ E_{tot} \right]\left[ A \right]\left[ B \right]}{K_{dA}K_{dB}}}{1+\frac{\left[ A \right]}{K_{dA}}+\frac{\left[ B \right]}{K_{dB}}+\frac{\left[ A \right]\left[ B \right]}{K_{dA}K_{dB}}\left( 1+\frac{\left[ I_{u} \right]}{K_{iu}} \right)+ \frac{\left[ P \right]}{K_{dP}}+\frac{\left[ Q \right]}{K_{dQ}}+\frac{\left[ P \right]\left[ Q \right]}{K_{dP}K_{dQ}}\left( 1+\frac{\left[ I_{u} \right]}{K_{iu}} \right)+\frac{\left[ I_{c} \right]}{K_{Ic}}}$$

$$v_{r}=\frac{\frac{k_{cat,r}\left[ E_{tot} \right]\left[ P \right]\left[ Q \right]}{K_{dP}K_{dQ}}}{1+\frac{\left[ A \right]}{K_{dA}}+\frac{\left[ B \right]}{K_{dB}}+\frac{\left[ A \right]\left[ B \right]}{K_{dA}K_{dB}}\left( 1+\frac{\left[ I_{u} \right]}{K_{iu}} \right)+ \frac{\left[ P \right]}{K_{dP}}+\frac{\left[ Q \right]}{K_{dQ}}+\frac{\left[ P \right]\left[ Q \right]}{K_{dP}K_{dQ}}\left( 1+\frac{\left[ I_{u} \right]}{K_{iu}} \right)+\frac{\left[ I_{c} \right]}{K_{Ic}}}$$

Finally, if we allow for substrates with stoichiometry other than 1 of the form:

$$aA+bB{E \atop\begin{aligned} \rightleftharpoons\\ \end{aligned}}pP+qQ$$

and substitute

$$V_{max,f}=k_{cat,f}\left[ E_{tot} \right], V_{max,r}=k_{cat,r}[E_{tot}]$$

we can generalize the rate form to:

$$v_{f}=\frac{V_{max,f}\left( \frac{\left[ A \right]}{K_{dA}} \right)^{a}\left( \frac{\left[ B \right]}{K_{dB}} \right)^{b}}{1+a\frac{\left[ A \right]}{K_{dA}}+b\frac{\left[ B \right]}{K_{dB}}+\left( \frac{\left[ A \right]}{K_{dA}} \right)^{a}\left( \frac{\left[ B \right]}{K_{dB}} \right)^{b}\left( 1+\frac{\left[ I_{u} \right]}{K_{iu}} \right)+ p\frac{\left[ P \right]}{K_{dP}}+q\frac{\left[ Q \right]}{K_{dQ}}+\left( \frac{\left[ P \right]}{K_{dP}} \right)^{p}\left( \frac{\left[ Q \right]}{K_{dQ}} \right)^{q}\left( 1+\frac{\left[ I_{u} \right]}{K_{iu}} \right)+\frac{\left[ I_{c} \right]}{K_{Ic}}}$$

$$v_{r}=\frac{V_{max,r}\left( \frac{\left[ P \right]}{K_{dP}} \right)^{p}\left( \frac{\left[ Q \right]}{K_{dQ}} \right)^{q}}{1+a\frac{\left[ A \right]}{K_{dA}}+b\frac{\left[ B \right]}{K_{dB}}+\left( \frac{\left[ A \right]}{K_{dA}} \right)^{a}\left( \frac{\left[ B \right]}{K_{dB}} \right)^{b}\left( 1+\frac{\left[ I_{u} \right]}{K_{iu}} \right)+ p\frac{\left[ P \right]}{K_{dP}}+q\frac{\left[ Q \right]}{K_{dQ}}+\left( \frac{\left[ P \right]}{K_{dP}} \right)^{p}\left( \frac{\left[ Q \right]}{K_{dQ}} \right)^{q}\left( 1+\frac{\left[ I_{u} \right]}{K_{iu}} \right)+\frac{\left[ I_{c} \right]}{K_{Ic}}}$$

### S3: Derivation of Thermodynamic Constraints For Rate Law

For the general reaction:

$$aA+bB{E \atop\begin{aligned} \rightleftharpoons\\ \end{aligned}}pP+qQ$$

The equilibrium constant is defined as

$$K_{eq}=\frac{\left[ P_{eq} \right]^{p}\left[ Q_{eq} \right]^{q}}{\left[ A_{eq} \right]^{a}\left[ B_{eq} \right]^{b}}$$

Further, at equilibrium, the forward and reverse rates of reaction are equal:

$$v_{f}=v_{r}$$

Substituting the above Eq. <X.X>,

$$V_{max,f}\left( \frac{\left[ A \right]}{K_{dA}} \right)^{a}\left( \frac{\left[ B \right]}{K_{dB}} \right)^{b}=V_{max,r}\left( \frac{\left[ P \right]}{K_{dP}} \right)^{p}\left( \frac{\left[ Q \right]}{K_{dQ}} \right)^{q}$$

Further substituting Eq. <X.X>,

$$K_{eq}=\frac{V_{max,f}}{V_{max,r}}\frac{K_{dP}^{p}K_{dQ}^{q}}{K_{dA}^{a}K_{dB}^{b}}$$

Or more generally, for reaction *j*,

$$K_{eq,j}=\frac{V_{max,f}}{V_{max,r}}{\prod_{i\in\{substrates, products\}} K_{d_{j,i}}}^{S_{i,j}}$$

This equation, which is equivalent to the Haldane relationship for Michaelis-Menten kinetics, can be used to constrain parameters within a reaction.

All $K_{eq}$ values were calculated with eQuilibrator. Because eQuilibrator uses the Alberty method, which is based on group contribution and defines protons (H^+^) to have zero free energy (Alberty, 2003), protons did not have a sampled dissociation constant and were not included in the above thermodynamic relationship. Water molecules, however, had non-zero free energies in the Alberty method. Therefore, a dissociation constant for water was sampled in each reaction where water was a substrate, though this constant was only used for the above thermodynamic relationship and was not applied in the kinetic rate forms during ODE integration.

### S4: Logarithmic Transformation of Thermodynamic Constraints

To allow the use of the above relationship as a set of convex linear constraints, logarithmic transformation was applied. To apply these linear constraints in the form

$$\boldsymbol{Ax}\leq\boldsymbol{b}$$

as expected in MATLAB, where ***A*** is a matrix, ***x*** is the vector of log-transformed variables, and **b** is a vector of constraints, the following were constructed:

$$A_{rate constants}=\left[ \begin{matrix} 1 & -1 & 0 & 0 & \ldots\\ 0 & 0 & 1 & -1 & \ldots\\ 0 & 0 & 0 & 0 & \ldots\\ \vdots& \vdots& \vdots& \vdots& \ddots\end{matrix} \right]$$

where each row is a net reaction corresponding to a single $K_{eq}$ value and each column corresponds to the rate constants (twice the number of net reactions),

$$A_{dissociation constants}=S_{j,i_{j}}$$

where each row is a net reaction corresponding to a single $K_{eq}$ value and each column corresponds to the dissociation constants, and $S_{j,i_{j}}$is the stoichiometry in reaction *j* of the *i*th dissociation constant (not to be confused with the *i*th metabolite in the system, since a single metabolite may have multiple dissociation constants).

We then transform the inequality

$$\boldsymbol{Ax}\leq\boldsymbol{b, l}\boldsymbol{b}_{\boldsymbol{x}}\boldsymbol{\leq x\leq u}\boldsymbol{b}_{\boldsymbol{x}}$$

using slack constraints so that it becomes

$$\boldsymbol{Ax-s=0, l}\boldsymbol{b}_{\boldsymbol{s}}\boldsymbol{\leq s\leq u}\boldsymbol{b}_{\boldsymbol{s}}$$

Which is then

$$\left[ \begin{matrix} \boldsymbol{A} & \boldsymbol{-I} \end{matrix} \right]\left[ \begin{matrix} \boldsymbol{x} \\ \boldsymbol{s} \end{matrix} \right]\boldsymbol{=0=A'x'}$$

The final constraints are then given by:

$$\boldsymbol{A}^{\boldsymbol{'}}\boldsymbol{=}\left[ \begin{matrix} \boldsymbol{A}_{\boldsymbol{rate constants}} & \boldsymbol{A}_{\boldsymbol{dissociation constants}} & \boldsymbol{I}_{\boldsymbol{j}} \end{matrix} \right]$$

$$\boldsymbol{x'=}\left[ \begin{matrix} \ln\left( V_{max,f,1} \right) \\ \ln\left( V_{max,r,1} \right) \\ \ln\left( V_{max,f,2} \right) \\ \ln\left( V_{max,r,2} \right) \\ \vdots\\ \ln\left( V_{max,f,j} \right) \\ \ln\left( V_{max,r,j} \right) \\ \vdots\\ \ln\left( K_{dX_{1},1} \right) \\ \ln\left( K_{dX_{2},1} \right) \\ \ln\left( K_{dX_{1},2} \right) \\ \ln\left( K_{dX_{2},2} \right) \\ \vdots\\ \ln\left( K_{dX_{i},j} \right) \\ \vdots\\ \ln\left( K_{eq,1} \right) \\ \ln\left( K_{eq,1} \right) \\ \vdots\\ \ln\left( K_{eq,j} \right) \end{matrix} \right]$$

### S5a: Metabolic Control Analysis

The features of metabolic control analysis relevant to understanding the response variable of reaction fluxes were concentration elasticities ($\epsilon_{ji}^{x}$), which calculates the local response of flux in each elementary (forward and reverse) reaction with respect to the control variable of metabolite concentrations within the same reaction, or flux control coefficients ($C_{lj}^{J}$), which calculates the global response of flux with respect to all other model fluxes. To analyze the response variable of metabolite concentrations, we used the Jacobian ($J_{ii^{'}}$), which calculates the global response of each concentration with respect to the control variables of all other metabolite concentrations, and concentration control coefficients ($C_{ij}^{x}$), which calculates the global response of concentrations with respect to all model fluxes. In terms of metabolite concentrations $x_{i}$ and $x_{i^{'}}$ and reaction fluxes $v_{j}$ and $v_{j^{'}}$, these quantities are defined in Equations <X.X> below:

$$\epsilon_{ji}^{x}=\frac{x_{i}}{v_{j}}\frac{\partial v_{j}}{\partial x_{i}}$$

$$C_{ij}^{x}=\frac{v_{j}}{x_{i}}\frac{\partial x_{i}}{\partial v_{j}}$$

$$C_{jj^{'}}^{v}=\frac{v_{j^{'}}}{v_{j}} \frac{\partial v_{j}}{\partial v_{j^{'}}}$$

$$J_{ii^{'}}=\frac{x_{i^{'}}}{x_{i}}\frac{\partial}{\partial x_{i^{'}}}\left( \frac{\partial x_{i}}{\partial t} \right)$$

Instead of numerically calculating each term according to its definition, an analytical method of calculation was used. Because quantities in metabolic control analysis are not able to be calculated in non-rank systems (i.e., those where not all metabolites are linearly independent of all others), metabolic conservation analysis was applied as described in (Greene et al., 2017), and so the reduced stoichiometric matrix $\boldsymbol{S}_{\boldsymbol{R}}$ replaces the full stoichiometric matrix $\boldsymbol{S}$**,** where $\boldsymbol{S}_{\boldsymbol{R}}$ contains only those metabolite species which are linearly independent, such that $\boldsymbol{S}_{\boldsymbol{R}}$ is full rank and the full stoichiometric matrix $\boldsymbol{S}$ can be described by $\boldsymbol{S}_{\boldsymbol{R}}$ and the link matrix $\boldsymbol{L}$ by the equation

$$\boldsymbol{S=L}\boldsymbol{S}_{\boldsymbol{R}}$$

First, the concentration elasticity was calculated from each reaction rate equation using the MATLAB Symbolic Math Toolbox (MATLAB 2020b, The MathWorks, Inc.). From this elasticity term, all other terms were calculated, as described in (Hofmeyr, 2001). These calculations are given below:

$$\boldsymbol{J=}\boldsymbol{S}_{\boldsymbol{R}}\boldsymbol{\epsilon}^{\boldsymbol{x}}\boldsymbol{L}$$

and $\boldsymbol{\epsilon}^{\boldsymbol{x}}$ is the matrix form of the reduced (independent species only) elasticities.

The concentration control coefficient matrix is then calculated as

$$\boldsymbol{C}^{\boldsymbol{x}}\boldsymbol{=-L}\boldsymbol{J}^{\boldsymbol{-1}}\boldsymbol{S}_{\boldsymbol{R}}$$

and the flux control coefficient matrix is calculated as

$$\boldsymbol{C}^{\boldsymbol{v}}\boldsymbol{=}\boldsymbol{\epsilon}^{\boldsymbol{x}}\boldsymbol{C}^{\boldsymbol{x}}\boldsymbol{+}\boldsymbol{I}_{\boldsymbol{j}}$$

where $\boldsymbol{I}_{\boldsymbol{j}}$ is the identity matrix with a row for each elementary reaction.

### S5b: MCA for Analysis of Conserved Parameter values

While the goal of this analysis was to determine if the values of individual parameters were conserved between parameter sets, it was not feasible to perform this analysis using the absolute values various parameters, as these values cannot be interpreted without the context of the concentration of their corresponding substrates and the concentrations and dissociation/inhibition constants of all other species in that reaction. Instead, we analyzed the concentration elasticity corresponding to each dissociation constant parameter, as this value takes this information into account to convert each parameter into a single value that is more representative of the sensitivity of a reaction rate with respect to its substrates. For example, if we desired to measure the conservation of the value of the dissociation constant *K_dA_* for metabolite *A* in reaction *A + B 🡪 P + Q*, the effect of the parameter *K_dA_* on the reaction rate is highly dependent not only on its own value but also on the concentration of *A, B, P,* and *Q,* as well as the parameters *K_dB_, K_dP_, and K_dQ_*; instead, the concentration elasticity of *A* takes these values into account implicitly.

Because there are far more species and parameters in the model than there are measured species, and because many of the model reactions are far from measured species, it was expected that many parameters would have low sensitivity, meaning that the concentration elasticities for these parameters could span the full range of values (from 0 to 1) and still adequately fit the model. Conversely, we expected that some other parameters would have high sensitivity, which would be necessary to provide enough control over metabolism to fit the observed data. The central question of this analysis was whether the sensitive parameters would be conserved across all the top models in the final ensemble, or if different models would have different sensitive parameters. Whether or not a parameter value (or elasticity value by proxy) is conserved in its value across models can also be thought of as an analysis on how “constrained” that parameter might be.

### S6: Parameter Sensitivity Analysis

Parameter sensitivity analysis was calculated numerically using the finite differences method. Each parameter $\theta_{i}$ was adjusted individually, and no corrections were made for other parameters, as it was assumed that the small change (1%) in parameter value would not move the adjusted $K_{eq}$ value out of the constrained range. The SSE fitness function was then recalculated as:

$$SSE_{adj \theta_{i}}=f\left( \theta_{1},\ldots,\theta_{i}+\Delta\theta_{i},\ldots,\theta_{k} \right)$$

where $f$ is the function to calculate the sum of squared errors, and $\Delta\theta_{i}$ is 1% of the initial value of $\theta_{i}$. The sensitivity for $\theta_{i}$ was then calculated as

$$s_{\theta_{i}}=\frac{SSE_{adj \theta_{i}}-SSE_{base}}{\Delta\theta_{i}}=\frac{f\left( \theta_{1},\ldots,\theta_{i}+\Delta\theta_{i},\ldots,\theta_{k} \right)-f\left( \theta_{1},\ldots,\theta_{i},\ldots,\theta_{k} \right)}{\Delta\theta_{i}}$$

This process was repeated for each parameter in a parameter set, and then again for all parameter set in the screened ensemble.

### S7: Software Settings

#### Sampling Settings

All parameters were sampled from a log-uniform distribution within one order of magnitude (0.1x to 10x) of the pre-assigned prior value. If no value was assigned for a dissociation constant, the value of 1.0 mM was used, and if no value was assigned for an inhibition constant, the value of 0.5 mM was used (pre-assigned values were required for rate constants).

The unitless Gibbs Free Energy at physiological conditions ($\Delta G^{'m}/ RT)$ for each reaction were gathered using the Equilibrator 3.0 python package, which supplied mean values as well as errors. For each parameter set, the utilized value of $\Delta G^{'m}/ RT$ was sampled from within a uniform distribution created by adding and subtracting the Equilibrator error from the mean. The equilibrium constant was then calculated using the relationship:

$$K_{eq}=e^{-\frac{\Delta G^{'m}}{RT}}$$

Once all dissociation constants and the equilibrium constant were sampled, the Haldane relationship was used to determine if the quantity of $V_{max,f}/V_{max,r}$ was greater or smaller than 1, and the larger of these terms was sampled from within its log-uniform distribution. The last term was then calculated directly.

#### Matlab Settings

All calculations were performed in MATLAB 2020b.

To allow repeatability in both serial and parallel parameter set generation, a random number seed corresponding to the index of the parameter set being generated was used, along with the MATLAB “twister” algorithm. For example, to generate parameter set *i*, we would initialize the random number generator with*:*

Rng(i, ‘twister’)

ODE integrations were performed with the MATLAB *ode15s* ODE solver. We used a default relative tolerance (RelTol) of 1e-30, and a custom event function which stopped any integrations requiring more than 5 seconds. A pre-written Jacobian function was passed into *ode15s*.

#### Northwestern QUEST HPCC Settings

The patternsearch algorithm ran in parallel with a maximum walltime of 4 hours on the Northwestern Quest HPCC. Optimization runs for each initial point used 250 CPUs across several computing nodes.

### S8: Fitness Function and Metabolite Weightings

The fitness function used in this work was the root mean squared error (RMSE) which was weighted by metabolite identities. This fitness for each parameter set is calculated as:

$$RMSE=\sum_{c} \sum_{i} \sum_{t} \sqrt{{w_{i}\left( \bar{x}_{c,i,t}^{pred}-x_{c,i,t}^{exp} \right)}^{2}}$$

where $x_{c,i,t}$ is the predicted or experimental metabolite concentration of metabolite *i* (glucose, succinate, acetate, ethanol, lactate, butanol, pyruvate, malate, and citrate) in condition *c* (butanol positive and butanol negative) at time *t* where experimental measurements were taken ($t=\left[ 0, 0.5, 1, 1.5, 2, 2.5, 3, 4, 5, 6, 7, 8, 8, 8, 20, 20, 20 \right]$ or $t=\left[ 0, 0.5, 1, 1.5, 2, 2.5, 3, 4, 5, 6, 7, 8, 8, 8, 24, 24, 24 \right]$). Replicate measurements at $t=8$ hours and $t=20$ or $t=24$ hours were treated as separate measurements and were not averaged before use in the fitness function. The utilized weights were $w_{butanol}=20$ and $w_{succinate}=6$, with all other species having a weight of 1.

### S9: Calculation of Metabolite Contribution from Component Fluxes

In many cases in this work, it was useful to attribute the change over time in the concentration of a metabolite to the reactions whose fluxes caused this change. To distinguish between multiple reactions which might either produce or consume a metabolite, we first used the known vector of metabolite concentrations, along with the known reaction rate laws, to calculate the instantaneous reaction fluxes $v_{net}$ at each time point in the ODE integration. Then, instead of performing the matrix multiplication

$$\boldsymbol{Sv=}\frac{d\boldsymbol{x}}{dt}$$

to calculate the change in each metabolite concentration from *all* fluxes, we calculated the change in each concentration from each single column of $\boldsymbol{S}$, corresponding to individual reactions *j*, by multiplying by the matrix of zeros with $\boldsymbol{v}$ on the diagonal:

$$\boldsymbol{S*diag}\left( \boldsymbol{v} \right)\boldsymbol{=}\left[ \begin{matrix} \frac{d\boldsymbol{x}_{\boldsymbol{1}}}{dt} & \frac{d\boldsymbol{x}_{\boldsymbol{2}}}{dt} & \boldsymbol{\ldots} & \frac{d\boldsymbol{x}_{\boldsymbol{j}}}{dt} \end{matrix} \right]$$

Each column $\frac{d\boldsymbol{x}_{\boldsymbol{j}}}{dt}$ now gives the individual contribution of reaction *j* to the change in each metabolite, and can be used for further analysis.

### S10: Paramater Sensitivity to Unmeasured Behaviors

In all the above analyses, the source of constraints on parameter is understood to be the selection and optimization of parameters to the experimental data; that is, the reason a given parameter tends toward a specific value across all models in the ensemble is because that value best allows the models to agree with data. However, we wanted to explore if parameter values were ever also tightly constrained with respect to their influence on unobserved behavior, or more generally, behavior that did not directly impact the fitness function used in optimization. To test this hypothesis, we performed an “alternative fitness” parameter sensitivity analysis, wherein the effect of small changes in each parameter value was measured not in the change of that model’s fit to experimental data, but in the change in some other behavior or metric. Because we had seen that redox cofactors were critical in controlling metabolic behavior and in connecting the behavior of disparate pathways, we chose this metric to be the cumulative rate of NADH oxidation to NAD^+^. Because the goal of this analysis was to understand which parameters were predominantly controlled by the metabolic redox balance, or potentially controlling this balance, this rate of NADH oxidation was calculated using only the reactions NAD transhydrogenase and NAD dehydrogenase and not other reactions including these cofactors.

The results of this analysis are summarized In Supplementary Fig. S12<X>. The single most influential parameter, as measured by the median behavior across the top 20 parameter sets, was the rate constant for NAD dehydrogenase. As with the analysis of elasticities, this result shows that these reactions exhibit a large amount of their own control over the redox balance, as opposed to simply being controlled by external changes in the redox balance. The dissociation constants for NADH, ubiquinol-8, and NAD^+^ in NAD dehydrogenase are also the three most sensitive dissociation constant parameters, though there is a range of predictions between models as to specific values of sensitivity. While this shows that NAD dehydrogenase maintains predominantly internal control over the redox balance, several other parameters also have strong sensitivity with respect to NADH oxidation rate. In particular, the rate constants and dissociation constants for PTA, responsible for acetate production, are among the most sensitive parameters. This is somewhat interesting, as the acetate pathway from acetyl-CoA does not itself use any redox equivalents; instead, this further demonstrates the interconnectedness of metabolism and the need for fine-tuning during parameter optimization to capture these subtle effects. Additionally, the rate constants for ACALD (ethanol production), and PDH/PFL (acetyl-CoA formation), which do explicitly use redox equivalents, show very high sensitivity for NADH oxidation rate. Other parameters of interest are the rate constants for thiolase as well as its dissociation constants for acetyl-CoA and CoA; the conserved sensitivity of the dissociation constant for CoA, in particular, is interesting because it is in the reverse (but favorable) direction with respect to butanol production, emphasizing that even fitting the production of butanol – the primary goal of the model – is heavily reliant on carefully optimized parameter values which have effects on many model behaviors beyond the experimental data. Lastly, the NADH oxidation rate was highly sensitive to the parameters governing malic enzymes (ME1 and ME2) and malate dehydrogenase. While it is not surprising that these parameters, whose reactions produce redox equivalents, would be sensitive to this metric, it is somewhat unexpected that these anaplerotic reactions consistently produce enough flux to impact redox balance, as the large majority of carbon in succinate production comes from glutamate rather than anaplerosis and TCA flux. Nonetheless, after optimization, nearly all models in the final ensemble predicted that these reactions carry significant flux, and so these reactions have impacts both in redox balance and in further connecting different pathways of the metabolic network.

### S11: Partial Differential Equation Modeling of Gas Transfer and Quasi-Equilibrated Kinetic Modeling

Cell-free experiments were conducted in a 1.5 mL tube, and reaction volume totaled approximately 30 µL. Therefore, we wanted to calculate the extent to which gasses were equilibrating between the aqueous phase in the reactions and the gaseous phase in the tube headspace, and whether diffusion limitations would establish a gradient in the aqueous phase. To test these possibilities, a partial differential equation (PDE) was simulated. The two gas species in the model were O_2_ and CO_2_. Initial concentrations in the gas phase were set by atmospheric conditions (Keeling, 2013). Henry’s law constants were used to calculate the initial concentration of aqueous gas species at the surface (z = 0) at 25°C, and initial aqueous gas species at the bottom of the tube (z = 4 mm) was assumed to be zero). Rates of aqueous gas consumption or production were set based on the maximum and minimum rate constants for reactions producing or consuming each species in the Varner model (Horvath et al., 2020). Diffusion is assumed to be fast in the gas phase, and for this initial study, the gas-phase concentration was assumed to be constant. Diffusion constants for each species in water were taken from (Tse & Sandall, 1979). The final PDE for each gas in the aqueous phase was as follows:

$$\frac{d\left[ gas_{aq} \right]}{dt}=D_{gas}\frac{d^{2}\left[ gas_{aq} \right]}{dx^{2}}-V_{max,cons}*\left[ gas_{aq} \right]+V_{max,prod}$$

Constants:

$D_{CO_{2}}={1.6*10}^{-3}\frac{mm^{2}}{s}$ (CRC Handbook, 2007)

$D_{O_{2}}={2.4*10}^{-3}\frac{mm^{2}}{s}$ (CRC Handbook, 2007)

$H_{CO_{2}}^{cc}=8.3*{10}^{-1}$ (Sander, 2015)

$H_{O_{2}}^{cc}=3.2*{10}^{-2}$ (Sander, 2015)

Consumption and Production Rates (Horvath et al., 2020):

$$V_{cons,{CO}_{2}}=9.1*{10}^{-2} s^{-1}$$

$$V_{prod,CO_{2}}=1.7*{10}^{-3}\frac{mM}{s}$$

$$V_{cons,O_{2}}=5.6*{10}^{-2} s^{-1}$$

$$V_{prod,O_{2}}=0 (no model reactions produced O_{2})$$

Atmospheric CO_2_ and equilibrium aqueous concentration:

$$CO_{2,atm}=400 ppm$$

$$\left[ CO_{2,gas} \right]=ppm*\frac{n}{V}=ppm*\frac{P}{RT}=\frac{400}{{10}^{6}}*\frac{1 atm}{\left( 8.206*{10}^{-2}\frac{L*atm}{K*mol} \right)\left( 298 K \right)}=1.6*{10}^{-2} mM$$

$$\left[ CO_{2,aq} \right]=H_{CO_{2}}^{cc}\left[ CO_{2,gas} \right]=1.4*{10}^{-2} mM$$

Atmospheric O_2_ and equilibrium aqueous concentration:

$$O_{2,atm}=2.1*10^5 ppm$$

$$\left[ O_{2,gas} \right]=ppm*\frac{n}{V}=ppm*\frac{P}{RT}=\frac{2.1*{10}^{5}}{{10}^{6}}*\frac{1 atm}{\left( 8.206*{10}^{-2}\frac{L*atm}{K*mol} \right)\left( 298 K \right)}=8.6 mM$$

$$\left[ CO_{2,aq} \right]=H_{CO_{2}}^{cc}\left[ CO_{2,gas} \right]=2.7*{10}^{-1} mM$$

PDE initial conditions:

$$\left[ gas_{aq} \right]\left( t=0 \right)=0 mM$$

PDE boundary conditions:

$$\left[ CO_{2,aq} \right]\left( x=0, t>0 \right)=1.4*{10}^{-2} mM$$

$$\left[ O_{2,aq} \right]\left( x=0, t>0 \right)=2.7*{10}^{-1} mM$$

$$\frac{d\left[ gas_{aq} \right]}{dt}\left( x=4 mm, t>0 \right)=0$$

The results of the PDE (Supplementary Fig. S<X>) show that for the wide range of consumption and production rates tested, there was a negligible gradient in concentration of each aqueous gas species in the *z-*direction. Therefore, we made the further assumption that there was no gradient, and gasses would be instantly equilibrated between the aqueous and gas phases.

To model this equilibration, we chose to implement a set of fast kinetic reactions. This was done instead of an explicit equilibrium implemented as a set of differential algebraic equations (DAE) as it was seen that the kinetic implementation ran significantly faster using the MATLAB *ode15s* ODE solver.

The relative ratio of gas exchange kinetics was set by the respective Henry’s Law constants. For example, for carbon dioxide:

$$C{O_{2}}_{g}\begin{matrix} k_{f} \\ \rightleftarrows\\ k_{r} \end{matrix}C{O_{2}}_{aq}$$

Each “kinetic” rate is assumed to be mass action:

$$v_{f}=k_{f}\left[ C{O_{2}}_{g} \right], v_{r}=k_{r}\left[ C{O_{2}}_{aq} \right]$$

Because the gas exchange is assumed to be fast compared to enzymatic reactions and is therefore at equilibrium,

$$v_{f}=v_{r}, k_{r}\left[ C{O_{2}}_{aq} \right]=k_{f}\left[ C{O_{2}}_{g} \right]$$

And therefore the Henry’s Law constant $H_{CO_{2}}^{cc}$constrains the kinetic parameters:

$$H_{CO_{2}}^{cc}=\frac{\left[ C{O_{2}}_{aq} \right]}{\left[ C{O_{2}}_{g} \right]}=\frac{k_{f}}{k_{r}}$$

To ensure that gas exchange is faster than enzymatic reactions, the smaller of $k_{f}$ and $k_{r}$ was set to an arbitrarily high value, which in this work was 10^10^ hr^-1^.
